# Up-regulation of cholesterol synthesis pathways and limited neurodegeneration in a knock-in *Sod1* mutant mouse model of ALS

**DOI:** 10.1101/2023.05.05.539444

**Authors:** Janice A. Dominov, Laura A. Madigan, Joshua P. Whitt, Katerina L. Rademacher, Kristin M. Webster, Hesheng Zhang, Haruhiko Banno, Siqi Tang, Yifan Zhang, Nicholas Wightman, Emma M. Shychuck, John Page, Alexandra Weiss, Karen Kelly, Alper Kucukural, Michael H. Brodsky, Alexander Jaworski, Justin R. Fallon, Diane Lipscombe, Robert H. Brown

## Abstract

Amyotrophic lateral sclerosis (ALS) is a severe neurodegenerative disorder affecting brain and spinal cord motor neurons. Mutations in the copper/zinc superoxide dismutase gene (*SOD1*) are associated with ∼20% of inherited and 1-2% of sporadic ALS cases. Much has been learned from mice expressing transgenic copies of mutant SOD1, which typically involve high-level transgene expression, thereby differing from ALS patients expressing one mutant gene copy. To generate a model that more closely represents patient gene expression, we created a knock-in point mutation (G85R, a human ALS-causing mutation) in the endogenous mouse *Sod1* gene, leading to mutant SOD1^G85R^ protein expression. Heterozygous *Sod1^G85R^* mutant mice resemble wild type, whereas homozygous mutants have reduced body weight and lifespan, a mild neurodegenerative phenotype, and express very low mutant SOD1 protein levels with no detectable SOD1 activity. Homozygous mutants exhibit partial neuromuscular junction denervation at 3-4 months of age. Spinal cord motor neuron transcriptome analyses of homozygous *Sod1^G85R^* mice revealed up-regulation of cholesterol synthesis pathway genes compared to wild type. Transcriptome and phenotypic features of these mice are similar to *Sod1* knock-out mice, suggesting the *Sod1^G85R^* phenotype is largely driven by loss of SOD1 function. By contrast, cholesterol synthesis genes are down-regulated in severely affected human *TgSOD1^G93A^* transgenic mice at 4 months. Our analyses implicate dysregulation of cholesterol or related lipid pathway genes in ALS pathogenesis. The *Sod1^G85R^* knock-in mouse is a useful ALS model to examine the importance of SOD1 activity in control of cholesterol homeostasis and motor neuron survival.

**SIGNIFICANCE STATEMENT:** Amyotrophic lateral sclerosis is a devastating disease involving the progressive loss of motor neurons and motor function for which there is currently no cure. Understanding biological mechanisms leading to motor neuron death is critical for developing new treatments. Using a new knock-in mutant mouse model carrying a *Sod1* mutation that causes ALS in patients, and in the mouse, causes a limited neurodegenerative phenotype similar to *Sod1* loss-of-function, we show that cholesterol synthesis pathway genes are up-regulated in mutant motor neurons, whereas the same genes are down-regulated in transgenic *SOD1* mice with a severe phenotype. Our data implicate dysregulation of cholesterol or other related lipid genes in ALS pathogenesis and provide new insights that could contribute to strategies for disease intervention.

## INTRODUCTION

Amyotrophic lateral sclerosis (ALS) is a lethal disorder leading to motor neuron (MN) degeneration in brain and spinal cord. About 10% of ALS cases are familial; approximately 20% of those are caused by mutations in the gene encoding copper/zinc cytosolic superoxide dismutase (SOD1) (Deng et al., 1993; Rosen et al., 1993; Cleveland and Rothstein, 2001; Pasinelli and Brown, 2006; Rothstein, 2009). Typically, a single mutant *SOD1* gene copy is sufficient to cause ALS, suggesting mutant gain-of-function pathology. While mechanisms underlying mutant SOD1 neurotoxicity have not been fully delineated, mutant SOD1 proteins disrupt many molecular pathways (Kiskinis et al., 2014; Kaur et al., 2016; Xu et al., 2022). Current therapeutic approaches target pathogenic gain-of-function *SOD1* mutations. However, recent observations suggest that SOD1 loss-of-function variants in humans can trigger an infantile progressive motor neurological syndrome (Andersen et al., 2019; Park et al., 2019; Ezer et al., 2022). Therefore, both SOD1 loss and gain-of-function mutations are implicated in pathogenesis (Saccon et al., 2013).

Transgenic mutant SOD1 protein expression causes severe motor neuropathology (Gurney et al., 1994; Dal Canto and Gurney, 1995; Ripps et al., 1995; Wong et al., 1995; Bruijn et al., 1997), typically in a dosage-dependent manner (Dal Canto and Gurney, 1995; Alexander et al., 2004). Transgenic wild-type (WT) SOD1 over-expression causes slowly progressive, subtle pathology (Dal Canto and Gurney, 1995; Tu et al., 1996; Jaarsma et al., 2000). Moreover, very high expression of WT SOD1 in mice homozygous for the WT SOD1 transgene (expressing 50-fold more than the endogenous levels, and 4-fold more than in transgenic G93A mice) causes early lethality (Graffmo et al., 2013). Some transgenic mutants do not develop phenotypes (e.g., transgenic *SOD1^A4V^* mice (Gurney et al., 1994)) unless they also express high levels of human WT SOD1 (Deng et al., 2006). These data suggest that an ALS phenotype can develop when SOD1 levels rise above a threshold.

SOD1 knock-out mice exhibit a slowly progressive distal motor axonopathy but without loss of MNs (Flood et al., 1999; Shefner et al., 1999; Muller et al., 2006; Fischer et al., 2012), and are more sensitive to neuronal injury compared to WT (Reaume et al., 1996). This observation is consistent with the view that SOD1 insufficiency contributes to ALS neuropathology in patients with SOD1 mutations that eliminate or suppress dismutation activity.

Mutant mouse *Sod1* knock-in alleles controlled by endogenous transcription elements eliminate confounding effects of high-level transgene overexpression, thereby more closely modeling mutant gene expression in human disease. To date, however, knock-in mutations in the *SOD1* gene do not induce severe pathology in mice. A spontaneous mutation in a highly conserved region of mouse *Sod1* (*Sod1^E77K^*, which is not ALS-associated) causes slightly reduced SOD1 activity but has no associated phenotype (Luche et al., 1997). A different ENU-induced ALS-associated point mutation, *Sod1^D83G^*, reduces SOD1 protein expression and SOD1 activity in homozygous mice. *Sod1^D83G/D83G^* mice have some associated MN loss but no paralysis, and phenotypic features resemble *Sod1* knock-outs, including distal axonopathy and liver tumors (Joyce et al., 2015).

We generated a new *Sod1* knock-in mouse, carrying the *Sod1^G85R^* mutation that causes human ALS, to test the hypothesis that *Sod1* knock-in models elicit ALS-related phenotypes and, as such, can identify ALS-associated pathogenetic gene pathways important for modeling disease. *Sod1^G85R/+^* heterozygotes are not distinguishable from WT, but *Sod1^G85R/G85R^* homozygous mice had very low SOD1 protein levels, no SOD1 activity, and phenotypic features similar to SOD-deficient mice including low body weight, reduced survival time, and partial muscle denervation. We compare electrical properties and nerve-muscle junction integrity, muscle fiber size, and MN transcriptomes of WT and *Sod1^G85R/G85R^* mice. We find genes involved in cholesterol synthesis are upregulated in MNs in *Sod1^G85R/G85R^* and *Sod1^-/-^* mice; perhaps contributing to MN survival. By contrast, cholesterol pathway genes are down-regulated in MNs of severely-affected transgenic *TgSOD1^G93/+^* mice compared to WT and *Sod1^G85R/G85R^*. Our studies provide new insights into MN pathophysiology associated with the *Sod1^G85R^* mutation.

## MATERIALS AND METHODS

### *Sod1* mutant mouse production

All procedures involving mice were approved by the University of Massachusetts Chan Medical School (UMass Chan) or Brown University Animal Care and Use Committees and followed guidelines of the US Public Health Service Policy on Humane Care and Use of Laboratory Animals and the USDA Animal Welfare Act.

*Sod1* knock-in mutant mice designed to emulate human *SOD1* pathogenic mutations were produced in the UMass Chan Transgenic Animal Modeling Core using CRISPR/SpCas9 mutagenesis reagents generated by the UMCMS Mutagenesis Core. Two types of knock-in mice were used in these studies: a *Sod1^G85R^* and a *Sod1^-/-^* line, *Sod1^insAA/insAA^*, which arose from mutagenesis targeting the beginning of *Sod1* exon1 intended to create *Sod^A4V^* mutant mice as part of another study. To create mutants, C57BL6 mouse single-cell embryos were injected with a mixture containing SpCas9 mRNA, guide RNAs (gRNAs) targeting specific regions of the *Sod1* gene (designed using the Bioconductor software package CRISPRseek (Zhu et al., 2014)), and single stranded oligonucleotide donor DNA (120-122 nt, IDT) containing single base changes that lead to the desired amino acid substitutions when introduced by homologous recombination (Extended Table 1-1). The donor DNA used in the mutagenesis that created the *Sod1^insAA^* (and the intended *Sod^A4V^*) mouse was not actually incorporated into the *Sod1^insAA^* locus by homologous recombination. Pups were screened for the desired mutations by PCR amplification and sequencing of ear snip DNA using primers listed in Extended Table 1-2. For these PCRs, we used Phire Hot Start II DNA Polymerase (Thermo Scientific), 98°C 90 seconds (sec), 30 cycles of 98°C 10 sec, 65°C 15 sec, 72°C 30 sec, then 72°C 10 minutes (min). Two founder lines were identified that carry the *Sod1^G85R^* mutation, Line 3, and Line 20. All experiments were done using Line 20 mice except for long term survival studies, which involved approximately equal numbers of both lines of mice with no discernable differences among them.

Mutant mice were backcrossed to C57BL6 (Jackson Laboratory) and interbred to generate desired genotypes. Allele-specific primers (Extended Table 1-2) that distinguish mutant and WT pups were used in PCR reactions to genotype subsequent progeny. For the *Sod1^G85R^* mutation, we used One Taq Hot Start Polymerase (New England Biolabs), 94°C 30 sec, then 30 cycles of 94°C 20 sec, 65°C 20 sec, 68°C 60 sec, then 68°C 5 min. PCR products were analyzed on 2% agarose TAE gels. For the *Sod1^insAA^* mutation we used Phire Hot Start II DNA Polymerase, 98°C 90 sec, 30 cycles of 98°C 10 sec, 71°C 15 sec, 72°C 30 sec, then 72°C 10 min.

We additionally used the transgenic mouse strain *TgSod1^G93A/+^* (C57BL6-based, B6.Cg-Tg(SOD1*G93A)1Gur/J, 004435, Jackson Laboratory), which develops severe ALS symptoms due to human mutant transgene overexpression and has a 50% survival time of 157 days and average lifespan of 171 days.

### Physiological assessment of forelimb and hindlimb grip strength

Various aspects of phenotypic characterization of *Sod1^G85R^* mice were conducted in three separate laboratories at Brown University and the UMass Chan Medical School as described in figure legends. Forelimb and hindlimb muscle strength was assessed by a grip strength behavior task. Mice were first weighed and then either scruffed for hindlimb analysis or lifted by the tail for forelimb analysis, enticing the hindlimbs or forelimbs to grasp the pull-bar assembly connected to the grip strength meter (Columbus Instruments). The mouse was drawn along a straight line leading away from the sensor until the grip was broken and the peak amount of force in kilograms per unit force (KGF) was recorded. This was repeated 5 times and the maximum value recorded was used for analysis. The experimenter was blind to the genotype of the mice during the experiment and analysis. Values that were greater than two standard deviations from the mean for each age/genotype group were removed from the analysis. Measurements were performed in three different laboratories and data compiled as described in the figure legends. Laboratory A tested forelimb and hindlimb grip strength on 4-, 12-, and 36-week-old male and female WT, *Sod1^G85R/+^*, and *Sod1^G85/G85R^* mice and Laboratory B performed forelimb grip strength analysis on a cohort of male WT, *Sod1^G85R/+^*, and *Sod1^G85R/G85R^* mice at approximately 1, 3, 6, and 9 months of age. Additional grip strength measurements of forelimb and all four limbs were similarly done in Laboratory C on older mice ranging from 4 to 32 months of age.

### Rotarod Performance

Motor coordination and balance of 6-month-old male WT, *Sod1^G85R/+^*, and *Sod1^G85R/G85R^* mice were measured using an accelerating rotarod (Ugo Basile, Model 7650, Varese, Italy). The mice were habituated to the rotarod wheel spinning at four revolutions per minute (RPM) for 90 seconds. During formal testing, mice were subjected to gradual rotarod acceleration from 4 to 40 RPM. Latency to fall was automatically recorded by the system. The mean value of 5 trials for each mouse was calculated. Individual mouse recordings were excluded if the mouse fell from the rod while moving backwards, slipped, or jumped at slow speed. All testing was conducted by an experimenter who was blind to genotype.

### Open Field Movement

Open field analysis was performed on a cohort of male WT, *Sod1^G85R/+^,* and *Sod1^G85R/G85R^* mice at 1, 3, 6, and 9 months of age to assess general locomotor activity and anxiety phenotypes. Mice were placed in a custom-made white, plastic arena (50X50X40cm) that was illuminated (∼10Lux) in a dimly lit room. Mice were acclimated to the testing room at least 15 min prior to testing and then placed in the center of the open field arena. Trace path was recorded and analyzed with Noldus EthoVision 10.1 software to determine the distance traveled by each mouse during the 7 min session using a video-tracking system.

### Evans blue dye labeling of motor neurons

MNs projecting to tibialis anterior (TA) and soleus muscles were visualized for slice physiology and histology by retrograde labelling. Mice were anesthetized with 3% isoflurane, and legs were shaved and cleaned prior to injection. Either TA or soleus muscles were surgically exposed and injected with 30-40 μl of 2% (w/v) Evans blue dye solution (Sigma Aldrich) in phosphate buffered saline (PBS) over 3 min time period using a 30-gauge needle. Surgical incisions were sutured, and the animal was allowed to recover for 16 to 36 hours prior to use.

### Spinal cord slice electrophysiology

Transverse slices of lumbar spinal cord of male and female *Sod1^G85R/G85R^* homozygotes and WT littermates were used for electrophysiological recordings. Mice were anesthetized with isoflurane and placed in a prone position. After confirming the lack of hind limb pinch reflexes, transcardiac perfusion was performed with 50 ml of 35°C dissection solution containing (in mM): 93 NMDG, 2.5 KCl, 1.2 NaH2PO4, 30 NaHCO3, 20 HEPES, 2 Thiourea, 5 sodium ascorbate, 3 myo-inositol, 10 N-Acetyl-L-cysteine, 25 dextrose, 5 ethyl pyruvate, 8 MgSO4, 0.5 CaCl2, aerated with 95% O2 and 5% CO_2_ to a final pH of 7.4 (final osmolarity, 297–307 mOsm). The spinal cord was removed *en bloc* and submerged in 4°C dissection solution prior to embedding in a 4% (w/v) low melt agarose solution cooled to 37°C. The agarose-embedded preparation was cooled at room temperature for 1 min then glued to a custom made vibratome stage. 350 μm transverse slices of lumbar cord were cut (Leica VT1200S) in 4°C dissection solution. Each spinal cord slice was transferred to 30% (w/v) polyethylene glycol-2000 for 1 min, washed twice for 1 min in 35°C dissection solution, and transferred to an incubation chamber for 30 min in the same solution. After recovery, slices were incubated in 35°C holding solution for at least 30 min containing (in mM): 93 NaCl, 2.5 KCl, 1.2 NaH_2_PO4, 30 NaHCO_3_, 20 HEPES, 2 Thiourea, 5 sodium ascorbate, 3 myo-inositol, 10 N-Acetyl-L-cysteine, 25 dextrose, 5 ethyl pyruvate, 8 MgSO_4_, 0.5 CaCl_2_, aerated with 95% O_2_ and 5% CO_2_ to a final pH of 7.4 (final osmolarity, 297–307 mOsm). For electrophysiology, spinal cord slices were transferred to a submersion style recording chamber containing (in mM): 125 NaCl, 3 KCl, 1.2 NaH_2_PO_4_, 25 NaHCO_3_, 15 dextrose, 1 MgSO_4_, 1.2 CaCl_2_, aerated with 95% O_2_ and 5% CO_2_ to a final pH of 7.4 (final osmolarity, 297–307 mOsm) exchanged at ∼3 ml/min. Large Evans blue-positive MNs were identified in the ventral horn of the spinal cord using epifluorescence. Whole cell patch recordings were performed using DIC microscopy at room temperature. Recording pipette resistances were 3 to 7 MΩ filled with (in mM): 130 KCl, 5 NaCl, 2 MgCl_2_, 10 HEPES, 4 MgATP, 0.3 NaGTP, 10 Na phosphocreatine, and 0.4% (w/v) biocytin adjusted to pH 7.3 with KOH and osmolarity of 300 mOsm. Recordings were performed in current clamp mode, low-pass filtered at 10 kHz, and digitized at 50 kHz. Data were acquired and analyzed using PClamp software (Axon Instruments). Data were compared across genotypes, muscle groups, and age using multivariate ANOVA analyses (IBM SPSS Statistics). The experimenter was blind to the genotype of the mice during the experiment and data analysis.

### Visualization and analysis of neuromuscular innervation

Animals received an intraperitoneal injection with 0.05 ml of pentobarbital euthanasia solution and were perfused with 20 ml of 1 mM ethylenediaminetetraacetic acid (EDTA) in PBS, followed by 15 ml 4% PFA in PBS. TA muscles were harvested and immersion-fixed in 4% PFA in PBS for 1 h, followed by three 20 min washes in PBS. Muscles were stored in 0.01% sodium azide in PBS at 4°C until sectioned.

All immunohistochemistry steps were performed at room temperature unless otherwise noted. TA muscles were embedded in 3% agarose in PBS and sectioned on a vibratome (Leica) at 100 μm or 200 μm. Sections were incubated in 100 mM glycine in PBS for 1 hour (h) followed by a PBS rinse. Sections were then blocked for 4 h in 2% bovine serum albumin (BSA), 4% goat serum, and 0.5% Triton X-100 in PBS, incubated overnight at 4°C with primary antibodies diluted in blocking buffer, and then washed 3 x 20 min in 0.1% Triton X-100 in PBS. After a 4-h incubation with secondary antibodies and fluorescent α-bungarotoxin diluted in blocking buffer, sections were washed 3 x 20 min in 0.1% Triton X-100 in PBS, then mounted under coverslips with Fluoromount G (Invitrogen) and stored at 4°C until imaging. Images were acquired on an Olympus FV3000 confocal microscope.

Primary antibodies used were a rabbit polyclonal antibody against vesicular acetylcholine transporter (VAChT, 1:200, Synaptic Systems) and a rabbit polyclonal antibody against class III β-tubulin (TUBB3, 1:1000, BioLegend). Secondary antibodies used were Alexa647-conjugated donkey anti-rabbit (1:200, Invitrogen) and Alexa647-conjugated goat anti-rabbit (1:200, Invitrogen). Alexa488-conjugated α-bungarotoxin (1:1000, ThermoFisher) was added with secondary antibodies.

For each animal, a minimum of 800 NMJs were analyzed. We confirmed that analyzing this number of synapses produces results that are not significantly different from fully analyzing the over 2000 NMJs per muscle (data not shown). ImageJ was used to view individual muscle sections in 3D image stacks, and analysis was performed blinded to the identity of the animal that the section originated from. NMJs were located based on postsynaptic staining before presynaptic coverage was assessed. NMJs were categorized as having 100%, between 50% and 100%, between 0% and 50%, or 0% presynaptic coverage of the postsynaptic site. After unblinding, NMJs in each category across all analyzed sections for a single animal were expressed as a percentage of all analyzed synapses. These values were averaged across all animals of the same age, sex, and genotype and expressed as mean ± SEM with N listed in figure legends. Pairwise comparisons between groups were performed using unpaired, two-tailed t-tests (GraphPad Prism), with *p* values indicated in the figures. Multiple comparisons were performed using one-way ANOVA followed by Holm’s post hoc test (α = 0.05; KaleidaGraph) with *p* values indicated in the figures.

### Quantification of retraction bulbs

For 3-month-old males, all sections that were analyzed for neuromuscular innervation were also used to count retraction bulbs, blinded to the identity of the animal. Retraction bulbs were defined as thickened structures at the ends of axons (at least twice the diameter of the axon shaft) that were not in direct contact with a postsynaptic site. After unblinding, the number of retraction bulbs across all analyzed sections for a single animal were expressed as a percentage of all postsynaptic sites in these sections. These values were averaged across all animals of the same genotype and expressed as mean values ± SEM with N listed in figure legends. Pairwise comparisons between groups were performed using unpaired, two-tailed t-tests (GraphPad Prism), with *p* values indicated in the figures.

### Compound Muscle Action Potential (CMAP)

A data acquisition interface (National Instruments USB-6343), a preamplifier (WPI DAM50), a stimulus isolator (WPI A365D), and electrodes (28-gauge TECA model 902-DMF25-S) were used for CMAP acquisition. Detailed procedures were previously described (Sakamoto et al., 2009). Briefly, isoflurane anesthesia was maintained at 1.5-2.5%. Ground electrode was inserted subcutaneously into the middle of the upper back. Recording electrode was subcutaneously inserted ∼1 mm into the belly of the TA muscle. The reference electrode was inserted subcutaneously near the ankle, between two tendons; the stimulation electrode was placed at 10 mm lateral to the midline, 5 mm above the base of the tail; and the return electrode was placed approximately 2-4 mm lateral to the midline, 5 mm above the base of the tail. At least 3 CMAP waves were acquired. The highest peak-to-peak CMAP amplitude was selected. The mean of both sides of the highest CMAP amplitude was used for statistical analyses. Data from individual mice were collected longitudinally from 30 to 180 days of age, except for 2 WT mice which were only measured at 30 days. Statistical analyses were performed with IBM SPSS Statistics, version 26 (IBM Japan) and applying Student’s t-test to assess differences between genotypes. *p* values are reported in figure legends.

### Myofiber size

TA muscles were dissected and frozen in freezing isopentane. Ten μm cryosections from the mid-belly region of the muscles were fixed with 4% PFA in PBS for 10 min, treated for 15 min with 0.1% Triton X-100 in PBS, washed 3 times with PBS, blocked with 5% BSA and 2% goat serum in PBS for 1 h at room temperature, and then incubated overnight at 4°C with 1:000 rabbit anti-laminin (Sigma-Aldrich L9393). Sections were washed 3 times with PBS and then incubated with 1:1000 goat anti-rabbit IgG Alexa-Fluor 488 (Invitrogen, #A-11034) in PBS for 1 h at room temperature and mounted in Vectashield Antifade Mounting Medium with DAPI (Vector Laboratories H-1002-10 and H-1000-10). Images used for analysis were captured using a Nikon E800 fluorescent microscope with a 10x objective. Images were processed using MyoVision (Wen et al., 2018) to determine minimum Feret diameter measurements for each myofiber (Briguet et al., 2004), 650-1800 fibers examined per mouse. Output images were manually examined, and any misidentified fibers were excluded. Representative images were acquired with a Zeiss LSM 800 Confocal Laser Scanning Microscope with a 20x objective. The average myofiber minimum Feret diameter for each mouse was used for statistical analysis using an unpaired t-test in GraphPad Prism 8. Approximately 900 fibers were analyzed per mouse. The experimenter was blinded to genotype for the analysis.

### Tissue collection for protein and RNA analyses

Spinal cord and brain tissue was collected from *Sod1* mutant and WT mice at 30, 70 and 120 days of age (postnatal P30, P70, P120). Mice were euthanized by isoflurane inhalation and bilateral thoracotomy then transcardially perfused with ice cold PBS for 1-2 min to remove blood from tissues. Spinal cord tissue was extruded using cold PBS and dissected into cervical, thoracic, and lumbar segments which were rapidly frozen by suspension in liquid N2 vapor. Brain tissue was kept on ice and dissected to separate cortex, cerebellum, brainstem, and “other” portions then rapidly frozen on dry ice. Tissues were stored at -80°C.

### Protein Analyses

Thoracic spinal cord and brain (“other”-not cortex, brainstem, cerebellum) tissues from 120-day-old mutant and WT mice were split in order to analyze a portion of each tissue for SOD1 protein expression on western blots and another portion of each for SOD1 activity. For western blots, tissues were lysed in RIPA buffer (40 mM Tris–HCl pH 8, 150 mM sodium chloride 1% Triton X--100, 0.5% sodium deoxycholate, 0.5% SDS, protease inhibitors (cOmplete, Roche)). Protein concentration was determined using a Pierce BCA assay (ThermoFisher). Protein extracts were mixed 1:1 with 2x Laemmli sample buffer containing B-mercaptoethanol, incubated at 100°C for 5 min, separated on 12% Tris-Glycine SDS-PAGE acrylamide gels (ThermoFisher) then blotted onto nitrocellulose using an iBlot 2 system (ThermoFisher). Blots were incubated in LI-COR Odyssey Blocking Buffer (BB) for 1 h at room temperature then in primary antibodies diluted in BB overnight at 4°C. Blots were rinsed with PBS, 0.1% Tween 20 buffer (PBST), incubated in secondary antibodies in BB for 1 h at room temperature, PBST and PBS rinsed, then imaged using a LI-COR Odyssey infrared imager. LI-COR Image Studio Lite software was used to quantitate protein expression from scanned images. Antibodies used were rabbit anti-SOD1 (Genetex, GTX100554) 1/1000, goat anti-β-actin (Abcam, ab8229) 1/3000, 680RD donkey anti-rabbit (LICOR, 926-68073) 1/5000, 800CW donkey anti-goat (LICOR, 926-32214) 1/5000.

SOD1 activity gels were processed as described (Beauchamp and Fridovich, 1971). Tissues were lysed in 0.1% NP-40 in 150mM NaCl with protease inhibitors (Roche) and protein concentration determined as above for western blots. Protein extracts, along with several dilutions of a SOD1 protein standard (Sigma, S9697) were separated on native 12% Tris-Glycine acrylamide gels (ThermoFisher) at 4°C then incubated in 2.45 mM nitro blue tetrazolium for 20 min followed by a solution containing 28mM TEMED, 28 µM riboflavin, and 36 mM potassium phosphate pH 7.8 for 15 min. The gel was illuminated on a white light box for 5-10 min, covered by a foil lined box then imaged using a GelDoc Imaging System (Bio-Rad).

### Laser capture microdissection and RNA expression analysis

Laser capture microdissection (LCM) was used to isolate MNs from the lumbar region of spinal cords, predominantly L4/L5. For this, 7 μm cryosections of lumbar cords (all from male mice) were fixed immediately in 75% ethanol and with stained using an Arcturus Histogene Frozen Section Staining Kit (ThermoFisher) following manufacturers procedures with one exception: An additional step was added after the first water rinse in which RNAlater-ICE Frozen Tissue Transition Solution (ThermoFisher) was added to preserve the RNA prior to adding the Histogene staining solution. MNs were identified as large (∼30+ μm diameter), blue-stained cells in the ventral horn of cord and were isolated using an Arcutus^XT^ Laser Capture Microdissection System. Approximately 600 - 800 MNs cells were collected per mouse. RNA was extracted using an PicoPure RNA Isolation Kit (ThermoFisher), which included a DNAse digestions step, and analyzed for integrity on an Advanced Analytical Fragment Analyzer (UMass Chan Molecular Biology Core). All but two samples (RIN 6.7, 6.8) had RIN values 7.0 or greater, (mean ± SD; 7.6 ± 0.3). We used Novogene Ultra Low RNA Input RNA sequencing services to produce cDNA libraries and perform the sequencing. For each sample, 2 ng RNA was reverse transcribed to create a library (Smart-Seq v4 Ultra Low Input RNA / cDNA library) which was sequenced to generate 50 million read pairs (150 bp paired reads) (Illumina NovaSeq 6000 PE150). A total of 42 RNA samples processed by Novogene in two separate RNA-seq library batches were used for differential RNA expression analysis (Extended Table 7-1). These included *Sod1^G85R/G85R^* at P120 (n = 4), P70 (n = 4), P30 (n = 4); *Sod1^-/-^* at P120 (n = 4), P70 (n=4), P30 days (n = 4); transgenic SOD1*^G93A/+^* (*TgSOD1^G93A/+^*) at P120 (n = 4), and WT at P120 days (n = 6), P70 (n = 5), P30 (n = 3). We used the DolphinNext RNA-seq Pipeline (Yukselen et al., 2020) to map reads to the mouse RefSeq genome using RSEM and STAR. Differential mRNA expression among samples was evaluated using normalized read counts in DEBrowser (Kucukural et al., 2019) using a Combat batch correction module to adjust for batch variation among samples at each age (batch correction samples: P120 n=20; P70 n=13, P30 n=11). We analyzed samples grouped by age (P120, P70 or P30) to identify genes differentially expressed in *Sod1* mutants compared to age matched WT mice and also between mutant mouse types. Differential expression was based on a cut-off adjusted p value (padj) < 0.05 for all pairwise comparisons. Pathway analyses were then performed to determine the biological features reflected buy the gene expression changes in the mutant mice at each age. For this we used the STRING database portal for protein-protein interaction networks and functional enrichment analysis (Szklarczyk et al., 2021), Gene Ontology (GO) analyses (Ashburner et al., 2000; Gene Ontology, 2021) and the KEGG Pathway Database for functional pathway analysis (Kanehisa and Goto, 2000; Kanehisa et al., 2021). In addition, we interrogated the expression of specific sets of genes associated with specific functional GO terms in the *Sod1* mutant and WT LCM MN samples to determine differences in gene expression specifically in these pathways These included a set of 78 genes associated with cholesterol synthesis (GO:0006695) and related pathways (Mazein et al., 2013; Hartmann et al., 2021) and a broad, pooled set of 877 ion channel, transporter and related genes (GO:0005326 neurotransmitter transmembrane transporter activity; GO:0015075 ion transmembrane transporter activity; GO:0022803 passive transmembrane transporter activity). Gene lists are in the first sheets of Extended Tables 9-1 or 11-1.

### Ventral and dorsal spinal cord tissue collection and RNA analysis

To further evaluate the expression of various genes in the spinal cord, additional cryosections were collected from the same lumbar spinal cord tissues as used for the LCM analyses (L4 - L6 regions). Cryosections (30 μm) were collected and processed the same way as done for LCM section staining except omitting the Histogene blue dye stain step. A scalpel blade was used to scrape the entire ventral portion of the sections into a tube containing 200 μl TRIzol. The dorsal portion of the section was then placed into a separate TRIzol tube. The central ∼1 mm strip of each section containing the central canal was left on the slide and not collected. A total of ∼ 60 x 30 μm sections (∼1.8 mm tissue depth) were collected per cord. RNA was purified from these ventral and dorsal scrapes using a Zymo Direct-zol RNA Microprep Kit, including a DNAse step. RNAs (150 ng) were reverse transcribed using an Applied Biosystems High-Capacity cDNA Reverse Transcription Kit (ThermoFisher) with Applied Biosystems RNAse inhibitor. All samples were processed at the same time and diluted to allow multiple PCR assays to be performed on the same cDNA preps, preventing any cDNA batch variability. Quantitative PCR (QPCR) was performed in a Bio-Rad CFX384 real-time PCR machine using 2X DyNAmo HotStart SYBR Green qPCR Kit reagents and methods (ThermoFisher) and gene transcript-specific primer pairs (Extended Table 12-1), triplet reactions per condition. ANOVA analyses were performed using GraphPad Prism software.

### Experimental Design and Statistical Analyses

Experiments were designed and evaluated using statistical methods as described in various sections above. Kaplan-Meier analysis (GraphPad Prism) was used to determine mutant and WT animal survival. ANOVA or Student’s t-test analyses (GraphPad Prism), as noted in figures, was used to evaluate mouse body weight, grip strength and rotarod performance.

## RESULTS

### *Sod1* mutant mouse model

We used CRISPR mutagenesis to introduce mutations into the endogenous *Sod1 mouse* gene. The resultant knock-in strains expressed mutant transcripts under the control of the endogenous *Sod1* promoter (Figure 1A). *Sod1^G85R^* mice contain a point mutation (transcript NM_011434.2, c.256G>C) equivalent to a homologous ALS-causing human mutation that leads to a glycine to arginine substitution at amino acid 85 of the mature SOD1 protein. We also created a *Sod1*-knock-out strain, *Sod1^-/-^* through a 2-base pair insertion (insAA) that creates a premature stop codon and terminates translation in exon 1 of the *Sod1* transcript (Figure 1A). Homozygous *Sod1^G85R/G85R^* mice express very low levels of SOD1 protein in spinal cord and brain; no SOD1 protein is detectable in homozygous *Sod1 ^-/-^* mice (Figure 1B). There is no detectable SOD1 dismutase activity in either *Sod1^G85R/G85R^* or *Sod1^-/-^* tissue (Figure 1C).

**Figure 1.**
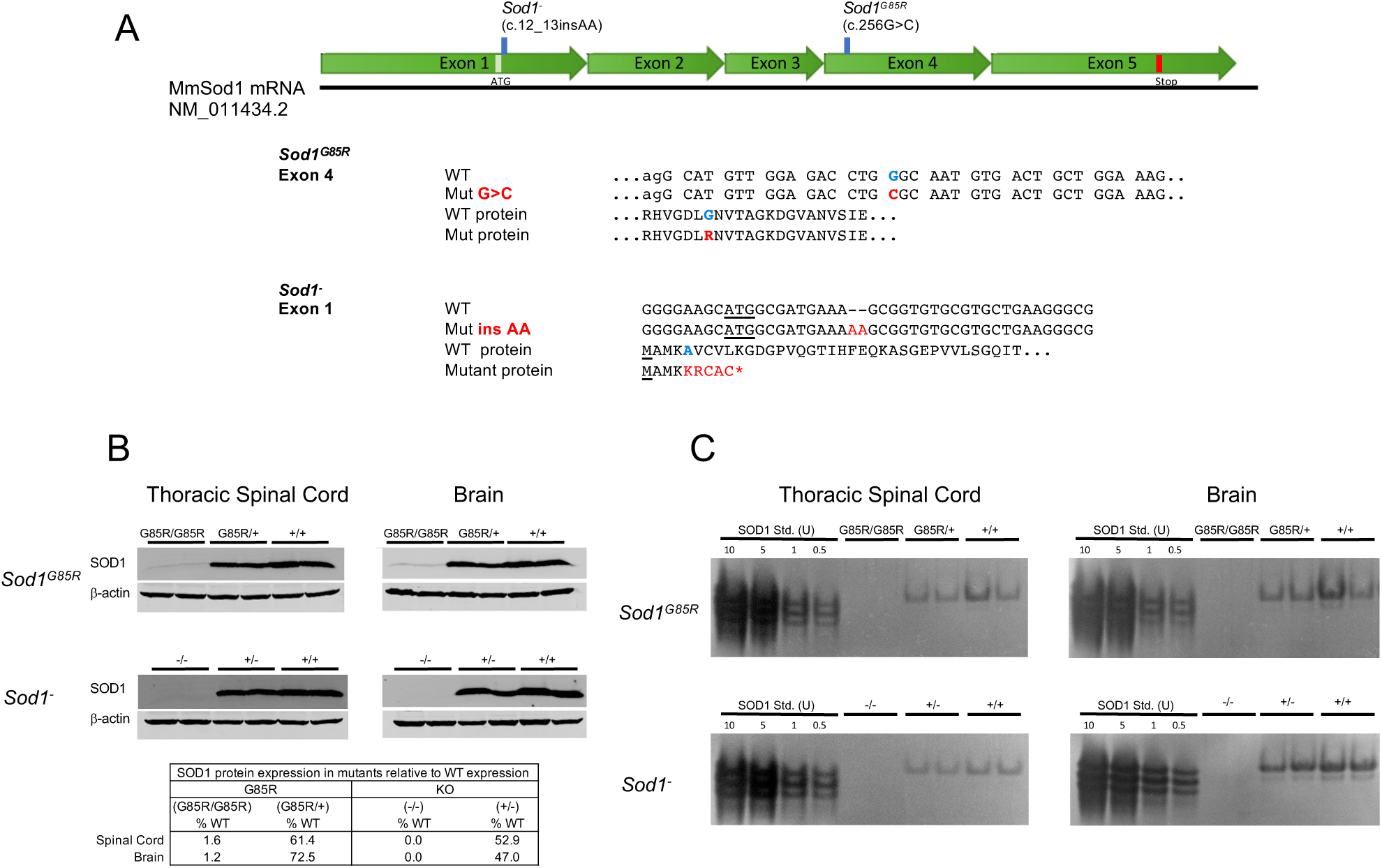
Generation of *Sod1^G85R^* knock-in and *Sod1^-^* knock-out mutant mice. **(A)** Mouse knock-in *Sod1^G85R^* and *Sod1^-^* (*Sod1^insAA^*) mutations located on a schematic diagram of *Sod1* mRNA (*top*). WT and mutated mRNA (exon 4 and exon 1) and protein sequences are shown. Intron sequences (lowercase), initial methionine sequences (underlined), and stop codon (*) are annotated. **(B)** Western blots of SOD1 protein together with β-actin control (probed on different regions of the same blots) for WT, *Sod1^G85R/G85R^* and *Sod1^-/-^* spinal cord and brain tissues, in duplicate.**(C)** SOD1 activity gels show no detectable SOD1 activity in either *Sod1^G85R/G85R^* or *Sod1^-/-^* tissues.

### *Sod1^G85R/G85R^* mice are smaller and have shorter lifespans and behavioral deficits

*Sod1^G85R^* mice were born at normal Mendelian frequencies (Figure 2A) and there was no gender bias (Extended Figure 2-1). *Sod1^G85R/+^* heterozygous mice were normal at weaning but *Sod1^G85R/G85R^* mice had significantly reduced body weight compared with WT or *Sod1^G85R/+^*(Figure 2C). Homozygous *Sod1^G85R/G85R^* mice had a significantly reduced lifespan compared to WT and *Sod1^G85R/+^* (Figure 2B). The median lifespan of *Sod1^G85R/G85R^* was ∼140-190 days shorter than WT and *Sod1^G85R/-^* for both sexes. The shorter lifespan of *Sod1^G85R/G85R^* mice was not due to severe loss of motor function or a neurodegenerative ALS-like phenotype, but rather to health issues associated with aging mice including tumors, skin lesions, rectal prolapse, eye problems and overall declining health requiring euthanasia. Liver or other tumors or abdominal masses were detected at death with comparable frequencies in WT, *Sod1^G85R/+^*, and *Sod1^G85R/G85R^* mice ∼37%, 21%, 39%, respectively). We also found brain aneurysms in several aged WT and *Sod1^G85R/+^* mice. Overall, tumors or aneurysms were not more frequent in older *Sod1^G85R/G85R^* mice as compared to *Sod1^G85R/+^* or WT mice.

**Figure 2.**
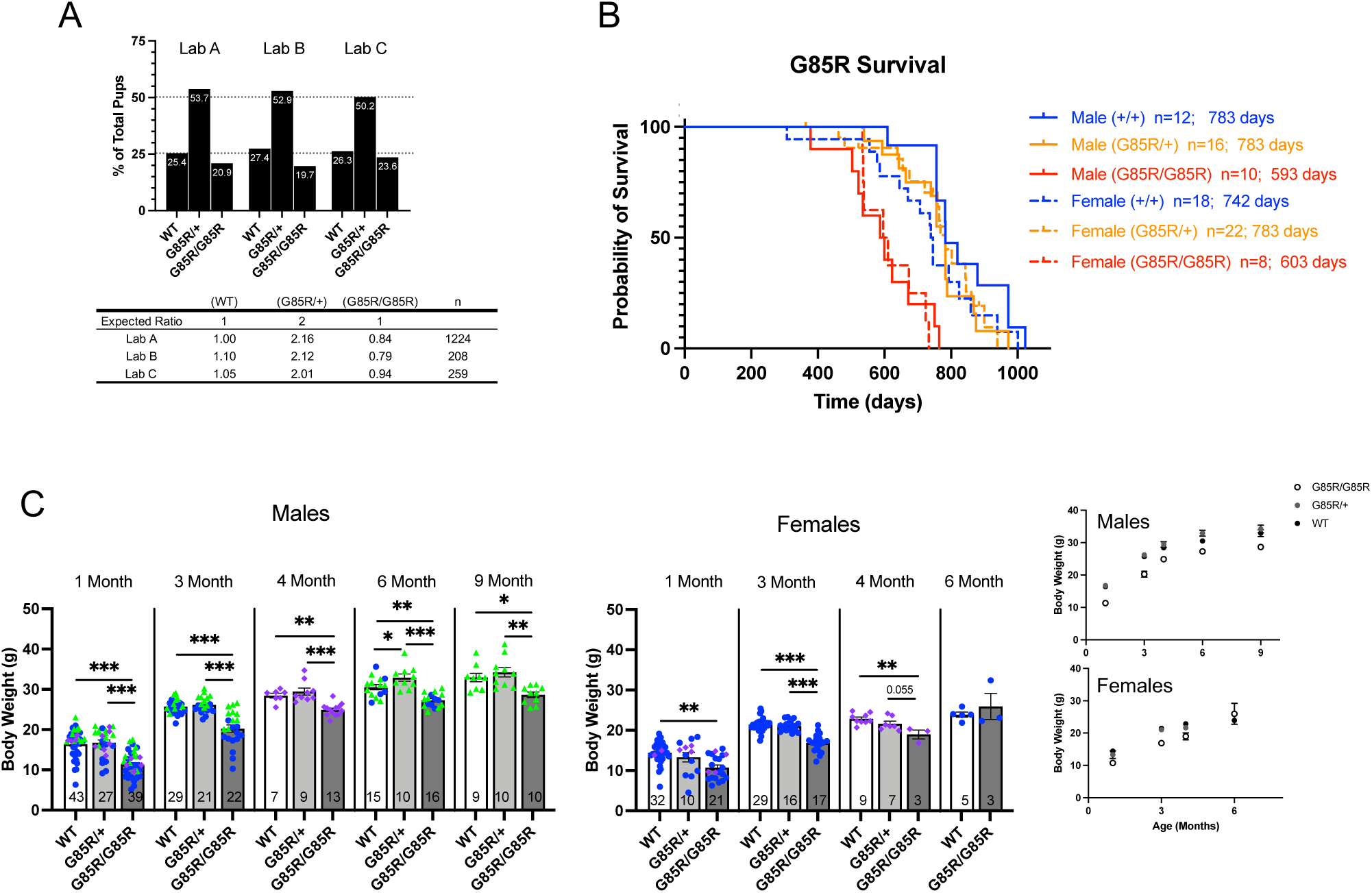
*Sod1^G85R/G85R^* mice are born at a normal frequency but exhibit a somewhat shortened lifespan and reduced body weight. **(A)** Genotype frequencies of wild-type (WT, *Sod1^+/+^*), *Sod1^G85R/+^* and *Sod1^G85R/G85R^* mutant mice compiled in each of the three laboratories conducting these studies show the expected 1:2:1 ratios. **(B)** Homozygous *Sod1^G85R/G85R^* mice have a reduced lifespan compared with WT and *Sod1^G85R/+^* siblings. The median survival time is listed with each genotype (days). Average male *Sod1^G85R/G85R^* lifespan is 189 days shorter than WT or *Sod1^G85R/+^* mice (p ≤ 0.0003); average female *Sod1^G85R/G85R^* lifespan is 138 and 179 days shorter than WT and *Sod1^G85R/+^* mice, respectively (p ≤ 0.0033). **(C)** *Sod1^G85R/G85R^* mice have reduce body weight compared with siblings. Data shown is compiled from the three laboratories conducting these studies with individual mice shown in blue (Lab A), green (Lab B) or purple (Lab C) symbols. Numbers in the bars = total n for each (Mean ± SEM; ANOVA *p < 0.05, **p < 0.005, ***p < 0.0001). The panels at the right show the same mean weight data plotted as a function of time.

While the growth and lifespan of *Sod1^G85R/G85R^* mice were not normal, they did not develop severe neuropathological changes or paralysis as reported in transgenic mutant mice expressing slightly elevated levels of human *SOD1^G85R^* (paralysis at ∼ 8 months) (Bruijn et al., 1997) or high levels of transgenic mouse *Sod1^G86R^* protein (paralysis at ∼ 4-5 months) (Ripps et al., 1995; Audet et al., 2010). Limb paralysis was only observed in two of 86 mice across our aged cohorts, both female, one WT (578 days old) and one *Sod1^G85R/+^* (675 days old). We evaluated the front (Figure 3A) and rear (Figure 3B) paw grip strength in cohorts of *Sod1^G85R^* mutant and WT mice at different ages. Raw grip strength measurements indicated that homozygous *Sod1^G85R/G85R^* males had significantly reduced front and rear paw grip strength compared to WT mice at 1, 3 and 6 months of age, and reduced front grip strength compared to *Sod1^G85R/+^* mice at 1 and 3 months (Figures 3A and 3B, top panels; rear grip strength for *Sod1^G85R/+^* mice was not measured). Similarly, *Sod1^G85R/G85R^* females exhibited lower front grip strength at 3 months, and lower rear grip strength at 1 and 3 months compared to WT or *Sod1^G85R/+^* cohorts. *Sod1^G85R/G85R^* mice have reduced body weight compared to WT mice and when grip strengths were normalized to body weight there was greatly reduced or no significant difference in front or rear grip strength between *Sod1^G85R/G85R^*, WT and *Sod1^G85R/+^* mice (Figures 3A and 3B, bottom panels). The reduced grip strength phenotype of *Sod1^G85R/G85R^* mice appears to be fully accounted for by reduced body mass.

**Figure 3.**
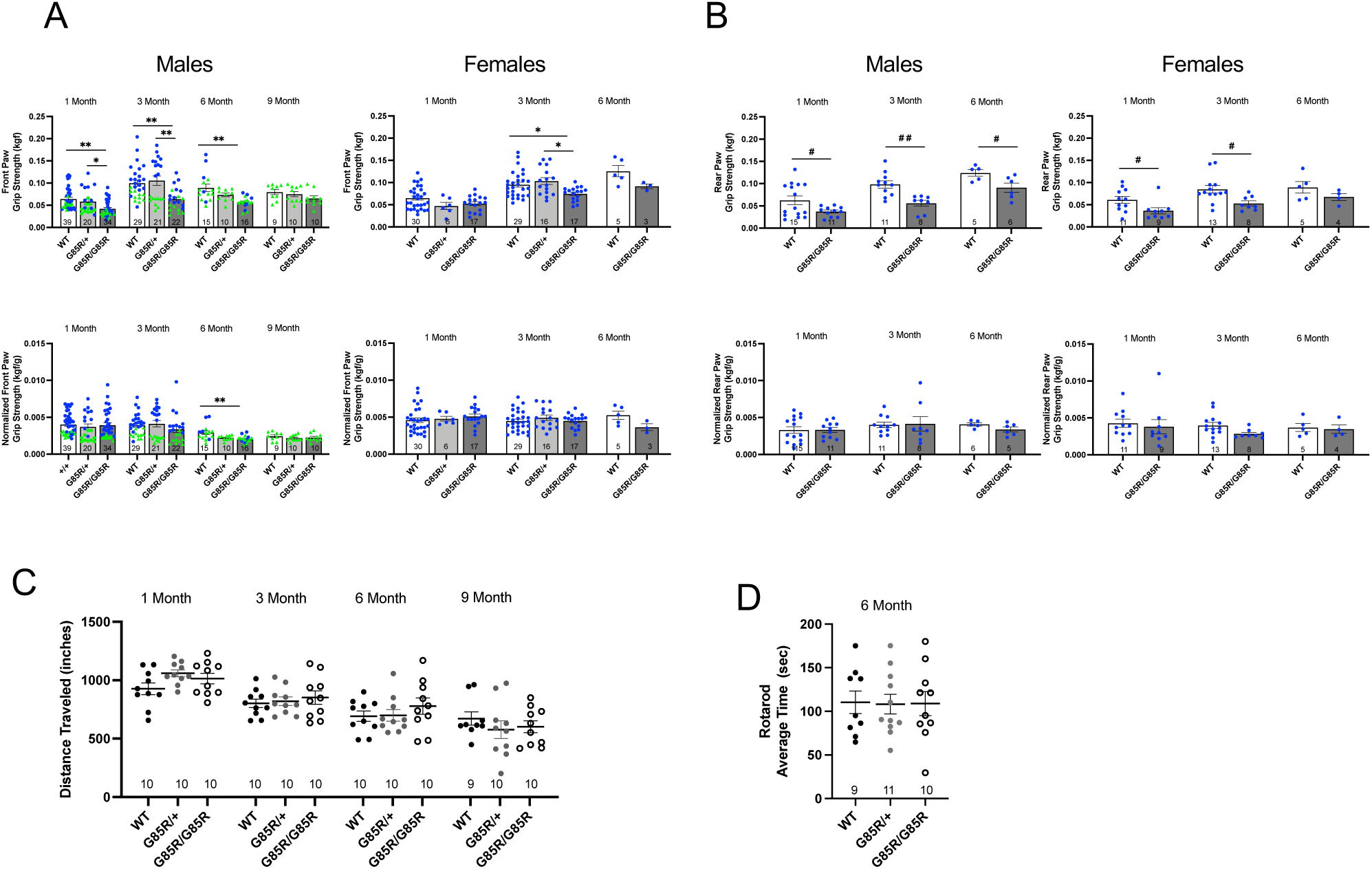
Mutant *SOD1^G85R/G85R^* mice have reduced grip strength compared to WT but similar overall motor activities. Raw front **(A)** and rear **(B)** paw grip strength data (top panels) compiled in two labs (blue symbols, Lab A; green symbols Lab B, numbers = total n) indicate that *Sod1^G85R/G85R^* mice have a reduced grip strength from one month of age on. When normalized for body weight (lower panels) there is no consistent difference in grip strength among WT and *Sod1^G85R/G85R^* mice (Mean ± SEM; ANOVA *p < 0.05, **p < 0.005; t-test #p < 0.05, ##p < 0.005). **(C)** Open field and **(D)** rotarod analyses show that overall mouse movement and latency to fall are similar between *Sod1^G85R/G85R^* and WT mice. For all panels, numbers at the bottom = n for each genotype.

We extended our observations of body weight, and grip strength in the cohorts of mice in our long-term survival study. The reduced body weight of *Sod1^G85R/G85R^* mice was a prominent phenotypic feature throughout the two-year period studied (Extended Figure 2-2). Front and four paw grip strengths of *Sod1^G85R/G85R^* mice were reduced compared to WT or *Sod1^G85R/+^* (Extended Figures 3-1 and 3-2) but, as with younger mice, this was accounted for by reduced body weight.

The overall movement of WT, *Sod1^G85R/+^*, or *Sod1^G85R/G85R^* mice at 1, 3, 6 or 9 months of age was not different across genotypes in open field tests based on distance traveled (Figure 3C). Similarly, WT, *Sod1^G85R/+^*, or *Sod1^G85R/G85R^* mice at 6 months of age did not differ in rotarod performance (Figure 3D). There were, however, subtle behavioral abnormalities evident in *Sod1^G85R/G85R^* mice from about 70 days of age onward. At 30 days, *Sod1^G85R/G85R^* mice suspended by their tails were indistinguishable from WT, but at 70 days they frequently exhibited tremors or intermittent spastic twitching behavior that became more evident at 120 days. This abnormal twitching persisted throughout the lifespan of the mice. Tail-suspended *Sod1^G85R/G85R^* mice also appeared less able to use axial muscles to twist laterally or turn backward toward their tails, a common behavior in normal mice. These observations suggested that motor deficits developed in *Sod1^G85R/G85R^* mice between 30 days and 70 days of age, but these did not progress to a severe loss of muscle function. We next evaluated MN function and the integrity of neuromuscular junctions in *Sod1^G85R/G85R^* mice.

### MNs in *Sod1^G85R/G85R^* mice are electrophysiologically indistinguishable from WT but their numbers are reduced

There is evidence that altered MN hyperexcitability precedes neuronal cell death and clinical symptoms in ALS (Vucic et al., 2008; Wainger et al., 2014; Ragagnin et al., 2019). We therefore examined the electrophysiological properties of MNs in the ventral horn of the spinal cord of *Sod1^G85R/G85R^* and WT littermates that project to either TA or soleus muscles. We studied mice at 1 and 3 months of age (Figure 4, Extended Table 4-1). Neurons projecting to either TA or soleus muscles were identified by retrolabeling using Evans blue dye injected into either muscle 1-3 days before recording. We measured multiple parameters including resting membrane potential, input resistance, rheobase, firing rate in response to current injection, and more. *Sod1^G85R/G85R^* and WT neurons were not functionally distinguishable across all parameters measured using multifactor ANOVA analyses. We were able to detect differences in the average threshold of current injection needed to trigger an action potential (rheobase), which was significantly higher in neurons projecting to the TA compared to those projecting to the soleus (p < 0.001, rheobase, Figure 4D), but this feature did not vary by age or genotype. The neurons sampled were consistent in size (capacitance, Figure 4A), input resistance (Figure 4B), and resting membrane potential (Figure 4C). *Sod1^G85R/G85R^* and WT neurons exhibited similar action potential firing rates (Figure 4E) and other physiological characteristics (Figure 4F-I). We only recorded from Evans blue labelled neurons and therefore our analyses were likely enriched in neurons capable of taking up the dye in muscle.

**Figure 4.**
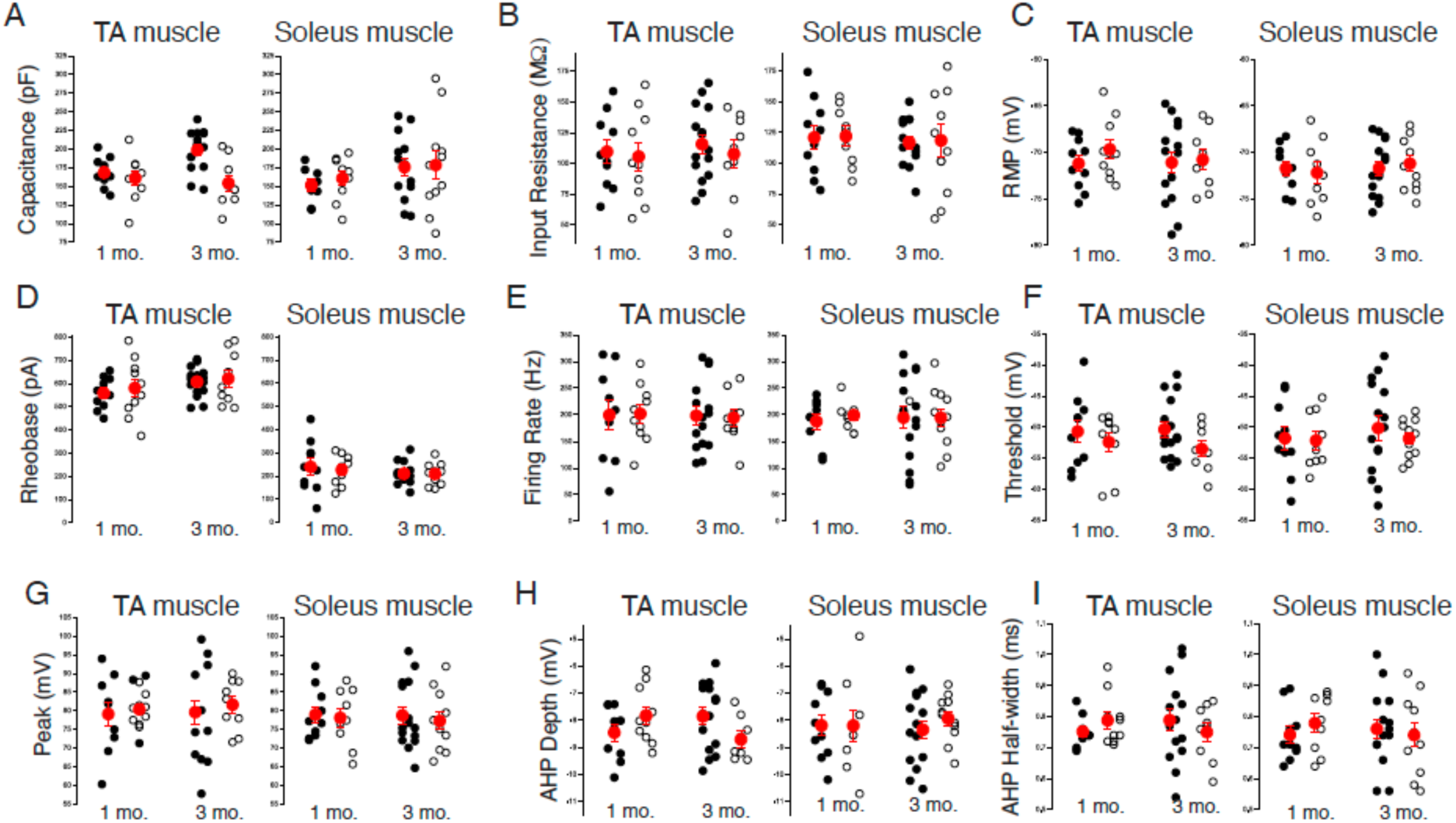
Retro-labelled motor neurons in *Sod1^G85R/G85R^* mice have electrophysiological properties that are indistinguishable from WT mice at one and three months of age. Retro-labeled motor neurons were recorded from acute spinal cord slices. One day prior to recording, Evan’s blue dye was injected into either tibialis anterior (*left columns*) or soleus (*right columns*) muscles. Neurons were studied at both 1 and 3 months of age. Closed circles represent data from individual recordings of neurons from WT mice and open circles from *Sod1^G85R/G85R^* mice. Average values together with standard errors are shown (red). **(A)** Capacitance (pF); **(B)** input resistance (MΩ); **(C)** resting membrane potential (mV); **(D)** rheobase; (pA); **(E)** firing rate (Hz); **(F)** threshold to trigger an action potential (mV); **(G)** peak of action potential (mV); **(H)** peak of action potential after-hyperpolarization (AHP) (mV); **(I)** AP half-width from the start to resting membrane potential (ms).

### *Sod1^G85R/G85R^* mice have disrupted muscle innervation

To assess the integrity of neuromuscular junctions (NMJs) in WT and *SOD1^G85R/G85R^* TA muscles, we performed immunohistochemistry for axonal class III β-tubulin (TUBB3), presynaptic vesicular acetylcholine transporter (VAChT), and postsynaptic acetylcholine receptors (AChRs). We found that, in 3-month-old WT mice, most postsynaptic sites are completely covered by presynaptic nerve terminals, but in age-matched *SOD1^G85R/G85R^* mice, many NMJs are partially or completely denervated (Figure 5A, Extended Video 5-1 and 5-2). Based on the fraction of postsynaptic area occupied by motor nerve terminals, we categorized individual NMJs as 100%, >50%, <50%, or 0% innervated (Figure 5A) and quantified the number of NMJs in each of these categories at 1, 3, and 6 months of age. In WT mice, over 85% of all NMJs are fully innervated and more than 95% are at least >50% innervated, irrespective of age or sex of the animals (Figure 5B). Similar results were obtained for 1-month old *Sod1^G85R/G85R^* mice, but NMJs in these mice exhibited increasing degrees of denervation at 3 and 6 months of age, resulting in about half of all synapses not being fully innervated and significant increases in the number partially or fully denervated NMJs (Figure 5B). Consistent with complete motor axon withdrawal from a subset of postsynaptic sites, *Sod1^G85R/G85R^* mice exhibit an accordingly increased number of axon retraction bulbs in the TA (Extended Figure 5-1). Comparison of male and female mice revealed that denervation is more severe in males than in females at 3 months; at 6 months, denervation in males does not progress any further, with ∼50% of NMJs remaining fully innervated, while severity of the phenotype in female mice increases and reaches the level of males (Figure 5B, C). Thus, *Sod1^G85R/G85R^* animals exhibit loss of neuromuscular innervation, and this neurodegenerative phenotype progresses more quickly in male mice than in female mice. We also compared NMJ innervation in 3-month-old male *Sod1^G85R/G85R^* mice to age- and sex-matched heterozygous *Sod1^G85R/+^* mice and homozygous knock-out *Sod^-/-^* mice. We found that *Sod1^G85R/+^* mice are indistinguishable from WT, while *Sod^-/-^* mice phenocopy *Sod1^G85R/G85R^* animals (Figure 5D), supporting the idea that loss of SOD1 function underlies muscle denervation in *Sod1^G85R/G85R^* mice.

**Figure 5.**
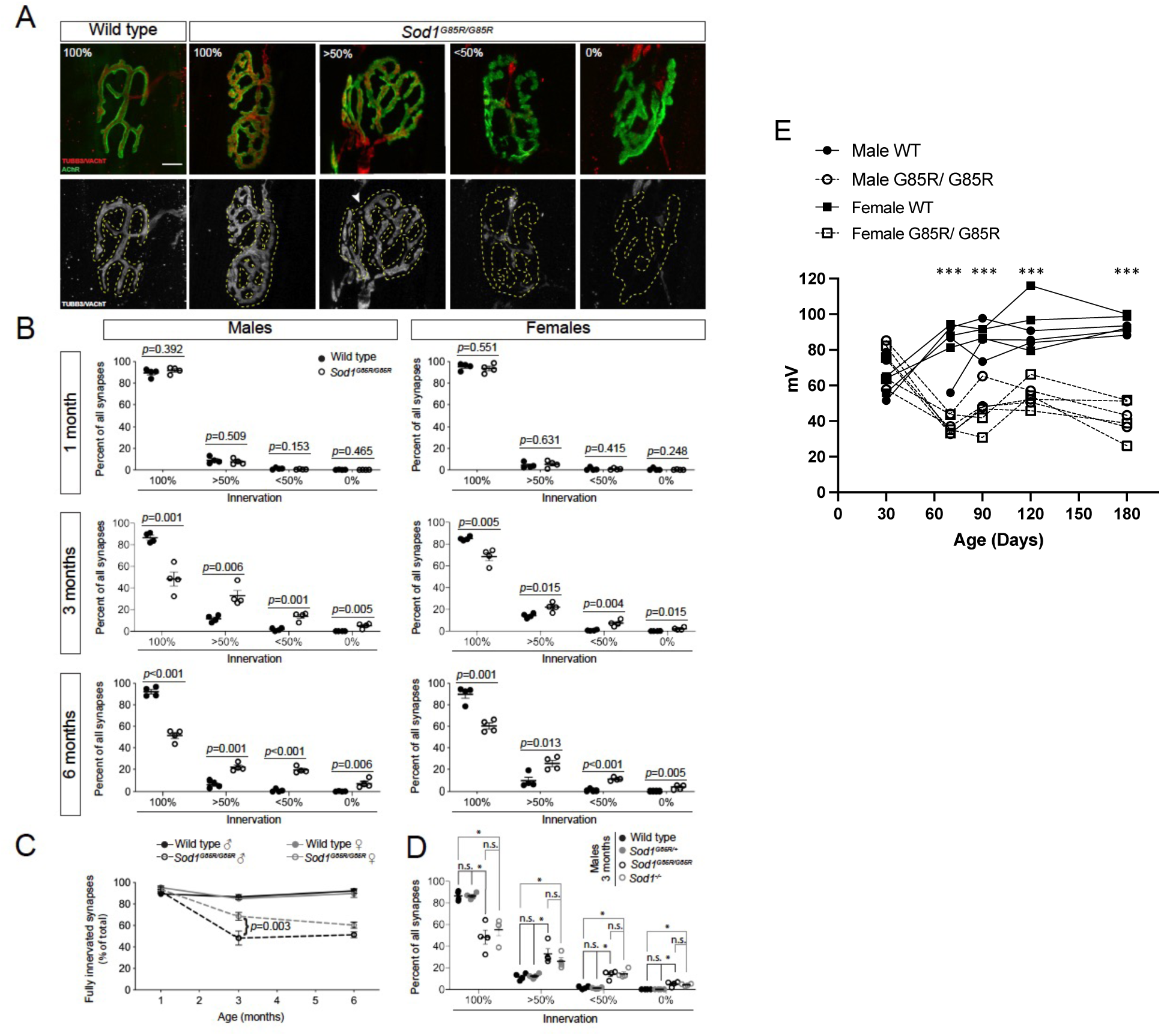
Partial denervation of neuromuscular junctions in *Sod1^G85R/G85R^* mice. **(A)** Longitudinal sections of TA muscles from 3-month-old, male WT and *Sod1^G85R/G85R^* mice were stained with antibodies against class III β-tubulin (TUBB3) and vesicular acetylcholine transporter (VAChT) to visualize motor axons and nerve terminals, respectively. Postsynaptic AChRs were visualized using fluorescent α-bungarotoxin. Bottom row shows isolated presynaptic immunostaining relative to postsynaptic area (dashed outline). NMJs in *Sod1^G85R/G85R^* mice, but not WT, are partially denervated, and the representative images show synapses with 100% presynaptic coverage of the postsynaptic site, between 50% and 100%, between 0% and 50%, and 0%. **(B)** Quantification of NMJs in in different innervation categories for male and female mice at different ages shows significantly decreased innervation in 3- and 6-month-old *Sod1^G85R/G85R^* mice of both sexes, as compared to WT (n = 4 animals per sex, age, and genotype). **(C)** Quantification of fully innervated synapses over time shows normal innervation of NMJs in *Sod1^G85R/G85R^* mice at 1 month and significant denervation compared to WT at 3 and 6 months, with a stronger phenotype in male mice than in females at 3, but not 6 months (n = 4 animals per sex, age, and genotype). **(D)** Quantification of NMJ innervation in 3-month-old, male WT, *Sod1^G85R/+^*, *Sod1^G85R/G85R^*, and *Sod1^-/-^* mice shows normal innervation in *Sod1^G85R/+^* mice and significantly reduced innervation in *Sod1^-/-^* mice when compared to WT; denervation in *Sod1^-/-^* mice is comparable to the *Sod1^G85R/G85R^* phenotype (n = 4 animals per genotype). Asterisks indicate statistically significant differences based on ANOVA and Holm’s post hoc test; n.s. indicates not significant. Scale bar: 10 μm. Error bars indicate SEM. **(E)** Compound muscle action potentials in TA muscles of *Sod1^G85R/G85R^* mice decline between 30 to 70 days of age but then remain stable through at least 180 days. CMAP data for both males and females (n=3 of each) genotype is shown. CMAP data indicates denervation of a subset of *Sod1^G85R/G85R^* neuromuscular junctions occurs after 1 month of age, then innervation levels remain relatively constant through 6 months of age. (t-test ***p < 0.0001).

To further interrogate the integrity of motor units in *Sod1^G85R/G85R^* mice, compound muscle action potentials (CMAP) were evaluated in mice at 30 to180 days of age (Figure 5E). The physiological output from TA muscles were measured and peak mV amplitude compared between *Sod1^G85R/G85R^* and WT mice. At P30 peak amplitudes between *Sod1^G85R/G85R^* and WT mice were not distinguishable, but by P70 CMAPs in *Sod1^G85R/G85R^* mice were reduced significantly compared to WT. CMAP values remained relatively constant between 70 and 180 days and there was no difference between sexes. The CMAP measurements confirm loss of motor unit function in *Sod1^G85R/G85R^* mice between P30 and P70. NMJ denervation and CMAP reduction in *Sod1^G85R/G85R^* mice plateau between 3 and 6 months (although female *Sod1^G85R/G85R^* NMJ denervation proceeds more slowly but reaches a 50% level similar to males at 6 months). The partial loss of motor units that stabilizes at ∼P70 is consistent with our observations of abnormal mouse twitching or tremors upon suspension, which begins at ∼P70 in *Sod1^G85R/G85R^* mice and does not progress to a more severe phenotype such as paralysis.

### *Sod1^G85R/G85R^* male mice have transiently smaller myofiber size early in development

To determine the impact of the *Sod1^G85R/G85R^* knock-in on myofiber size we measured the minimum Feret diameter of TA myofibers at 1, 3, and 6 months of age in males and females (Figure 6). Representative images (Figures 6A, B) show laminin immunohistochemistry of TA cross-sections that delineate the basal lamina surrounding myofibers of *Sod1^G85R/G85R^* and WT male and female mice, which was used to measure fiber diameters. In male *Sod1^G85R/G85R^* mice, we observed a transient reduction in myofiber size at 1 and 3 months of age, but not at 6 months. The average myofiber diameter of *Sod1^G85R/G85R^* male mice was significantly reduced compared to WT males (Figures 6C, E, G, I). In males, myofibers in *Sod1^G85R/G85R^* mice at 1 month of age were ∼20% smaller than WT (WT= 38.4 μm, *Sod1^G85R/G85R^* = 30.6 μm; p=0.004) and at 3 months of age were ∼10% smaller (WT= 45.4 μm, *Sod1^G85R/G85R^*= 40.7 μm; p=0.02), while 6 months fiber size is equivalent (WT = 43.6 μm, *Sod1^G85R/G85R^*= 43.7 μm; p=0.95). In females we noted a similar trend toward a smaller myofiber size in *Sod1^G85R/G85R^* mice, particularly at one month of age (WT= 36.05 μm, *Sod1^G85R/G85R^* = 32.4 μm; p=0.12) and 6 months of age (WT= 43.0 μm, *Sod1^G85R/G85R^*= 40.5 μm; p=0.16); however, these differences did not reach statistical significance (Figures 6D, F, H, J). The sex differences in fiber diameter, which is more profound in *Sod1^G85R/G85R^* male compared to female mice, is consistent with the sex difference observed in the NMJ analysis. The smaller muscle fiber size in 1-month *Sod1^G85R/G85R^* mice could reflect an intrinsic muscle phenotype that precedes the reduced NMJ integrity that is evident at 3 months.

**Figure 6.**
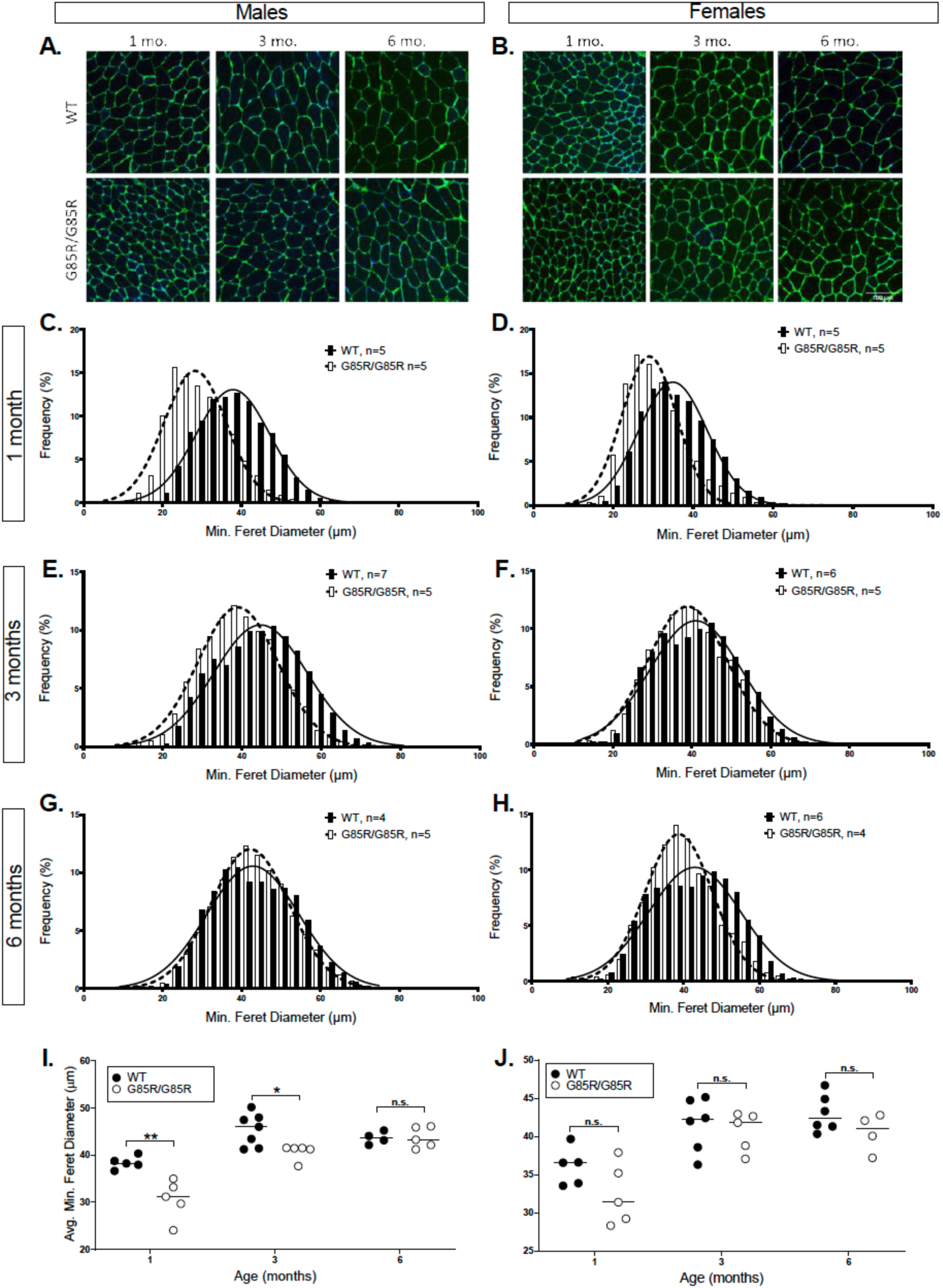
Myofiber size in *Sod1^G85R/G85R^* and WT muscle at one, three and six months of age. **(A, B)** Representative images show laminin immunohistochemistry of TA cross-sections used to calculate minimum Feret diameter for males and females at 1, 3, and 6 months of age. Pooled frequency distribution of minimum Feret diameter of myofibers from tibialis anterior (TA) muscles in male **(C, E, G)** and female **(D, F, H)** mice at 1, 3, and 6 mo. of age, respectively. **(I)** Male *Sod1^G85R/G85R^* mice at 1 month and 3 months of age show significant average myofiber size reduction, but no consistent difference is observed at 6 months (1 mo: WT = 38.4, n = 5; *Sod1^G85R/G85R^*= 30.6, n = 5, p = 0.004. 3 mo: WT = 45.4, n = 7; *Sod1^G85R/G85R^*= 40.7, n = 5, p = 0.02. 6 mo: WT = 43.6, n = 4; *Sod1^G85R/G85R^* = 43.7, n = 5, p = 0.95). **(J)** No consistent differences in average myofiber size are observed at any age in female *Sod1^G85R/G85R^* muscle (1 mo: WT = 36.05, n = 5; *Sod1^G85R/G85R^*= 32.4, p = 0.12, n = 5; 3 mo: WT = 41.5, n = 6; *Sod1^G85R/G85R^* = 40.7, n = 5, p = 0.65; 6mo: WT= 43.0, n = 6; *Sod1^G85R/G85R^* = 40.5, n = 4, p = 0.16).

### *Sod1^G85R/G85R^* MN transcriptomes reveal up-regulation of cholesterol, sterol and lipid pathway genes compared to WT

Given the emergence of a neurodegenerative phenotype in *Sod1^G85R/G85R^* mice after one month of age, we performed motor neuron transcriptome analyses at several ages to determine which genes are differentially expressed in mutant and WT mice during this age period. We used laser capture microdissection to isolate MNs from lumbar spinal cords of mice and performed RNA-seq transcriptional analysis to determine differences in RNA expression among *Sod1* mutant and WT mice (all males) at three different ages: P30, P70 and P120 days. We compared gene expression patterns among homozygous *Sod1^G85R/G85R^* and *Sod1^-/-^* to WT mice. In addition, we compared WT mice to the widely used mouse strain expressing a human *SOD1^G93A/+^* transgene that causes a severe ALS phenotype. As shown in Figure 7, at P30, P70 and P120, genes are differentially up- and down-regulated in the *Sod1^G85R/G85R^* and *Sod1^-/-^* mutants compared to WT, with the extent of differential gene expression increasing with age. By comparison, the degree of altered gene expression in MNs of *TgSOD1^G93A/+^* mutants compared to WT, was greater overall than those associated with either *Sod1^G85R/G85R^* or *Sod1^-/-^* at P120. Consistent with our analyses of NMJ denervation (Figure 5D), we find similar gene expression patterns in MNs of *Sod1^G85R/G85R^* and *Sod1^-/-^* mice. Detailed transcriptome comparison data sets are listed in the Extended Table 7-2.

**Figure 7.**
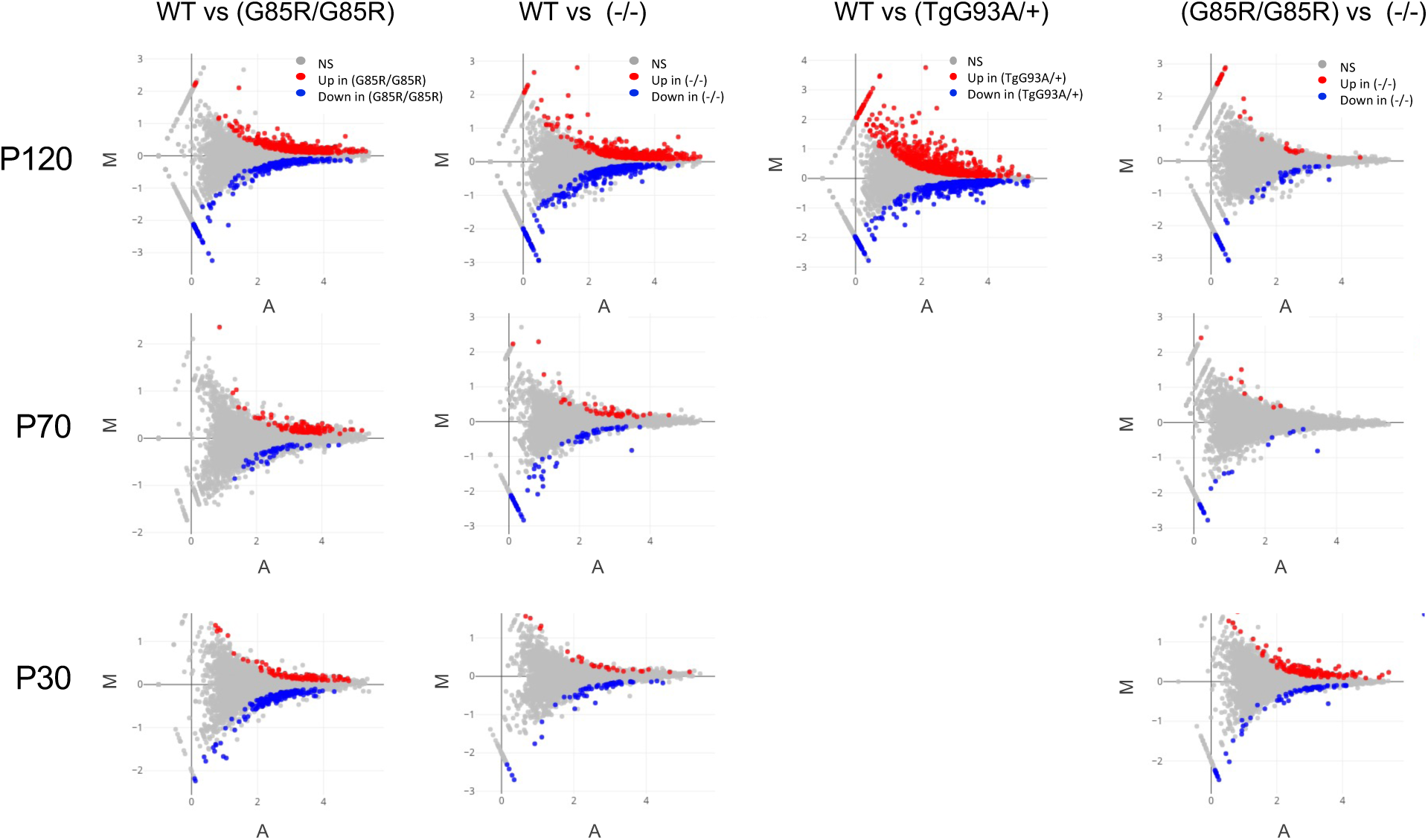
Differential gene expression in motor neurons of *Sod1* mutants compared to WT mice. Laser capture microdissection was used to collect motor neurons from the lumbar spinal cords of homozygous mutant *Sod1^G85R/G85R^*, *Sod1^-/-^*, heterozygous transgenic mutant *TgSOD1^G93A/+^* and WT mice at the ages shown (P120, P70, P30 days). RNA sequencing and differential gene expression analyses were performed to identify genes that are significantly up (red dots) or down (blue dots) regulated in each mutant population compared to WT cohorts (padj < 0.05; gray dots no significant difference between genotypes). Also shown are differential expression plots comparing *Sod1^G85R/G85R^* and *Sod1^-/-^* mice. M: Log2 Fold Change; A: Log10 Normalized Mean Read Counts.

Hierarchical cluster analyses of *Sod1^G85R/G85R^* or *Sod1^-/-^* mice compared to WT at P120 showed the degree of overlap between these ALS mutants and highlighted genes that are commonly altered. (Extended Figure 7-1). Volcano plots of differentially expressed genes (Figure 8A) further demonstrate common patterns of altered gene expression in *Sod1^G85R/G85R^* and *Sod1^-/-^* mice compared to WT. We identified 254 up-regulated genes and 69 down-regulated genes that are common to both *Sod1^G85R/G85R^* and *Sod1^-/-^* MNs compared to WT expression levels (Figure 8B, Extended Table 8-1). Among the 254 genes commonly up-regulated in *Sod1^G85R/G85R^* and *Sod1^-/-^* MNs, there is an enrichment of genes involved in in sterol, cholesterol, and lipid biosynthetic and metabolic processes. For example, of the genes associated with the GO terms “sterol biosynthetic process” (40 genes) and “cholesterol biosynthesis process” (33 genes), more than half are up-regulated in both the *Sod1^G85R/G85R^* and *Sod1^-/-^* MNs compared to WT controls (significance reflected in the false discovery rates of 1.9e-22, and 2.02e-19, respectively). The 69 commonly down-regulated genes do not reveal pathways that are as strongly affected, with neurotransmitter receptor and postsynaptic signal transmission genes the most likely involved (false discovery rate 0.0018). Similar Gene Ontology and KEGG Pathway analyses combining both up- and down-regulated genes that are common to both *Sod1^G85R/G85R^* and *Sod1^-/-^* MNs also show steroid and cholesterol biosynthesis and metabolic pathways as the most prominent processes affected at P120 (Figure 8C). The changes in cholesterol and sterol biosynthetic and metabolic processes begin as early as P70, but not P30, (Figure 8C and Extended Figure 8-1). Differential gene expression does occur in both *Sod1^G85R/G85R^* and *Sod1^-/-^* MNs compared to WT and compared to each other at P70 and P30 (Figure 7, Extended Figures 7-2, 7-3, 7-4, 8-1, 11-3, Extended Tables 7-2, 8-1), but these changes are not strongly associated with additional pathways or biological processes.

**Figure 8.**
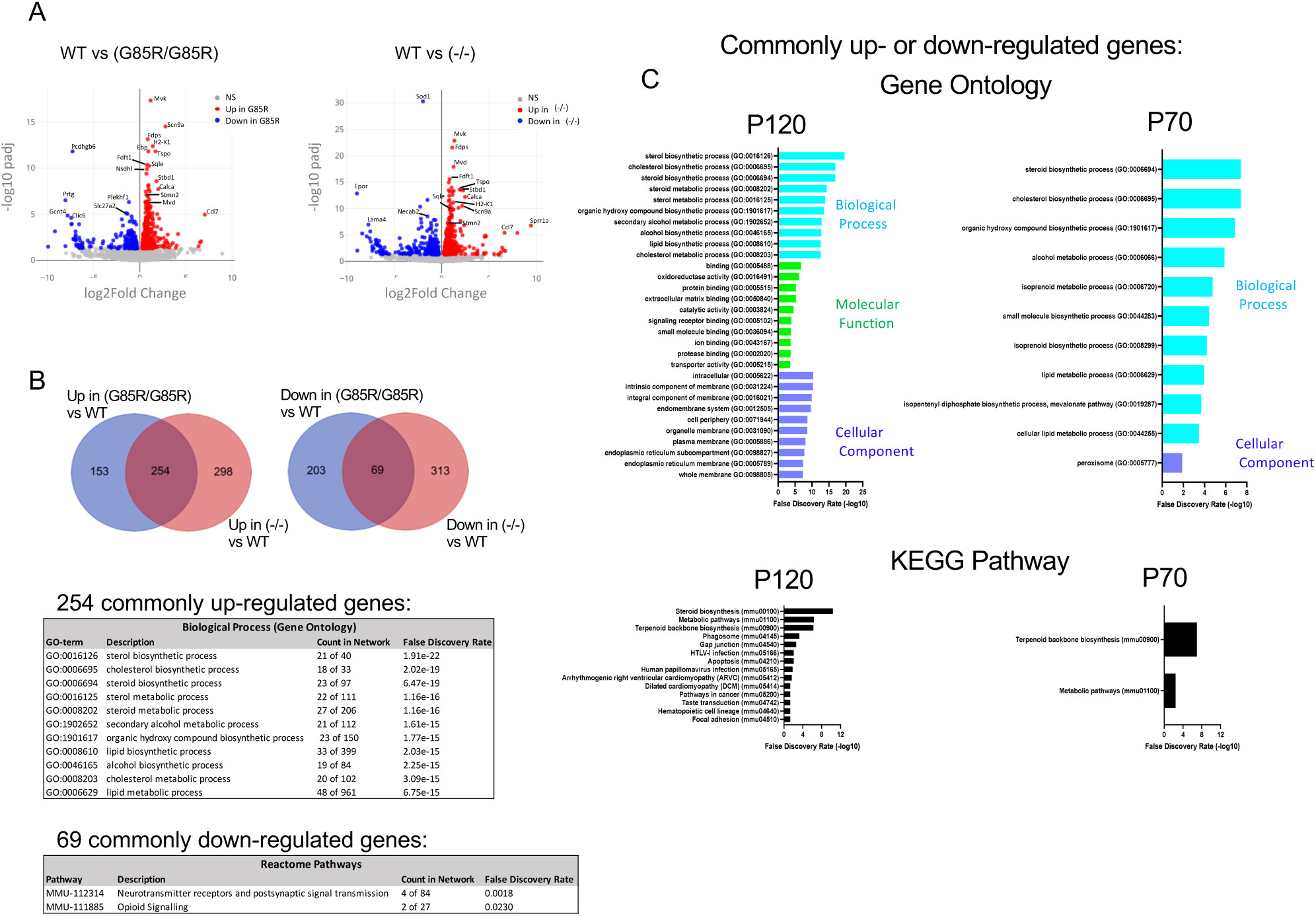
Cholesterol and sterol biosynthesis and metabolic pathway genes are significantly upregulated in both *Sod1^G85R/G85R^* and *Sod1^-/-^* motor neurons compared to WT. Differential expression analyses of mRNAs in day P120 *Sod1^G85R/G85R^* and *Sod1^-/-^* LCM-captured motor neurons compared with WT show an overlapping pattern of gene expression. **(A)** Volcano plots showing significantly up- and downregulated genes in each day P120 mutant (padj < 0.05). Many of the most significantly upregulated genes in *Sod1^G85R/G85R^* mutants compared to WT are also upregulated in *Sod1^-/-^* mutants compared to WT. **(B)** Venn diagrams show that 254 genes are upregulated at P120 in both *Sod1^G85R/G85R^* vs WT and *Sod1^-/-^* vs WT while 69 genes are downregulated in both *Sod1^G85R/G85R^* vs WT and *Sod1^-/-^* vs WT. Pathway analyses of the commonly upregulated genes reveals a highly significant (indicated by low false discovery rate) upregulation of genes associated with sterol and cholesterol biosynthetic pathways and other lipid-associated pathways. Count in network indicates the number of genes associated with each GO-term that are present in the upregulated gene set. The most significant pathways associated with commonly downregulated genes in the mutants involve neurotransmitter receptors and postsynaptic signal transmission and opioid signaling. **(C)** Additional Gene Ontology and KEGG Pathway analyses using combined up- and downregulated data sets from each differential expression analysis showing processes, functions, or cellular components most affected in both *Sod1^G85R/G85R^* and *Sod1^-/-^* mutant motor neurons relative to WT. Several similar processes are commonly affected at P70 as well as P120 in the *Sod1^G85R/G85R^* and *Sod1^-/-^* motor neurons.

To further explore the impact of the *Sod1^G85R/G85R^* and *Sod1^-/-^* mutations on cholesterol synthesis or metabolism in MNs, we specifically evaluated the expression of genes involved in these pathways. At P120, almost all the enzymes involved in the synthesis of cholesterol from acetyl CoA (Figure 9A) are significantly up-regulated in both *Sod1^G85R/G85R^* and *Sod1^-/-^* MNs compared to WT (Figure 9B, Extended Table 9-1). The patterns of gene expression changes are nearly identical for the *Sod1^G85R/G85R^* and *Sod1^-/-^* mutants. In stark contrast, in P120 *TgSOD1^G93A/+^* mice, many of these same genes are down-regulated, indicating that at this point in the pathology of these more severely affected *TgSOD1^G93A/+^* mice, cholesterol synthesis pathways are shutting down. We similarly looked at the genes involved in some pathways of cholesterol use and clearance (Extended Figures 9-1A, 9-1B, Extended Table 9-1). Of the genes examined, only *Ldlr* (low-density lipoprotein receptor) is significantly up-regulated in both *Sod1^G85R/G85R^* and *Sod1^-/-^* MNs. *Ldlr* encodes a protein involved in cholesterol uptake into cells, consistent with the overall effect of increasing cholesterol in the cells. Other genes examined either did not change significantly in expression or were not expressed at all. *Abcg1* (ATP binding cassette subfamily G1), encoding a protein involve in cholesterol efflux from cells, is significantly down-regulated in *Sod1^-/-^* cells, with a similar trend in *Sod1^G85R/G85R^* cells. Notably, in *TgSOD1^G93A/+^* cells, *Ldlr* is down-regulated, concomitant with an up-regulation of several genes involved in lipid oxidation and clearance either through efflux or storage as cholesterol esters or lipid droplets, indicating cellular mechanisms to reduce cholesterol are engaged in *TgSOD1^G93A/+^* cells.

**Figure 9.**
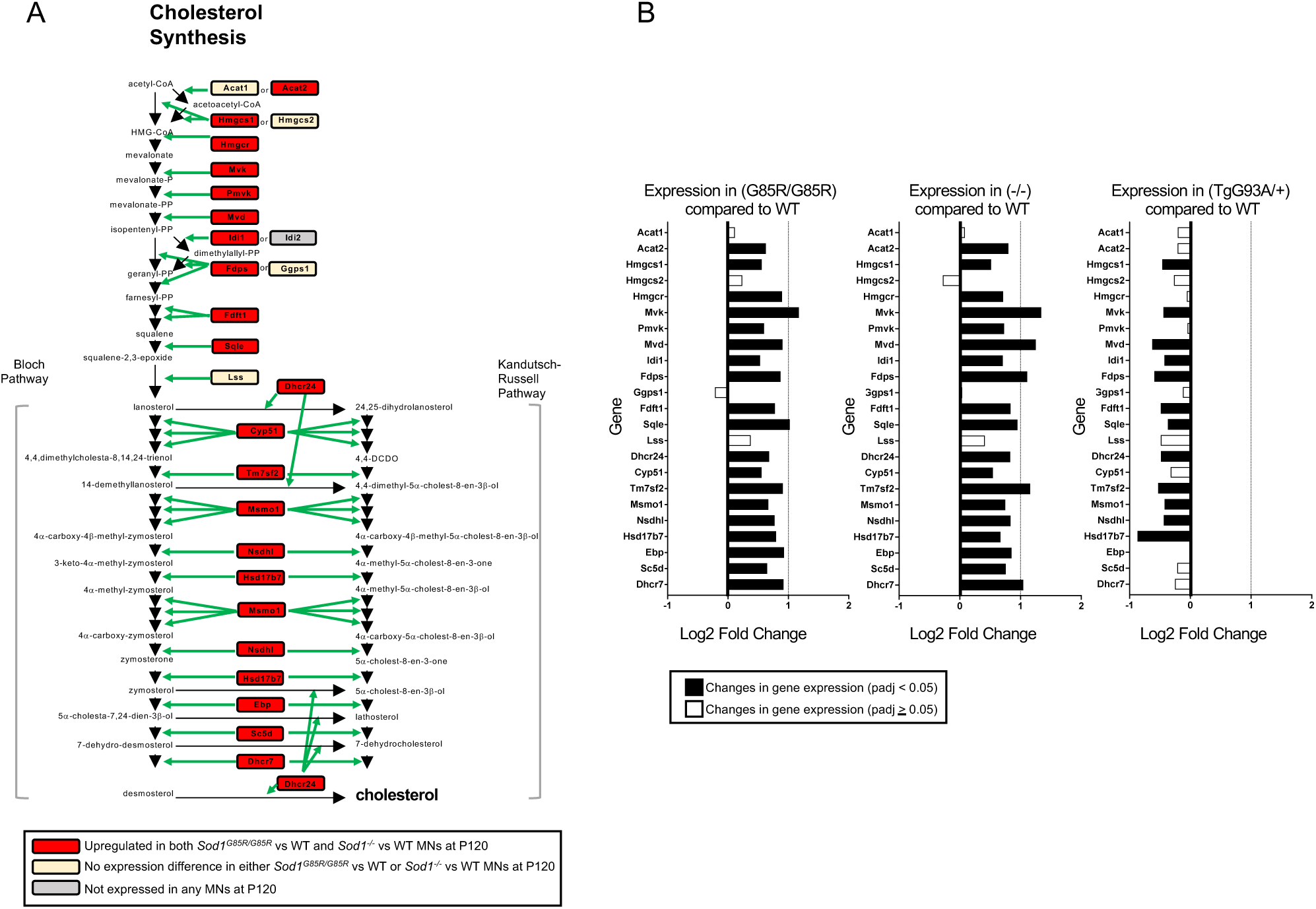
Genes involved in cholesterol synthesis are upregulated in *Sod1^G85R/G85R^* and *Sod1^-/-^* motor neurons and these expression profiles are different from those of *TgSOD1^G93A/+^* motor neurons. **(A)** Genes for proteins involved in cholesterol synthesis pathways (in boxes) and the pathway intermediates are shown. Genes highlighted in red are upregulated at P120 in both *Sod1^G85R/G85R^* vs WT and *Sod1^-/-^* vs WT motor neurons; yellow boxes: no significant difference in expression in mutant vs WT; gray boxes: not expressed. **(B)** The differential expression of the genes shown in **(A)** in each of the mutant vs WT comparisons is shown. Black bars: significant expression differences (padj < 0.05); white bars: differences are not significant. The overall pattern differential expression at P120 of *Sod1^G85R/G85R^* vs WT and *Sod1^-/-^* vs WT are very similar, while the more severe *TgSOD1^G93A/+^* mutation leads to an opposite pattern of gene expression changes at the same age.

Other in lipid-related pathway genes are also differentially expressed in similar patterns in *Sod1^G85R/G85R^* and *Sod1^-/-^* MNs compared to WT at P120; most of these expression differences are not observed in *TgSOD1^G93A/+^* MNs compared to WT (Extended Figure 9-1C, Extended Table 9-1). Compared to WT, *Sod1^G85R/G85R^* and *Sod1^-/-^* MNs up-regulate *Insig1* (insulin induced gene 1), a protein involved in regulating cholesterol synthesis; *TSPO* (translocator protein), which promotes cholesterol movement into mitochondria; *Elolv7* (elongation of very long chain fatty acids protein), the rate-limiting step in long-chain fatty acid elongation; and *Abcg2* (ATP binding cassette subfamily G2), a cholesterol-sensitive broad-substrate transporter of heme, vitamins, toxins and drugs across membranes. *Sod1^G85R/G85R^* and *Sod1^-/-^* MNs down-regulate *Slc27A2* (solute carrier family 27 member 2), which converts free long chain fatty acids to acetyl-CoA esters and is involved in lipid biosynthesis and fatty acid degradation. These gene expression changes further demonstrate that *Sod1^G85R/G85R^* and *Sod1^-/-^* mutations have the similar effects on cholesterol and lipid-related pathways in MNs, which are different from effects observed in *TgSOD1^G93A/+^* MNs.

### Transcriptional changes in *Sod1^G85R/G85R^* MNs include changes in ion channel and transporter expression

Among the more significant RNA expression changes in both *Sod1^G85R/G85R^* mice and *Sod1^-/-^* MNs compared to WT are up-regulation of the sodium channel *Scn9A* (Nav1.7; voltage-gated sodium ion channel alpha subunit 9), which is involved in nociception signaling, and up-regulation of *Pkd2l1* (polycystic kidney disease 2-like 1 protein), the non-selective cation channel which can act as a chemosensor or mechanosensor in spinal cord ependymal cells and might play a role in sour taste perception (Figure 10). Interestingly *Pkd2l1* is also up-regulated in both *Sod1^G85R/G85R^* mice and *Sod1^-/-^* MNs at P70 (Extended Figures 8-1, 11-3). Up-regulation of *Scn9A* but not *Pkd2l1* also occurs in *TgSOD1^G93A/+^* MNs. At P120, *Sod1^G85R/G85R^* mice and *Sod1^- /-^* MNs have strikingly similar patterns of gene up-regulation in cytoskeletal genes (*Tubb* (tubulin beta) genes and *Stmn2*, (stathmin 2, a regulator of neurite growth and axon regeneration)), the *Calca* and *Calcb* genes encoding the neuromodulator and vasodilator CGRP2 (calcitonin gene-related peptide 2; the predominant differentially expressed *Calca* transcripts our data encode CGRP and not calcitonin, an alternate splice product of the *Calca* gene), and the major histocompatibility complex gene *H2-K1*, which functions in the immune response and is also protective for damaged MN axons and against astrocyte-mediated toxicity in ALS MNs (Thams et al., 2009; Song et al., 2016). Down-regulation of some genes also occurs in P120 *Sod1^G85R/G85R^* MNs, among them Ca^2+^- binding proteins such including the signaling scaffold protein *Necab2* (N-terminal EF-hand calcium binding protein 2), and the calcium-binding protein *Pcdhgb6* (protocadherin gamma-B6) which could function in cell adhesion and neural connections. P120 *Sod1^-/-^* MNs also undergo down-regulation of *Necab2*. *Tubb6* and *H2-K1* are also up-regulated in *TgSOD1^G93A/+^* MNs. Overall, compared to WT, the type and direction of gene expression changes observed in *Sod1^G85R/G85R^* and *Sod1^-/-^* MNs are very similar to each other and distinct from altered gene expression in *TgSOD1^G93A/+^* MNs.

**Figure 10.**
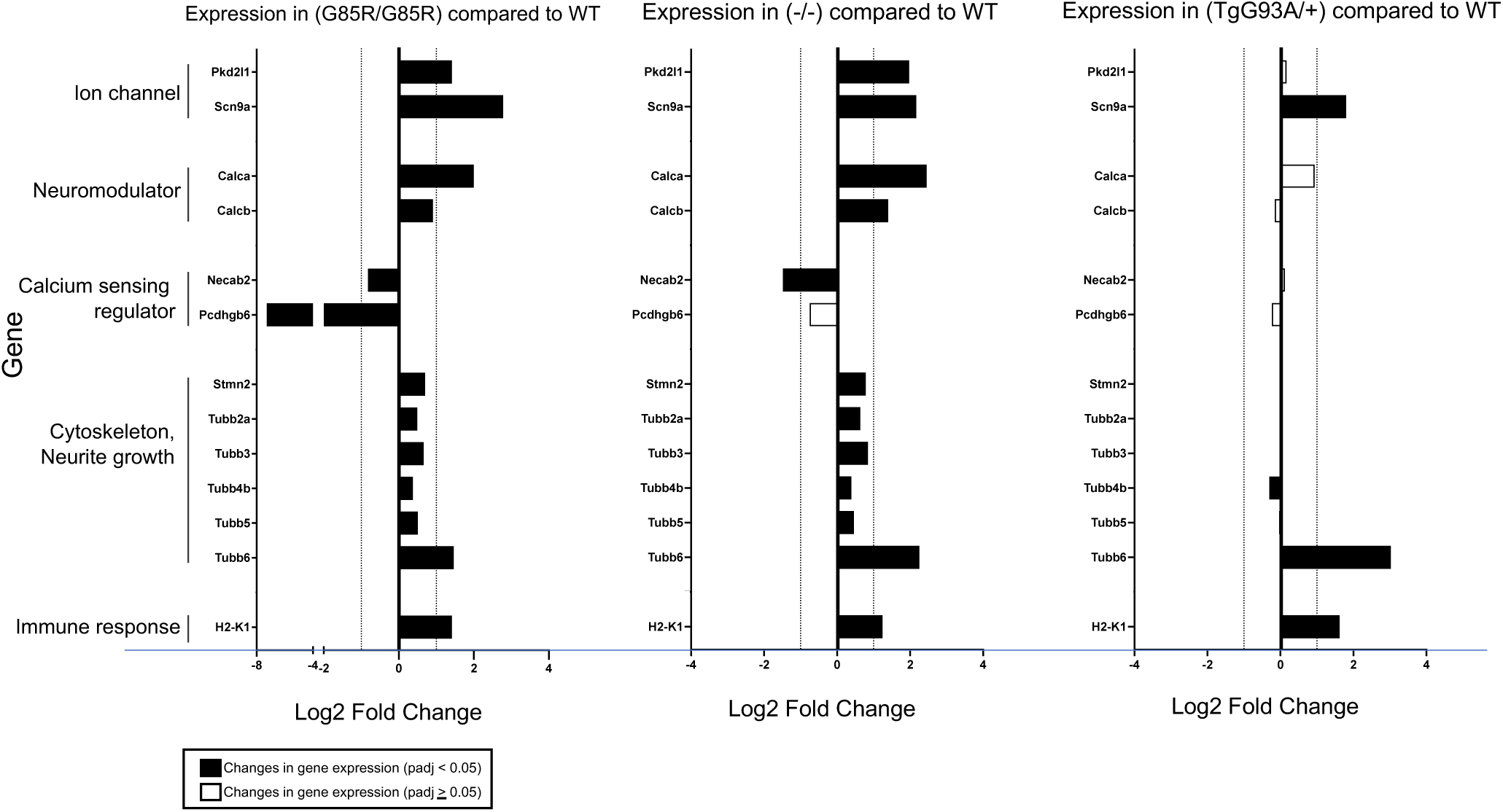
*Sod1^G85R/G85R^* and *Sod1^-/-^* motor neurons exhibit similar altered expression patterns for genes other than those involved in cholesterol synthesis. Several genes showing a highly significant differential expression in P120 *Sod1^G85R/G85R^* vs WT and *Sod1^-/-^* vs WT motor neurons are shown, along with *TgSOD1^G93A/+^* vs WT motor neuron expression comparisons. Black bars: significant expression differences (padj < 0.05); white bars: differences are not significant. As with cholesterol pathway genes, the differential expression patterns for many of these is similar in *Sod1^G85R/G85R^* vs WT and *Sod1^-/-^* vs WT motor neurons and distinct from *TgSOD1^G93A/+^* vs WT motor neurons at this age, with only a few overlapping patterns.

To further examine the effects of *Sod1* mutations on the expression of ion channels and transporters in MNs, we evaluated the RNA expression of 877 genes that are associated with GO terms related to ion channel, transmembrane transporter, and neurotransmitter genes (Extended Table 11-1). Of these 877 genes, 148 genes are differentially expressed in MNs of one or more of *Sod1* mutants compared to WT (Figure 11A, Extended Figures 11-1 and 11-2). There is ∼60% overlap (32 of 48) in genes differentially expressed in *Sod1^G85R/G85R^* and *Sod1^-/-^* MNs compared to WT (Figure 11B). By contrast, there is a much lower overlap in MN genes differentially expressed in *TgSOD1^G93A/+^* compared to WT and *Sod1^G85R/G85R^* or *Sod1^-/-^* compared to WT (Figure 11C, D, E).

**Figure 11.**
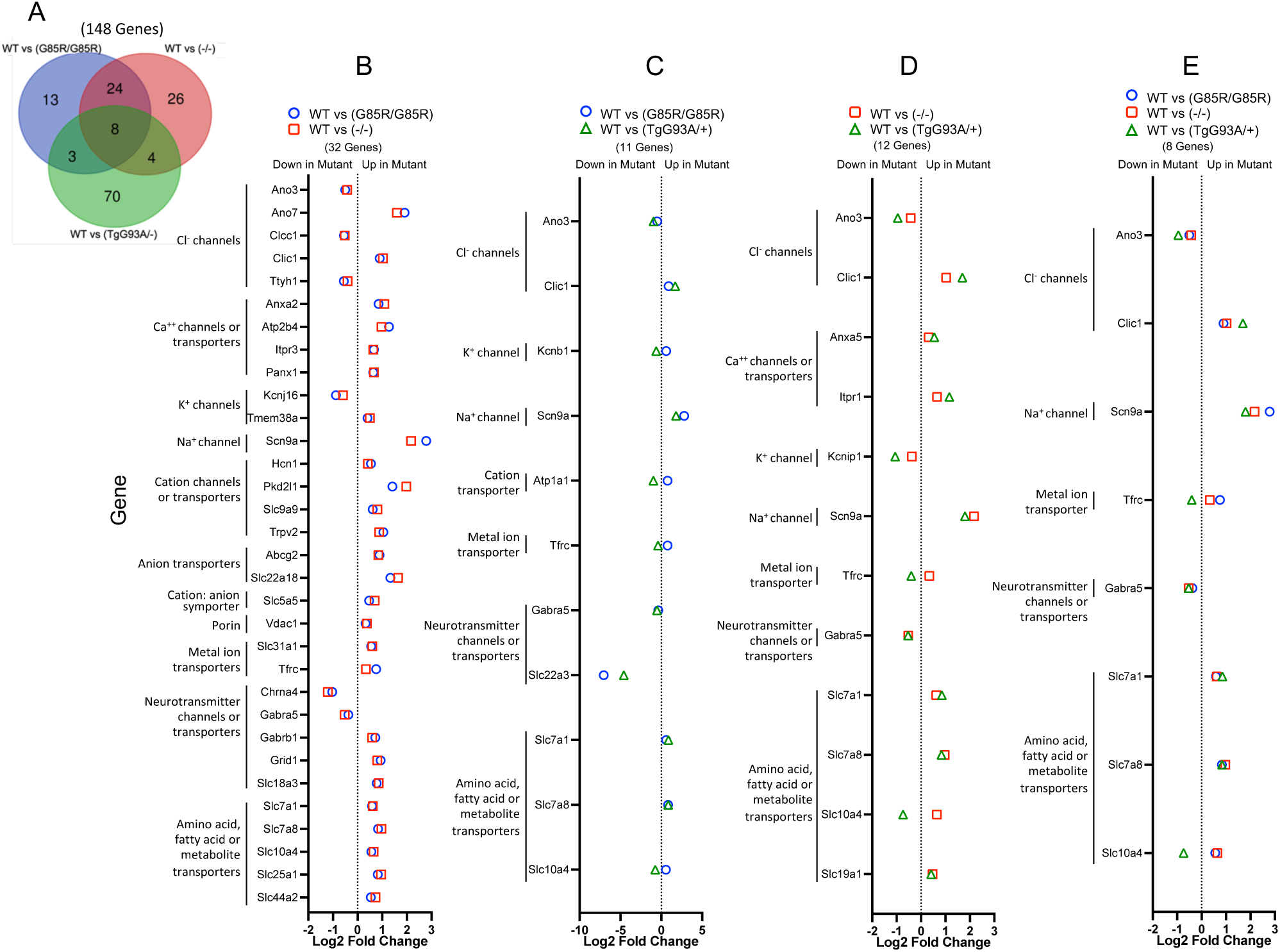
The expression of channel, transporter and neurotransmitter genes are similarly altered in *Sod1^G85R/G85R^* and *Sod1^-/-^* mutant motor neurons as compared to WT. A set of 877 genes associated with ion channel, transmembrane transporter, and neurotransmitter Gene Ontology terms was examined. Of these, 148 are differentially expressed in the LCM MNs from one or more P120 *Sod1* mutant compared to WT (padj < 0.05). **(A)** Venn diagram shows the number of genes that are differentially expressed in P120 *Sod1^G85R/G85R^* vs WT, *Sod1^-/-^* vs WT or *TgSOD1^G93A/+^* vs WT MNs and the number of gene changes the mutants have in common. **(B - E)** Expression differences for the genes listed that are differentially expressed in each of the pairwise comparisons shown. The differential gene expression patterns are highly similar in *Sod1^G85R/G85R^* vs WT and *Sod1^-/-^* vs WT motor neurons, with fewer similarities between *Sod1^G85R/G85R^* vs WT and *TgSOD1^G93A/+^* vs WT or *Sod1^-/-^* vs WT and *TgSOD1^G93A/+^* vs WT.

The similarity of gene expression changes observed in *Sod1^G85R/G85R^* and *Sod1^-/-^* compared to WT mice is consistent with their overall phenotypes. At P120, both *Sod1^G85R/G85R^* and *Sod1^-/-^* mice exhibit a similar mild tremor phenotype but this does not progress significantly with age and paralysis is not a feature of these mutant mice. The gene expression patterns of *Sod1^G85R/G85R^* and *Sod1^-/-^* are distinct from *TgSOD1^G93A/+^* which exhibit prominent tremors at P120 followed, about 2 weeks later, by progressively severe symptoms including paralysis and ultimately death by ∼165 days. Overall, our data indicate that *Sod1^G85R/G85R^* and *Sod1^-/-^* MNs have similar molecular profiles suggesting that SOD1 loss of function is the predominant driver of the *Sod1^G85R/G85R^* phenotype.

### Differential gene expression in ventral and dorsal spinal cord tissues

We used quantitative RT-PCR analysis of mRNA from separated ventral and dorsal regions of lumbar spinal cords to determine whether genes, differentially expressed in *Sod1^G85R/G85R^* and *Sod1^-/-^* LCM-captured MNs, change similarly in other regions of the spinal cord. We compared gene expression in *Sod1^G85R/G85R^* and *Sod1^-/-^* mutants relative to WT in ventral and dorsal tissues, and additionally compared gene expression in ventral and dorsal tissues (Figure12 and Extended Figure 12-1). As expected, *Sod1* mRNA expression is greatly reduced in both the ventral and dorsal *Sod1^-/-^* spinal cord tissues compared to WT, while *Sod1^G85R/G85R^* tissue *Sod1* expression levels are similar to WT. *Sod1* levels are not different between ventral and dorsal tissues of the same genotype. As anticipated, expression of *Chat*, a MN marker, is extremely low and variable in dorsal tissue. There is no significant difference in *Chat* RNA expression in bulk ventral or dorsal tissue of *Sod1^G85R/G85R^* or *Sod1^-/-^* compared to WT, although LCM-enriched large MNs show a slight but significant increase in *Chat* expression in both *Sod1^G85R/G85R^* and *Sod1^-/-^* mice relative to WT. In bulk tissue samples, significant *Chat* expression differences in the small subset of large MN cells might be masked by more uniform expression by other smaller cells in the region.

Several cholesterol synthesis pathway genes that are significantly up-regulated in LCM captured MNs of both *Sod1^G85R/G85R^* and *Sod1^-/-^* compared to WT mice (*Mvk, Mvd, Ebp, Tm7sf2, Tspo,* Figure 9 and Extended Figure 9-1), also show a slight trend toward up-regulation in bulk ventral spinal cord tissue of *Sod1^G85R/G85R^* and *Sod1^-/-^* compared to WT mice, although most differences do not reach significance (i.e. p>0.05) (Figures 12, Extended Figure 12-1; p values for differential expression of these genes are 0.116, 0.121, 0.006, 0.182, 0.002, respectively in *Sod1^G85R/G85R^* vs WT ventral tissue and 0.165, 0.393, 0.167, 0.267, 0.091 in *Sod1^-/-^* vs WT ventral tissue). This trend toward up-regulation among multiple cholesterol pathway genes in the bulk tissues is consistent with up-regulation in MNs of *Sod1^G85R/G85R^* and *Sod1^-/-^* mutants compared to WT that could be masked by more uniform expression in other cells of ventral cord. For most of these genes, expression is higher in ventral spinal cord than dorsal cord in all genotypes.

**Figure 12.**
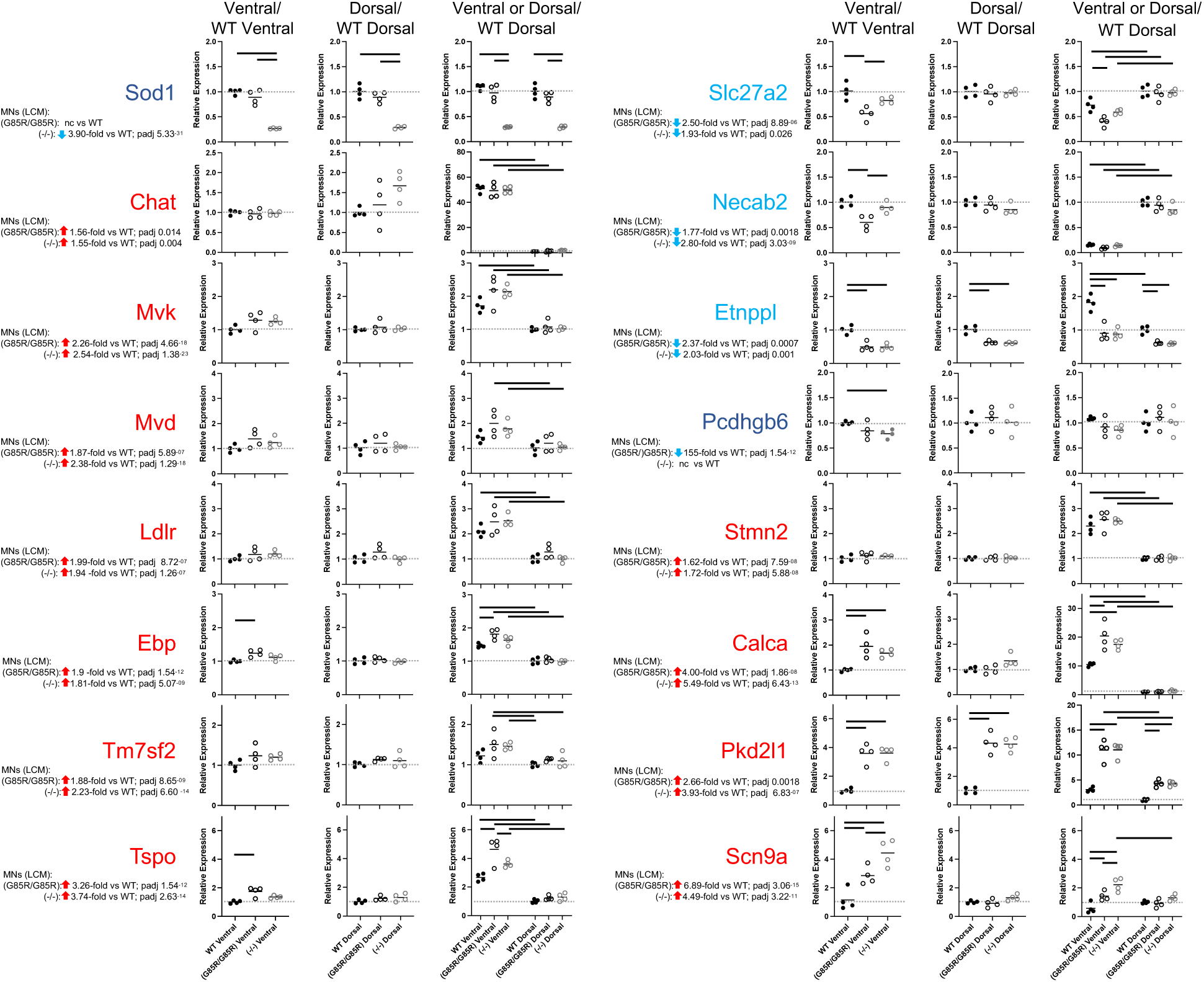
Ventral and dorsal spinal cord tissues exhibit differential expression of cholesterol pathway and other genes, some of which are significantly altered in *Sod1^G85R/G85R^* and *Sod1^-/-^* mutants. **(A)** We used quantitative RT-PCR analyses to examine relative gene expression levels in bulk mRNA from ventral or dorsal regions of lumbar spinal cord tissue collected from P120 WT, *Sod1^G85R/G85R^* and *Sod1^-/-^* mice. Genes examined were among those differentially expressed in *Sod1^G85R/G85R^* vs WT and *Sod1^-/-^* vs WT LCM motor neurons. For each gene, the relative expression from the LCM motor neuron analysis is listed and the expression in the ventral and dorsal tissues is shown in graphs. For each gene, the data shown compares the expression in each tissue to its respective to WT expression (WT = 1). Graphs show i) WT, *Sod1^G85R/G85R^* and *Sod1^-/-^* mutant ventral tissue expression relative to WT ventral tissue; ii) WT, *Sod1^G85R/G85R^* and *Sod1^-/-^* mutant dorsal tissue expression relative to WT dorsal tissue; and iii) both ventral or dorsal WT, *Sod1^G85R/G85R^* and *Sod1^-/-^* tissue expression relative to WT dorsal tissue to show expression differences in each region of the spinal cord. Genes highlighted in red are upregulated in both *Sod1^G85R/G85R^* vs WT and *Sod1^-/-^* vs WT lumbar LCM motor neurons; those in blue are downregulated in both *Sod1^G85R/G85R^* vs WT and *Sod1^-/-^* vs WT lumbar LCM motor neurons; those in dark blue is downregulated only in one of the mutant vs WT LCM motor neurons. Solid black circles: WT, open black circles: *Sod1^G85R/G85R^*; open gray circles: *Sod1^-/-^*, n= 4 mice for each. Bars represent significant differences in gene expression between individual samples (Mean ± SEM; ANOVA, p< 0.05), with only a subset of these displayed for the last panel in each set. Bars representing all significant differences in the third panels are shown in Extended Figure 12-1.

Some cholesterol/lipid pathway genes (*Slc27a2, Necab, Etnppl*) that are significantly down-regulated in LCM MNs of both *Sod1^G85R/G85R^* and *Sod1^-/-^* mice are also significantly down-regulated in the bulk ventral cord tissues of one or both *Sod1^G85R/G85R^* and *Sod1^-/-^* mutants, or trend toward down-regulation. By contrast, in dorsal spinal cord tissue *Slc27a2* and *Necab* genes are not down-regulated, while *Etnppl* (ethanolamine-phosphate phospho-lyase which catalyzes the breakdown of phosphoethanolamine and may play a role in regulating phospholipid levels), is down-regulated in both ventral and dorsal regions of *Sod1^G85R/G85R^* and *Sod1^-/-^* mice compared to WT.

As previously noted, *Stmn2*, which is highly expressed in MNs and plays an important role in neurite outgrowth and neuron development, is higher in LCM MNs of both *Sod1^G85R/G85R^* and *Sod1^-/-^* mice (Figure 10). This difference in *Stmn2* expression is not observed in analysis of bulk ventral or dorsal tissues, perhaps because the analysis is confounded by more uniform *Stmn2* expression in other cells in the surrounding cord tissue (Figures 12, Extended Figure 12-1). Comparing bulk ventral and dorsal tissues, expression of both *Stmn2 and Calca* (encoding CGRP2) is higher in ventral compared to dorsal tissues in all genotypes. *Calca*, and *Scn9a* are up-regulated in *Sod1^G85R/G85R^* and *Sod1^-/-^* ventral tissue compared to WT. *Pkd2l* is up-regulated in both ventral and dorsal tissues of *Sod1^G85R/G85R^* and *Sod1^-/-^* mutants relative to WT implicating a role for this protein throughout the diseased spinal cords.

Together, transcriptional analyses of bulk cord tissue and LCM MNs highlight the similarities in gene expression patterns in spinal cord tissue between *Sod1^G85R/G85R^* and *Sod1^-/-^* mutant mice. We conclude that *Sod1* loss-of-function is driving the phenotypic changes in *Sod1^G85R/G85R^* mutant mice. *Sod1* mutation-associated gene expression differences in spinal cord MNs, which are particularly striking for cholesterol synthesis pathway genes, could play a role in *Sod1*-related pathogenesis or in compensatory stabilization mechanisms to prevent more severe pathology that is a feature of the *TgSOD1^G93A/+^* mouse model of ALS.

## DISCUSSION

Homozygous *Sod1^G85R/G85R^* knock-in mice express normal amounts of *Sod1* RNA but extremely low SOD1 protein and have no detectable SOD1 activity. This is consistent with reduced SOD1 activity in *SOD1^G85R^* patients (Deng et al., 1993), cells, and mice carrying human *SOD1^G85R^* and mouse *Sod1^G86R^* transgenes (Borchelt et al., 1994; Ripps et al., 1995; Wang et al., 2009; Audet et al., 2010). Low SOD1^G85R^ protein accumulation likely reflects protein instability due to its reduced affinity for copper and zinc (Valentine et al., 2005). SOD1^G85R^ is one of the more unstable SOD1 protein variants *in vitro* (protein half-life 7.5 hours compared to 30 h for WT SOD1) (Borchelt et al., 1994).

*Sod1^G85R/G85R^* mice have a mild phenotype that includes reduced body weight compared to *Sod1^G85R/+^* or WT mice, leading to reduced grip strength. *Sod1^G85R/G85R^* mice are not distinguishable from WT in rotarod or open field movement or weight-normalized grip strength. Abnormal *Sod1^G85R/G85R^* mouse behavior is evident from ∼70 days of age as hindlimb twitching when mice are tail-suspended, but this does not progress to severe motor function loss. Reduced lifespan in *Sod1^G85R/G85R^* mice is not due to paralysis, but is instead associated with other health issues often observed in aged mice, similar to WT.

Concurrent with onset of subtle movement abnormalities, *Sod1^G85R/G85R^* mice exhibit ∼30-40% reduction in the proportion of fully innervated NMJs between one and three months in males, one and six months in females, and a parallel loss of functional motor units (CMAPs). At about three (or six) months, this pathophysiology stabilizes, with no further loss of CMAPs or NMJ innervation. Denervation might reflect susceptibility of a particular MN type, as reported in transgenic *SOD1^G85R^* and *SOD1^G93A^* mice, in which fast-fatigable (type-IIb) MNs were selectively vulnerable compared with fast-fatigue-resistant (type-IIa) or slow (type-I) MNs (Pun et al., 2006; Nijssen et al., 2017; Ragagnin et al., 2019). Preferential denervation of fast twitch muscles also occurs in *Sod1^-/-^* mice but without MN loss (Fischer et al., 2012). Very young (6-10 day-old) transgenic S*OD1^G85R^* and low-expressing *SOD1^G93A^* mice exhibit MN electrophysiological abnormalities, some indicative of hypoexcitability or developmental immaturity (Bories et al., 2007; Pambo-Pambo et al., 2009; Filipchuk et al., 2021). We, however, observe no electrophysiological differences between *Sod1^G85R/G85R^* and WT MNs innervating TA or soleus muscles at one or three months, although we only record from MNs that take up dye injected into muscle and thereby likely exclude MNs that have completely withdrawn their axons from postsynaptic sites. Our data indicate that following denervation involving a subset of *Sod1^G85R/G85R^* MNs, there is a period of stability, at least for six-months studied here. *Sod1^G85R/G85R^* MNs surviving at six months could be fast-fatigue-resistant and slow cell types.

Our NMJ denervation and myofiber size observations indicate *Sod1^G85R/G85R^* males are more affected than females at earliest disease stages, although there are no gender differences in motor unit loss (CMAPs). Some transgenic *SOD1^G93A^* rodent studies showed earlier disease onset in males than females (Veldink et al., 2003; Suzuki et al., 2007), but other analyses of *SOD1^G93A^* mouse motor unit or muscle force decline showed no gender differences (Hegedus et al., 2009). In patients, ALS prevalence is greater in males than females (∼1.3:1, M:F). Mechanisms underlying this are unknown, but might involve effects of sex hormones (McCombe and Henderson, 2010; Trojsi et al., 2020) such as changes in androgen receptor function (McLeod et al., 2022). Early gender differences we observe might reflect enhanced male vulnerability in a subset of MNs and offer insight into mechanisms initiating MN loss.

The *Sod1^G85R/G85R^* phenotype, including reduced body weight, twitching that does not progress to paralysis, partial NMJ denervation, motor unit loss, and reduced lifespan, is similar to that described for *Sod1^D83G/D83G^* mice (Joyce et al., 2015), which also express reduced SOD1 protein and very low SOD1 activity. Many of these features, in particular, our NMJ denervation observations, resemble *Sod1^-/-^* mice that have distal axonopathy but without loss of spinal cord MNs (Flood et al., 1999; Shefner et al., 1999; Muller et al., 2006; Fischer et al., 2012). *Sod1^G85R/G85R^* mice therefore largely phenocopy SOD1 loss-of-function, not more severe toxic mutant gain-of-function ALS models.

It is known that cholesterol metabolism is perturbed in human and rodent ALS, however, findings are disparate and challenging to integrate into clear hypotheses about ALS pathogenesis. Elevated cholesterol in patient sera or plasma is reportedly a risk factor for ALS (Bandres-Ciga et al., 2019; Zeng and Zhou, 2019; van Rheenen et al., 2021; Hop et al., 2022). By contrast, elevated circulating cholesterol is reportedly beneficial to ALS patient survival once disease has begun (Ingre et al., 2020; Hartmann et al., 2021). Excessive accumulation of cholesterol metabolites reported in ALS patients and mouse models can be neurotoxic (Hartmann et al., 2021). Our transcriptome analyses reveal enhanced expression of cholesterol biosynthesis genes as a prominent feature of *Sod1^G85R/G85R^* MNs. Surprisingly, different *Sod1* mutants have distinct profiles. In *Sod1^G85R/G85R^* and *Sod1^-/-^* at P120, cholesterol and sterol biosynthesis genes are up-regulated, while in P120 *TgSOD1^G93A/+^* MNs, these genes are down-regulated, and others involved in cholesterol metabolism or clearance are up-regulated.

Our P120 *Sod1^G85R/G85R^* MN transcriptome analyses suggest that activation of cholesterol synthesis could be protective, as this transcriptional activity reflects MNs present after the period of denervation that occurs within three months of age, apparently remaining functional long-term without detectable physiological deficits. Cholesterol is essential for synthesis of steroid hormones, oxysterols, bile acids, membrane lipids and myelin. It is a key component of membrane lipid rafts important for receptors, transporters, enzymatic complexes (Brown and London, 1998; Simons and Toomre, 2000; Simons and Ehehalt, 2002), neurotransmitters and AChRs (Zhu et al., 2006; Allen et al., 2007). Appropriate cholesterol levels are likely critical for MNs to maintain extremely long cell projections and distant synapses (Pfrieger, 2003; Krivoi and Petrov, 2019). Addition of cholesterol to denervated muscles or nerve-muscle explants stabilizes AChR clusters at the NMJ (Willmann et al., 2006), and treatment of presymptomatic *TgSOD1^G93A/+^* mice with 25-hydroxycholesterol stabilizes NMJ lipid rafts (Zakyrjanova et al., 2021), supporting a protective role for cholesterols in MN survival.

In contrast to P120 *Sod1^G85R/G85R^* and *Sod1^-/-^* MNs, our transcriptomic analysis of P120 *TgSOD1^G93A/+^* MNs, in which neurodegenerative changes are well underway, show that cholesterol synthesis pathways are down-regulated, and genes involved in cholesterol clearance are up-regulated. One might speculate that neuroprotective signaling of cholesterol synthesis activation occurs in *TgSOD1^G93A/+^* MNs at earlier time points, possibly triggered by NMJ destabilization or distal axonopathy. If not efficiently used to stabilize the cell, accumulation of excess cholesterol and its toxic byproducts would suppress cholesterol synthesis in a cellular attempt to maintain cholesterol homeostasis.

Cholesterol cannot pass through membranes or the blood brain barrier. Control of cholesterol homeostasis within the CNS is separate from that of circulating cholesterol (Dietschy and Turley, 2004; Orth and Bellosta, 2012; Hartmann et al., 2021). Excess cholesterol can be cleared by enzymatic conversion (by enzymes such as Cyp27a1, Ch25h, Cyp46a1) to oxysterols that can either freely pass out of cells or activate cholesterol transporters and efflux pathways (ApoE, ABC transporters). We observe up-regulation of *Ch25h*, *Abca1* and *ApoE*, pointing to cholesterol clearance pathway activation, in *TgSOD1^G93A/+^* but not *Sod1^G85R/G85R^* or *Sod1^-/-^* MNs. Non-enzymatic cholesterol oxidation (auto-oxidation) also occurs. Oxysterol accumulation occurs in ALS patient and *TgSOD1^G93A/+^* mouse spinal cords (Dodge et al., 2021). Auto-oxidation-derived oxysterols are more toxic than enzymatically-derived oxysterols in human iPSC-MNs, implicating toxic auto-oxidized species in the pathogenesis of ALS under high cholesterol conditions (Dodge et al., 2021). Excess cholesterol can also be lowered by conversion to cholesterol esters by sterol O-acyltransferase 1 (SOAT1). We observe *Soat1* up-regulation in *TgSOD1^G93A/+^* but not *Sod1^G85R/G85R^* or *Sod1^-/-^* MNs. Cholesterol esters normally provide a form of cholesterol stored in lipid droplets for later use. However, cholesterol ester levels are elevated in ALS models under oxidative stress (Cutler et al., 2002; Chaves-Filho et al., 2019). Lysophosphatidylcholine (Lyso-PC), a by-product of cholesterol ester synthesis, is toxic to human iPSC-MNs, induces demyelination (Plemel et al., 2018), and is elevated in ALS patients and *TgSOD1^G93A/+^* spinal cords (Hanrieder and Ewing, 2014; Dodge et al., 2020). These findings support the view that elevated cholesterol in ALS MNs could induce accumulation of toxic cholesterol ester storage products or related lipid species that promote MN death. Differences in the ability of *Sod1^G85R/G85R^* and *Sod1^-/-^*, compared to *TgSOD1^G93A/+^* MNs, to maintain cholesterol homeostasis could contribute to differences in their survival.

In addition to cholesterol pathway genes, mRNAs encoding ion channels and transporters with a range of functions are expressed in strikingly similar, abnormal patterns in *Sod1^G85R/G85R^* and *Sod1^-/-^* MNs. Collectively, gene expression analyses of LCM isolated MNs and bulk ventral and dorsal spinal cord tissues demonstrate that *Sod1^G85R/G85R^* and *Sod1^-/-^* mice are very similar at the molecular level, and that very low-level mutant SOD1 expression in *Sod1^G85R/G85R^* mice phenocopies SOD1 loss-of-function in S*od1^-/-^* mice. *Sod1^G85R/G85R^* mice will be useful for determining mechanisms underlying selective vulnerability of some MNs and provide insight into pathways, potentially involving cholesterol homeostasis, that contribute to long-term mutant MN survival. Such information could lead to new strategies for ALS therapeutic intervention.

## DATA AVAILABILITY

All RNA sequencing data for samples described in this manuscript will be publicly available at the time of publication.

## AUTHOR CONTRIBUTIONS

J.A.D., A.J., J.R.F, D.L., R.H.B. designed the research.

J.A.D., L.A.M., J.P.W., K.L.R., H.Z., H.B., S.T., Y.Z., N.W., E.M.S., J.P., A.W., K.K., M.H.B. performed the research.

J.A.D., L.A.M., J.P.W., K.L.R., K.M.W., H.B., A.K., E.M.S., M.H.B., A.J., J.R.F, D.L., R.H.B. analyzed the data.

J.A.D. compiled the first draft of the paper.

J.A.D., L.A.M., K.M.W., A.J., J.R.F., D.L., R.H.B wrote and edited the paper.

## ACKNOWLEDGEMENTS

This research was supported by a grant from ALS Finding a Cure and the Leandro P. Rizzuto Foundation (R.H.B., J.F., D.L.), DEARS Foundation Brown University (A.J.), National Institute of Neurological Disorders and Stroke NS121977 (A.J.), NS055251 (D.L.), NS064295 and NS112743 (J.R.F.), National Institute on Aging AG073743 (J.R.F.), National Human Genome Research Institute U01 HG007910–01 and National Center for Advancing Translational Sciences UL1 TR001453–01 (A.K.), Carney Institute for Brain Science (D.L.), Brown University (A.J.), and visiting scholar support from West China Hospital and West China School of Medicine (H.Z., S.T., Y.Z).

## EXTENDED MEDIA LEGENDS

**Extended Video 5-1. 3D reconstruction of NMJs in TA muscle from a 3-month-old, male WT mouse.** Presynaptic axons and nerve terminals are stained with antibodies against TUBB3 and VAChT in red, and postsynaptic AChRs are visualized using green fluorescent α-bungarotoxin. NMJs are fully innervated and motor axons are continuous. Scale bar: 50 μm.

**Extended Video 5-2. 3D reconstruction of NMJs in TA muscle from a 3-month-old, male *SOD-1^G85R/G85R^* mouse.** Presynaptic axons and nerve terminals are stained with antibodies against TUBB3 and VAChT in red, and postsynaptic AChRs are visualized using green fluorescent α-bungarotoxin. NMJs are partially innervated, and some motor axons are fragmented or end in retraction bulbs. Scale bar: 50 μm.

## EXTENDED DATA FIGURES

**Figure 2-1.**
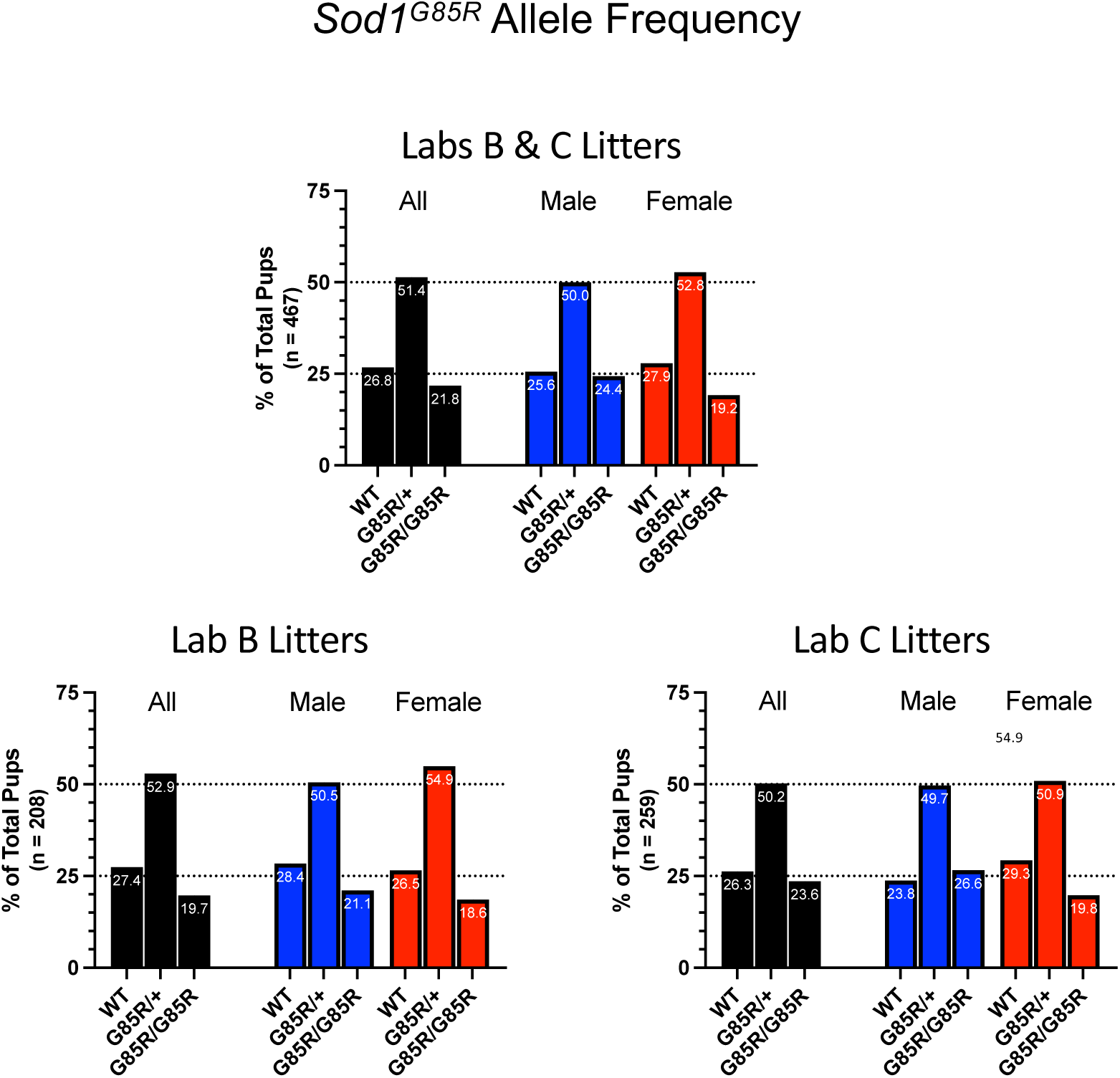
There is no difference in *Sod1^G85R^* allele frequency between males and females. Genotype frequencies of WT, *Sod1^G85R/+^* and *Sod1^G85R/G85R^* mutant mice compiled in two laboratories (Lab B and Lab C) conducting these studies show the expected 1:2:1 ratio for both males and females.

**Figure 2-2.**
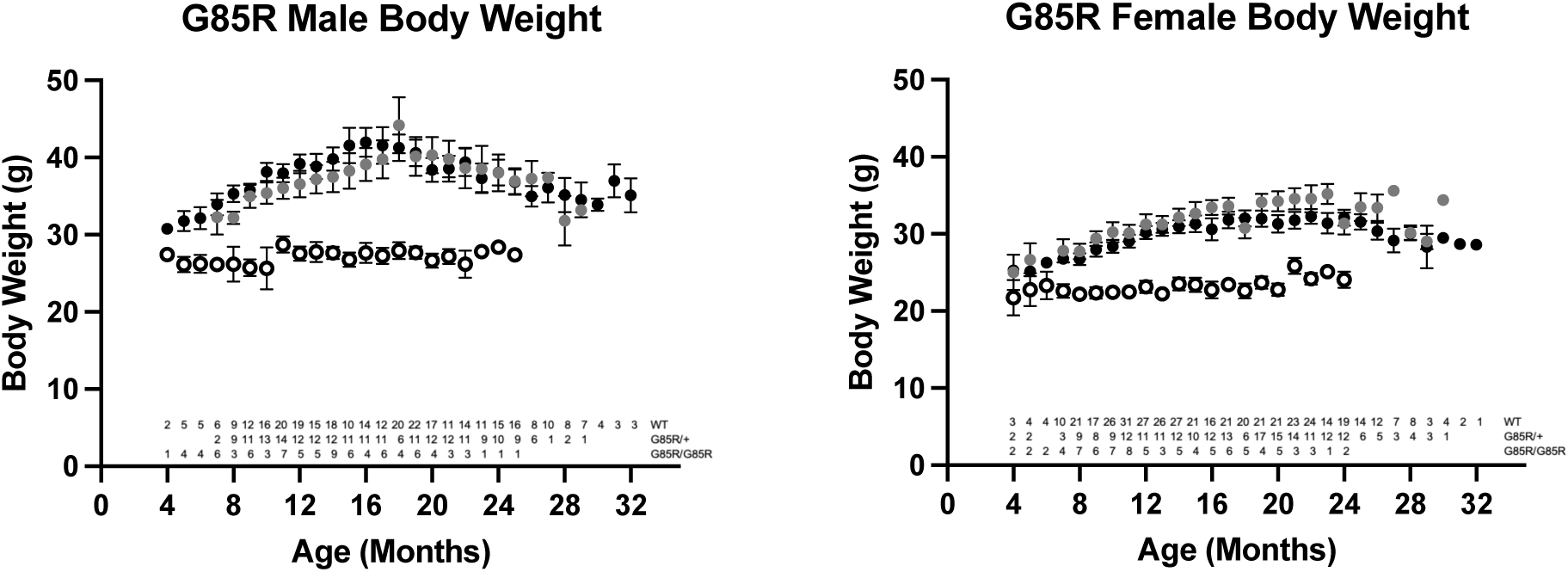
The body weight of *Sod1^G85R/G85R^* mice is lower that WT or *Sod1^G85R/+^* mice throughout their lifespan. Body weights of groups of mice monitored at different ages through 32-months of age. Solid black circles: WT, solid gray circles: *Sod1^G85R/+^*, open black circles: *Sod1^G85R/G85R^* (mean ± SE, numbers at bottom are the n at each point for each genotype).

**Figure 3-1.**
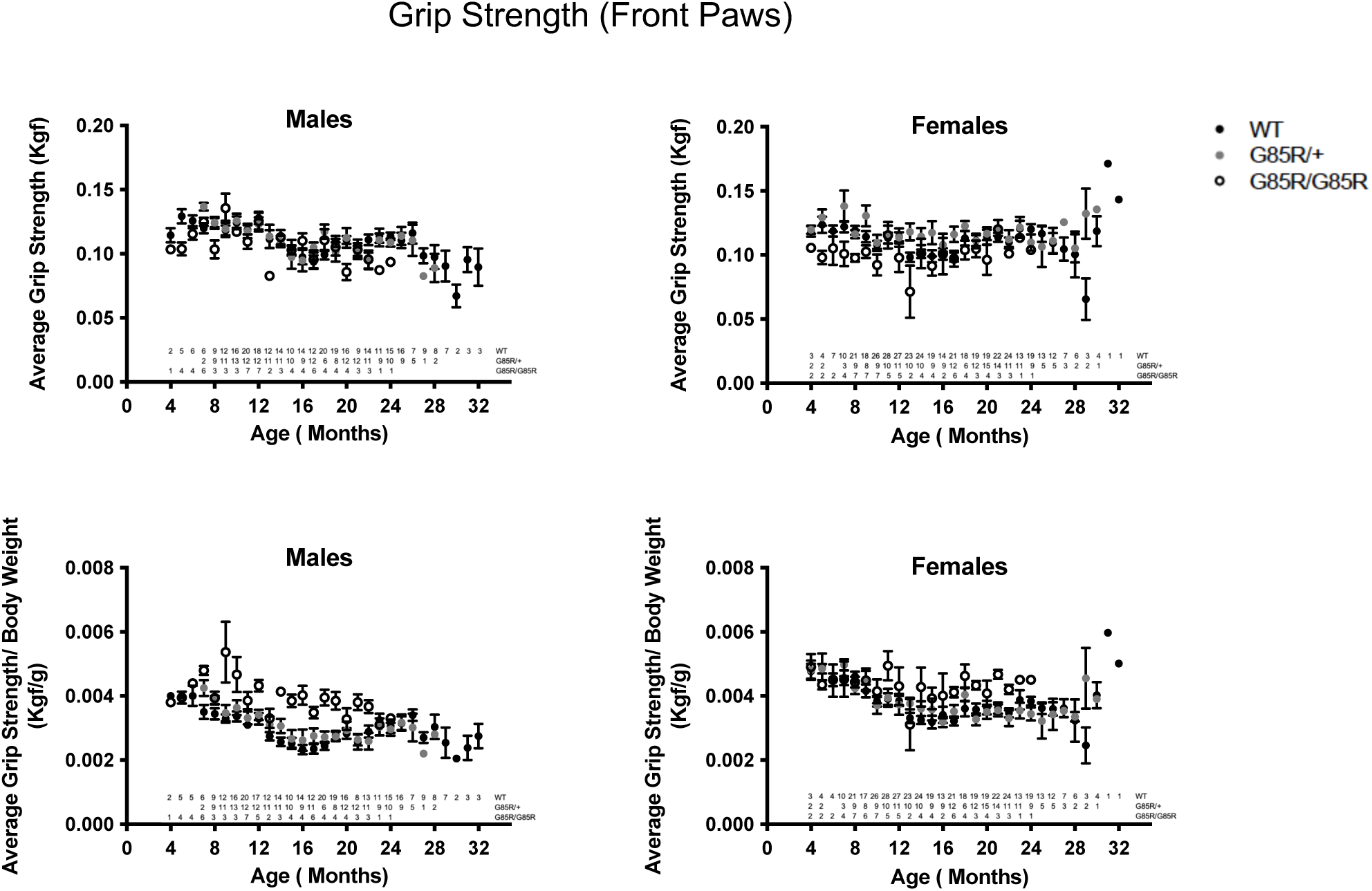
There is no consistent difference in normalized front paw grip strength in *Sod1^G85R/G85R^*, *Sod1^G85R/+^* or WT mice throughout their lifespan. Front paw grip strength measurements of groups of mice monitored at different ages through 32-months of age. Solid black circles: WT, solid gray circles: *Sod1^G85R/+^*, open black circles: *Sod1^G85R/G85R^*. Top panels are raw grip strength and lower panels are grip strength normalized to body weight. (mean ± SE, numbers at bottom are the n at each point for each genotype.)

**Figure 3-2.**
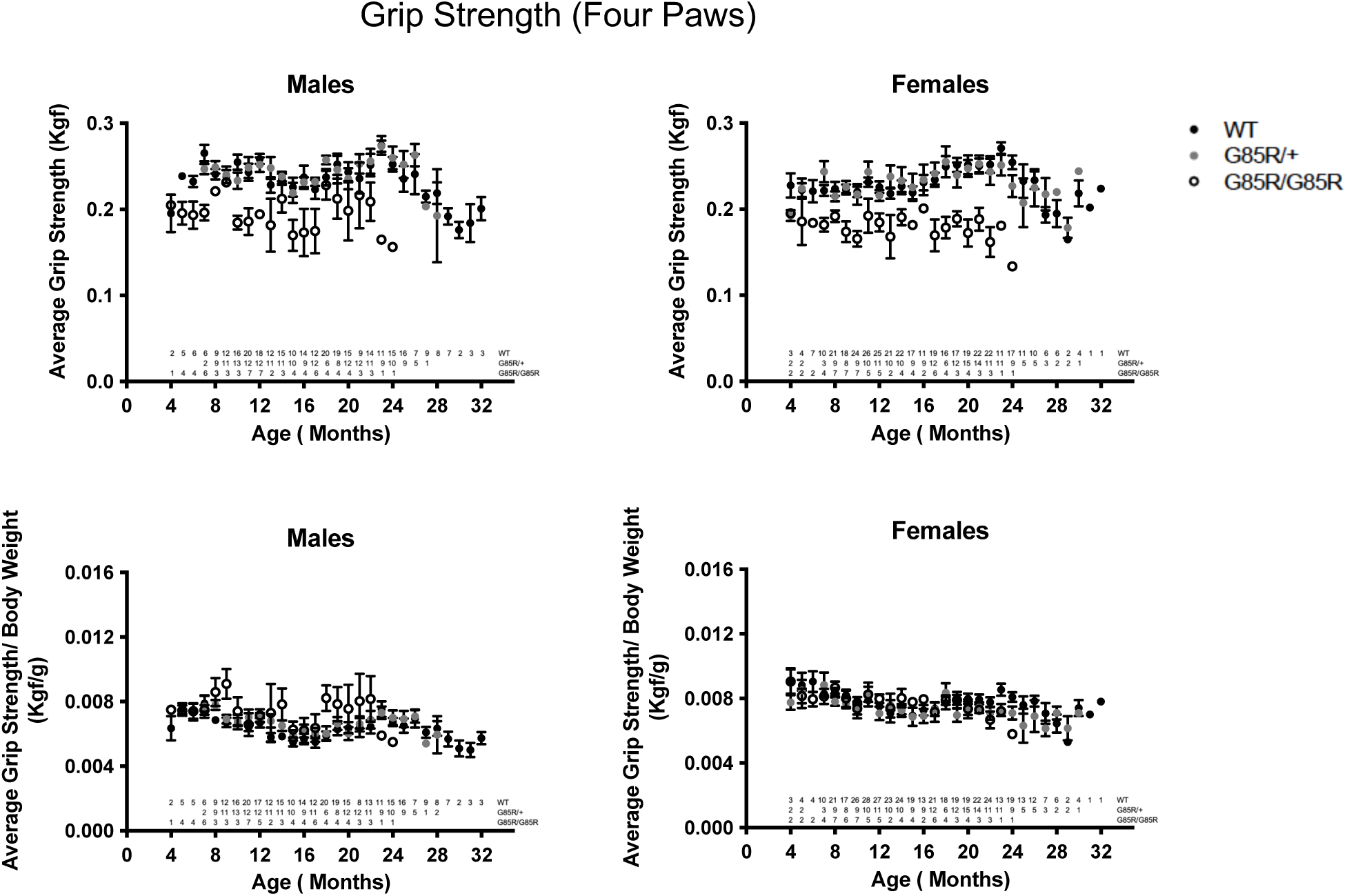
There is no consistent difference in normalized four paw grip strength in *Sod1^G85R/G85R^*, *Sod1^G85R/+^* or WT mice throughout their lifespan. Four paw grip strength measurements of groups of mice monitored at different ages through 32-months of age. Solid black circles: WT, solid gray circles: *Sod1^G85R/+^*, open black circles: *Sod1^G85R/G85R^*. Top panels are raw grip strength and lower panels are grip strength normalized to body weight. (mean ± SE, numbers at bottom are the n at each point for each genotype.)

**Figure 5-1.**
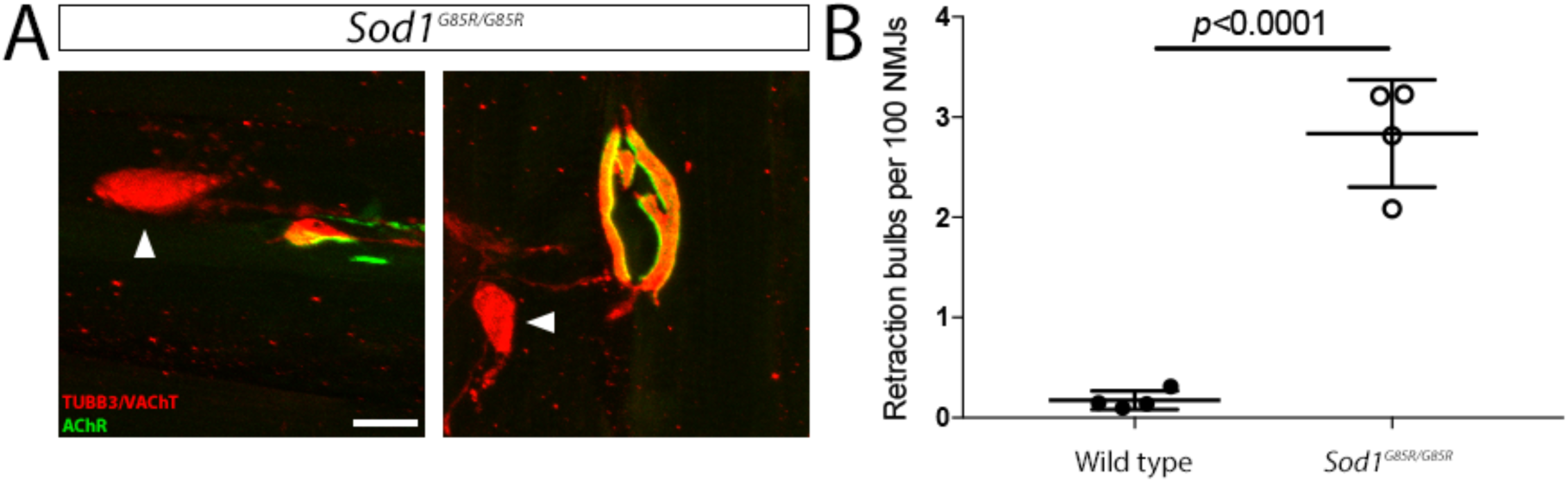
Retraction bulbs are observed in *Sod1^G85R/G85R^* mice. **(A)** Longitudinal sections of TA muscles from 3-month-old, male WT (not shown) and *Sod1^G85R/G85R^* mice (two examples) were stained with antibodies against TUBB3 and VAChT to visualize motor axons and nerve terminals, respectively. Postsynaptic AChRs were visualized using fluorescent α-bungarotoxin. Retraction bulbs (arrowheads) are frequently observed in *Sod1^G85R/G85R^*, but not WT mice. **(B)** Quantification of retraction bulbs normalized to the number of NMJs in 3-month-old males shows significantly increased incidence of retraction bulbs in *Sod1^G85R/G85R^* mice, as compared to WT (n = 4 animals per genotype). Scale bar: 10 μm. Error bars indicate SEM.

**Figure 7-1.**
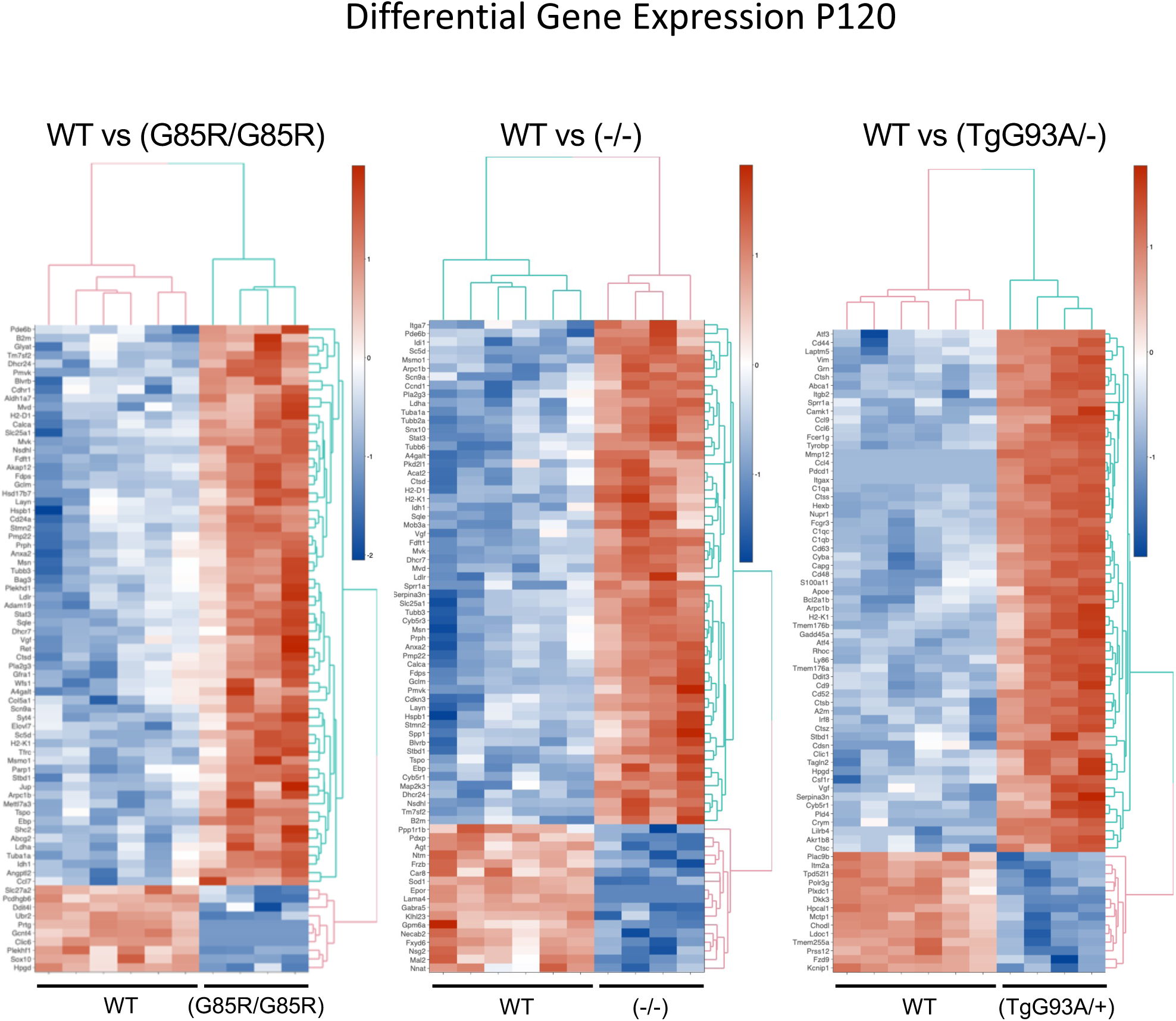
Top differentially expressed genes in *Sod1* mutants compared to WT at P120. Hierarchical clustering heat maps of the top 75 most differentially expressed genes at day P120 in LCM motor neurons of WT compared to homozygous *Sod1^G85R/G85R^*, *Sod1^-/-^*, or heterozygous *TgSOD1^G93A/+^* mutants (n = 6 WT mice; n= 4 of each mutant type). At this age, in each comparison, there are more upregulated genes in the mutants (red shades) than downregulated genes (blue shades).

**Figure 7-2.**
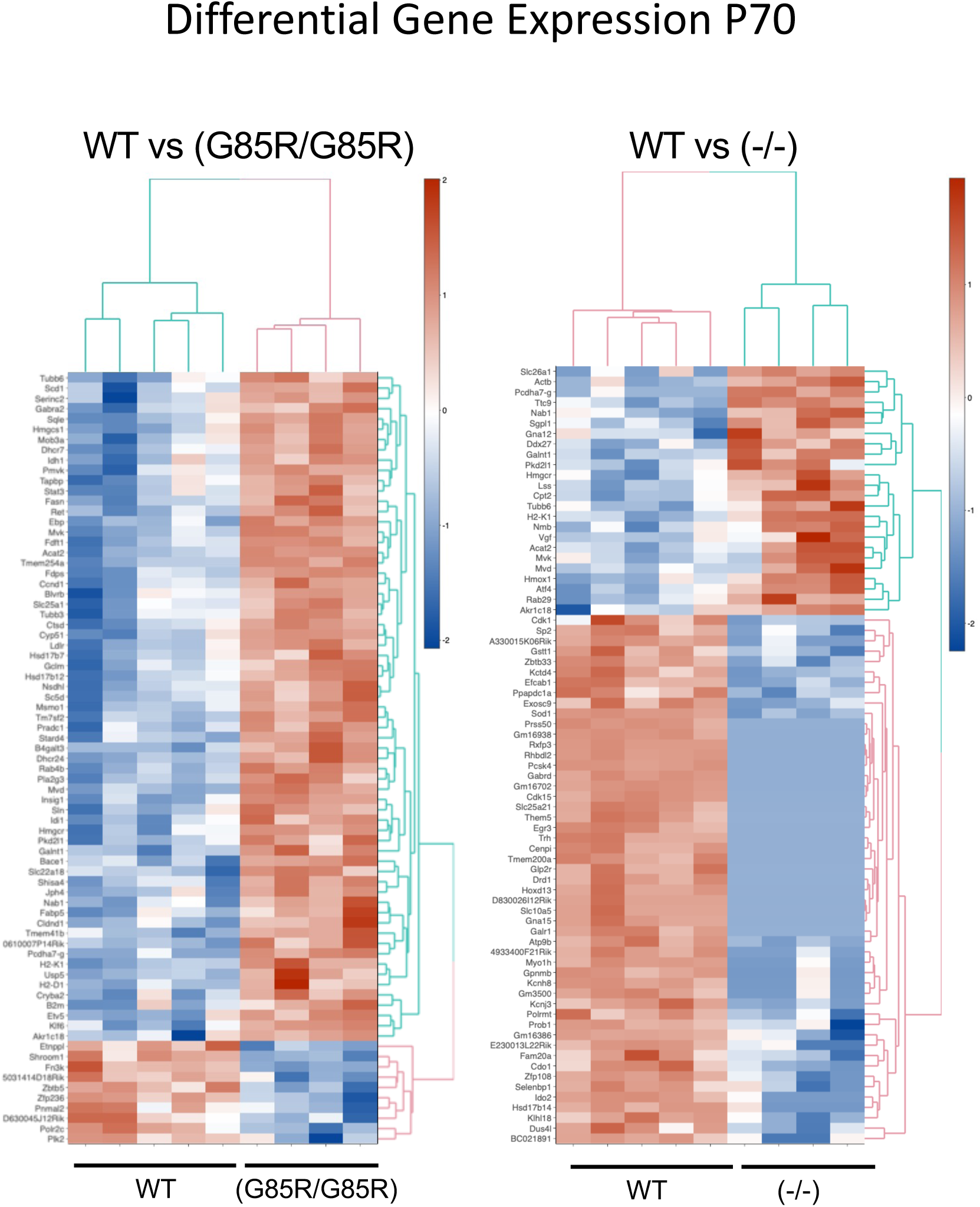
Top differentially expressed genes in *Sod1* mutants compared to WT at P70. Hierarchical clustering heat maps of the top 75 most differentially expressed genes at day P70 in LCM motor neurons of WT compared to homozygous *Sod1^G85R/G85R^* or *Sod1^-/-^* (n = 5 WT mice; n= 4 mutant). Upregulated genes in red shades, downregulated genes in blue shades.

**Figure 7-3.**
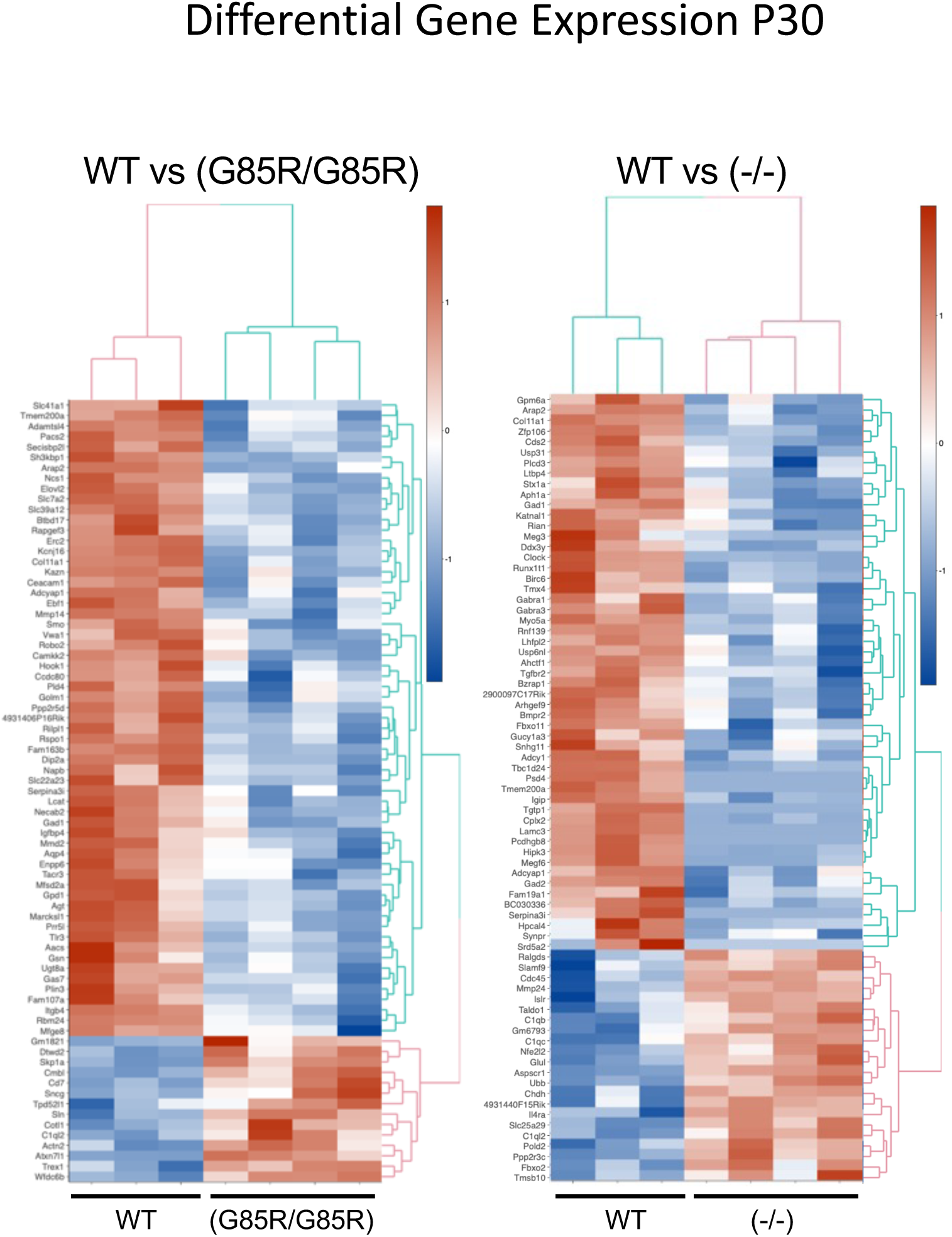
Top differentially expressed genes in *Sod1* mutants compared to WT at P30. Hierarchical clustering heat maps of the top 75 most differentially expressed genes at day P30 in LCM motor neurons of WT compared to homozygous *Sod1^G85R/G85R^* or *Sod1^-/-^* (n = 3 WT mice; n= 4 mutant). Upregulated genes in red shades, downregulated genes in blue shades.

**Figure 7-4.**
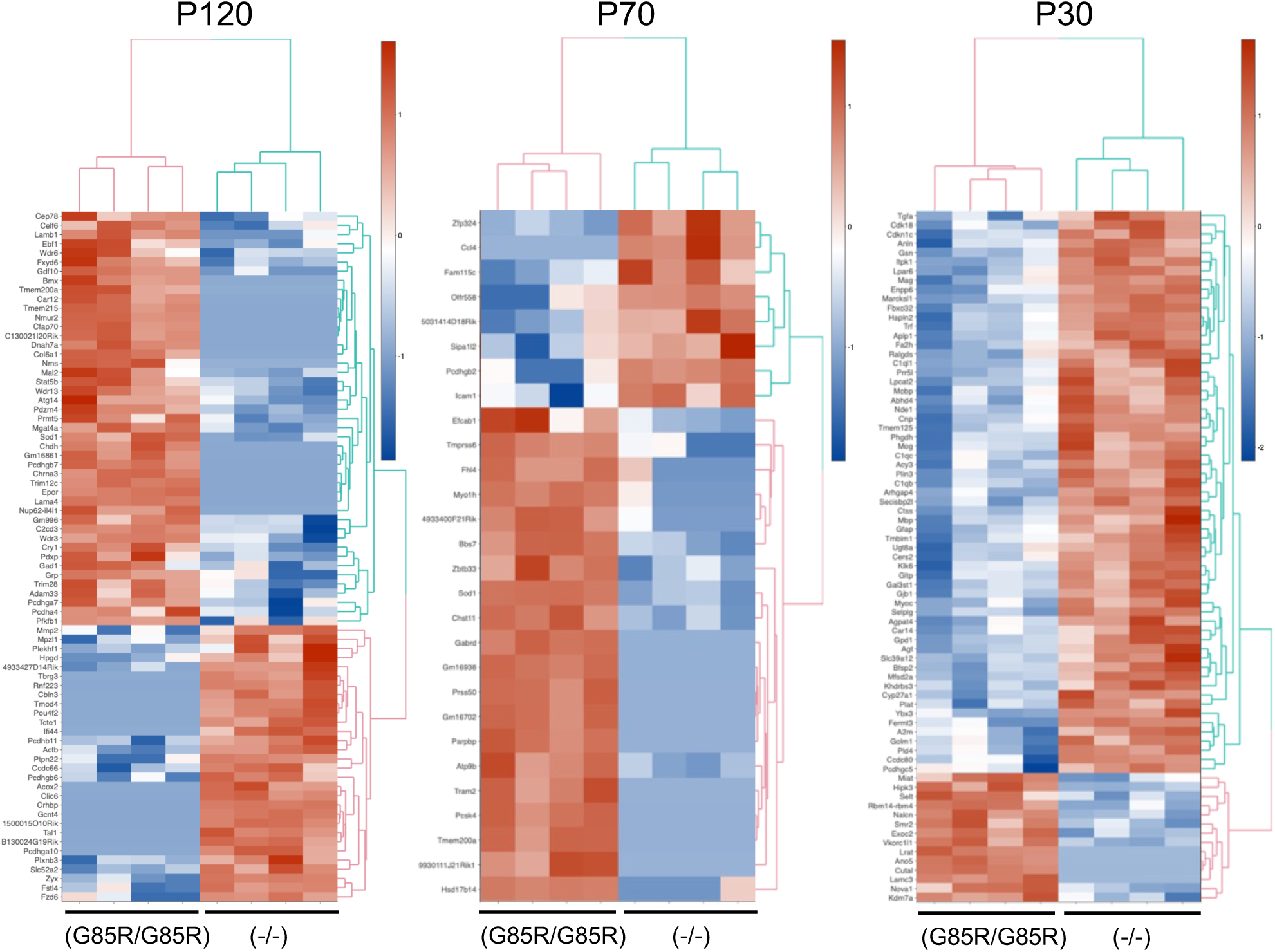
Top differentially expressed genes in *Sod1^G85R/G85R^* mutants compared to *Sod1^-/-^* mutants. Hierarchical clustering heat maps of the top differentially expressed genes at each age (day P120, P70, P30) in LCM Motor neurons of *Sod1^G85R/G85R^* compared to *Sod1^-/-^* mice (n= 4 of each mutant type).

**Figure 8-1.**
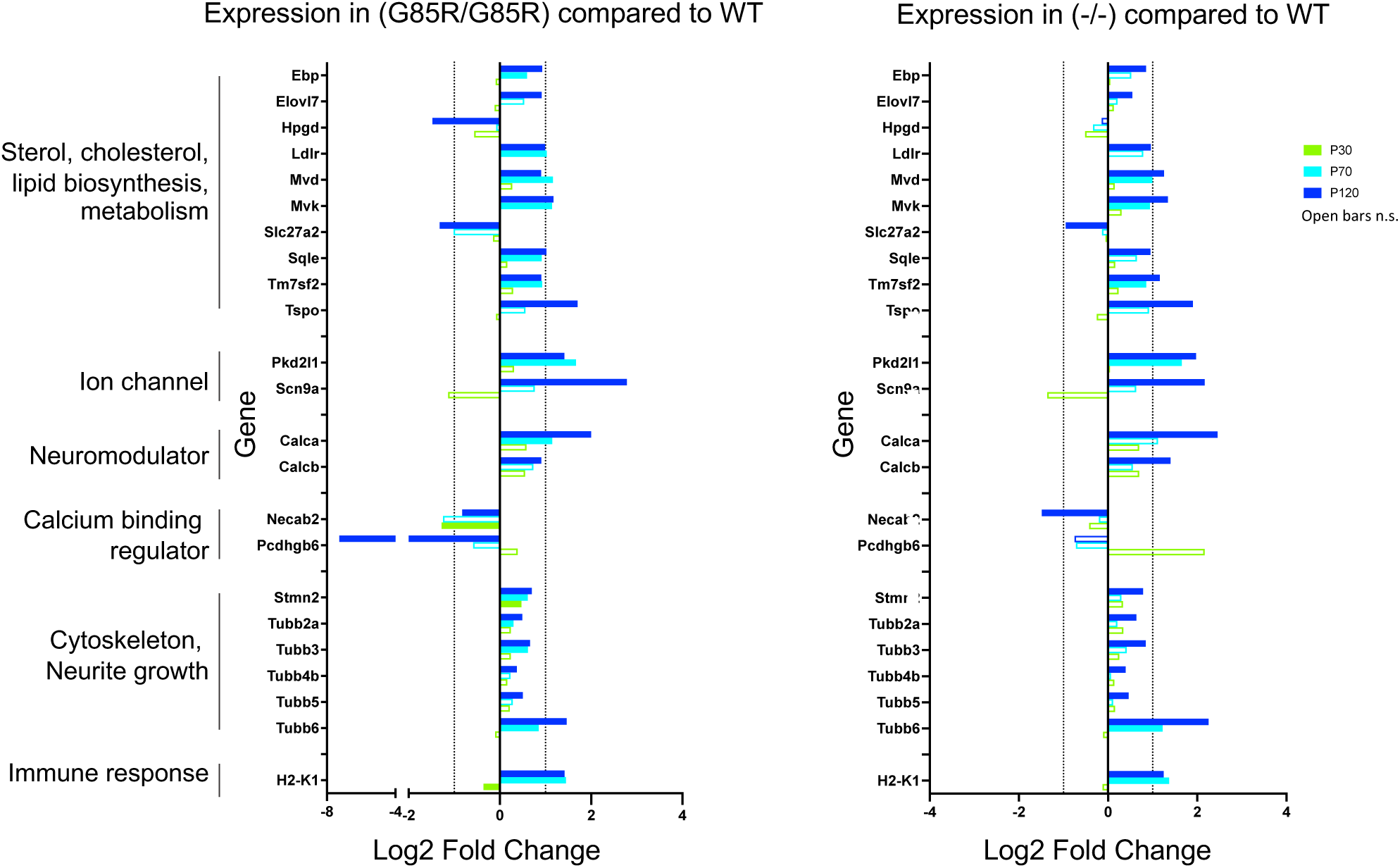
*Sod1^G85R/G85R^* and *Sod1^-/-^* motor neurons have common changes in gene expression at both day P70 and P120. Several genes that are differentially expressed at P120 in *Sod1^G85R/G85R^* vs WT or *Sod1^-/-^* vs WT Motor neurons are also differentially expressed at P70. There is almost no significant differential expression of these genes at P30. Solid bars show statistically significant expression differences at P120 (dark blue), P70 (light blue) and P30 (green) (padj < 0.05). Open bars: not significant.

**Figure 9-1.**
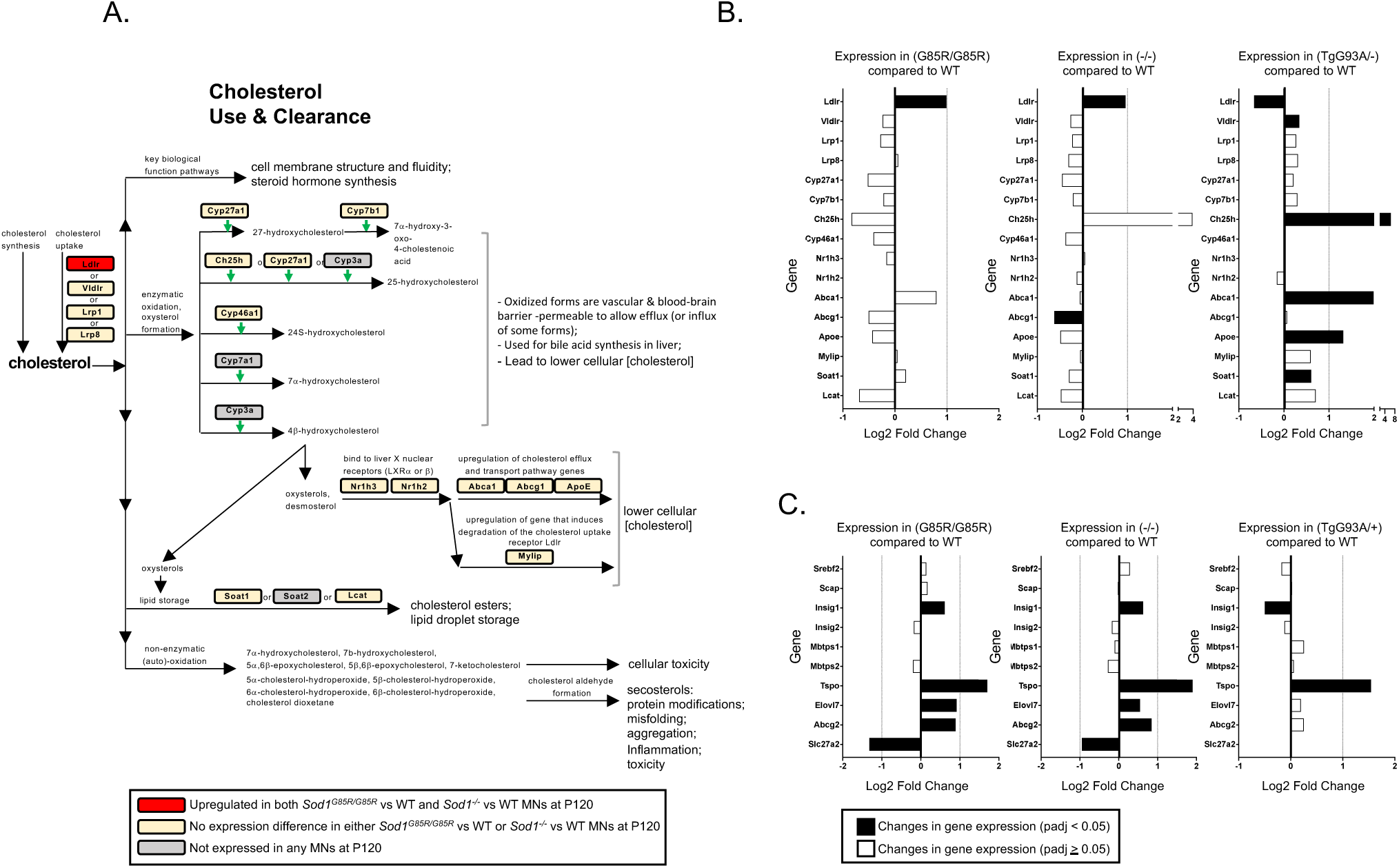
*Sod1^G85R/G85R^* and *Sod1^-/-^* motor neurons do not upregulate genes involved cholesterol clearance and have gene expression profiles very different from *TgSOD1^G93A/+^* motor neurons. **(A)** Genes for proteins (in boxes) involved in various cholesterol use and clearance pathways and metabolites are shown (genes highlighted in red are upregulated at P120 in both *Sod1^G85R/G85R^* vs WT and *Sod1^-/-^* vs WT motor neurons; yellow boxes: no significant difference in expression in mutant vs WT; gray boxes: not expressed). **(B)** Only *Ldlr*, involved in cholesterol uptake, is upregulated in both *Sod1^G85R/G85R^* vs WT and *Sod1^-/-^* vs WT motor neurons at P120, while it is downregulated in *TgSOD1^G93A/+^* vs WT (black bars: significant expression differences (padj < 0.05); white bars: differences not significant (padj ≥0.05)). Several genes involved in cholesterol clearance are upregulated in *TgSOD1^G93A/+^* vs WT motor neurons while an opposite trend, though not reaching significance, is observed in both *Sod1^G85R/G85R^* vs WT and *Sod1^-/-^* vs WT motor neurons. **(C)** Other genes involved in cholesterol regulation and fatty acid metabolic pathways have similar differential expression patterns in *Sod1^G85R/G85R^* vs WT and *Sod1^-/-^* vs WT motor neurons at P120 that are distinct from *TgSOD1^G93A/+^* vs WT motor neurons (black bars padj < 0.05; white bars padj ≥0.05).

**Figure 11-1.**
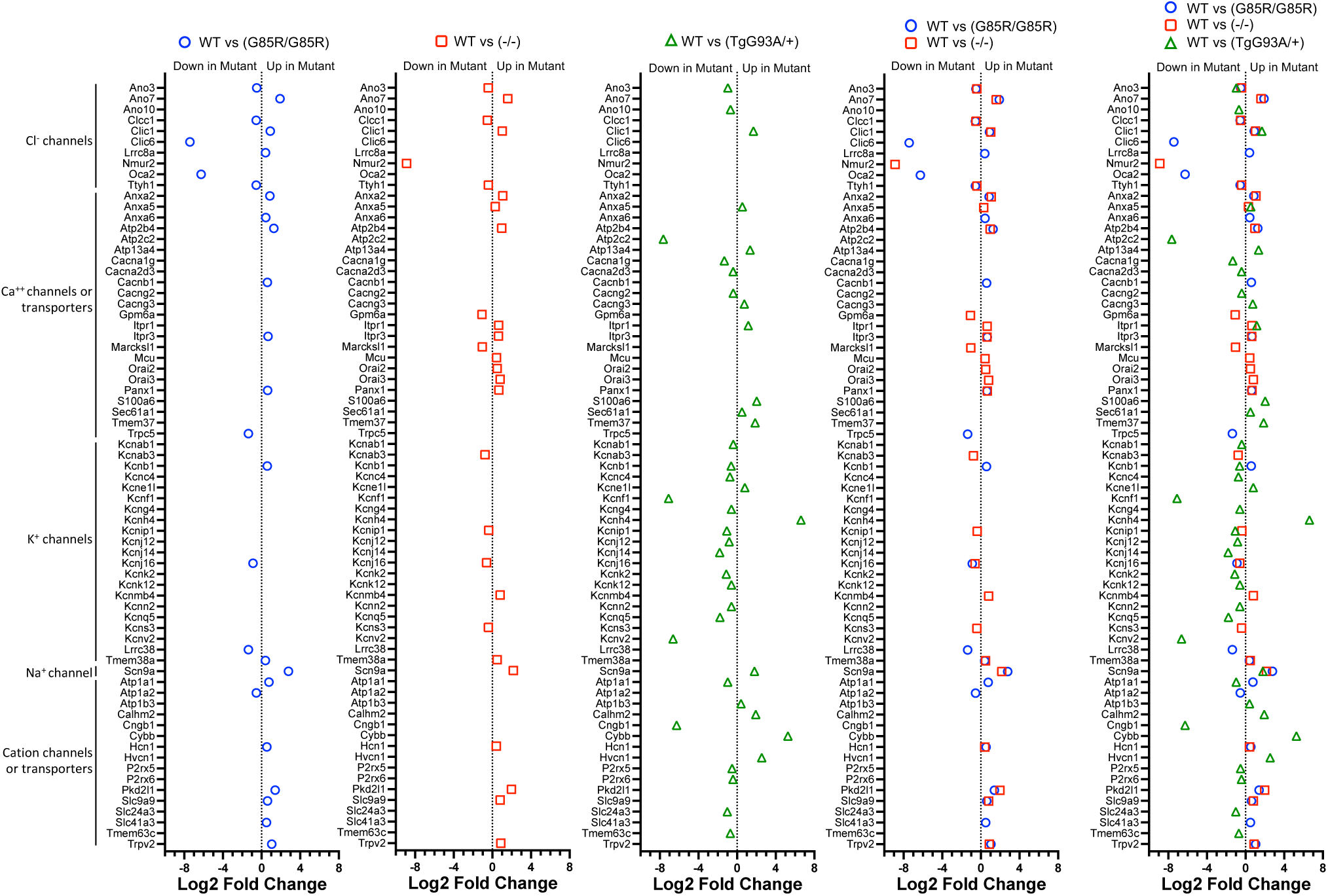
*Sod1^G85R/G85R^*, *Sod1^-/-^* and *TgSOD1^G93A/+^* mutant motor neurons differentially express ion channel, transporter and neurotransmitter genes compared to WT motor neurons. All 148 genes depicted in the Figure 11A Venn diagram of differentially expressed ion channel, transporter or neurotransmitter genes are listed here or in Figure 11-2, along with their level of differential expression in the pairwise comparisons shown in each panel (padj <0.05).

**Figure 11-2.**
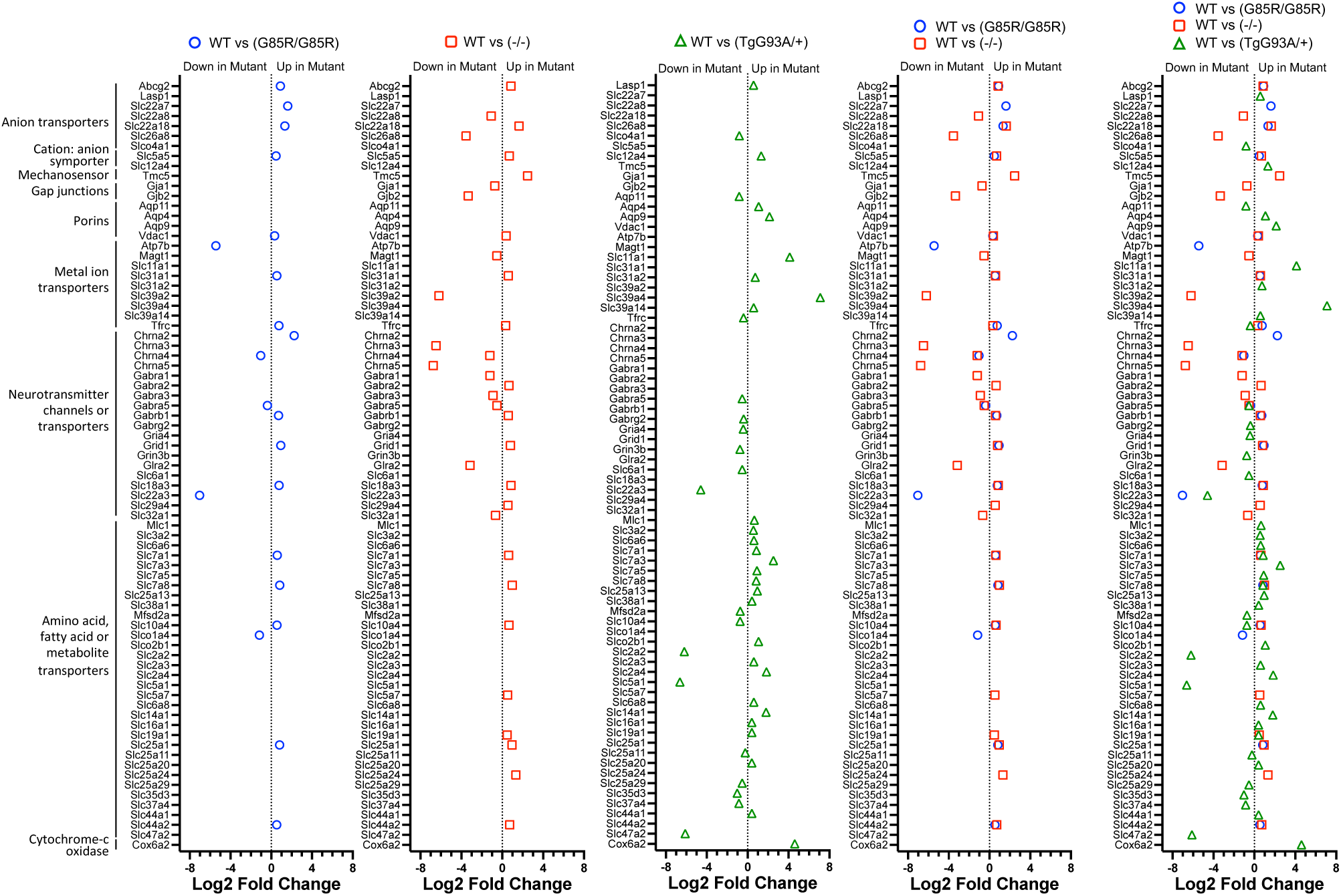
*Sod1^G85R/G85R^*, *Sod1^-/-^* and *TgSOD1^G93A/+^* mutant motor neurons differentially express ion channel, transporter and neurotransmitter genes compared to WT motor neurons. Continuation, from Figure 11-1, of the listing of all 148 genes depicted in the Figure 11A Venn diagram of differentially expressed ion channel, transporter, or neurotransmitter genes, along with their level of differential expression in the pairwise comparisons shown in each panel (padj <0.05).

**Figure 11-3.**
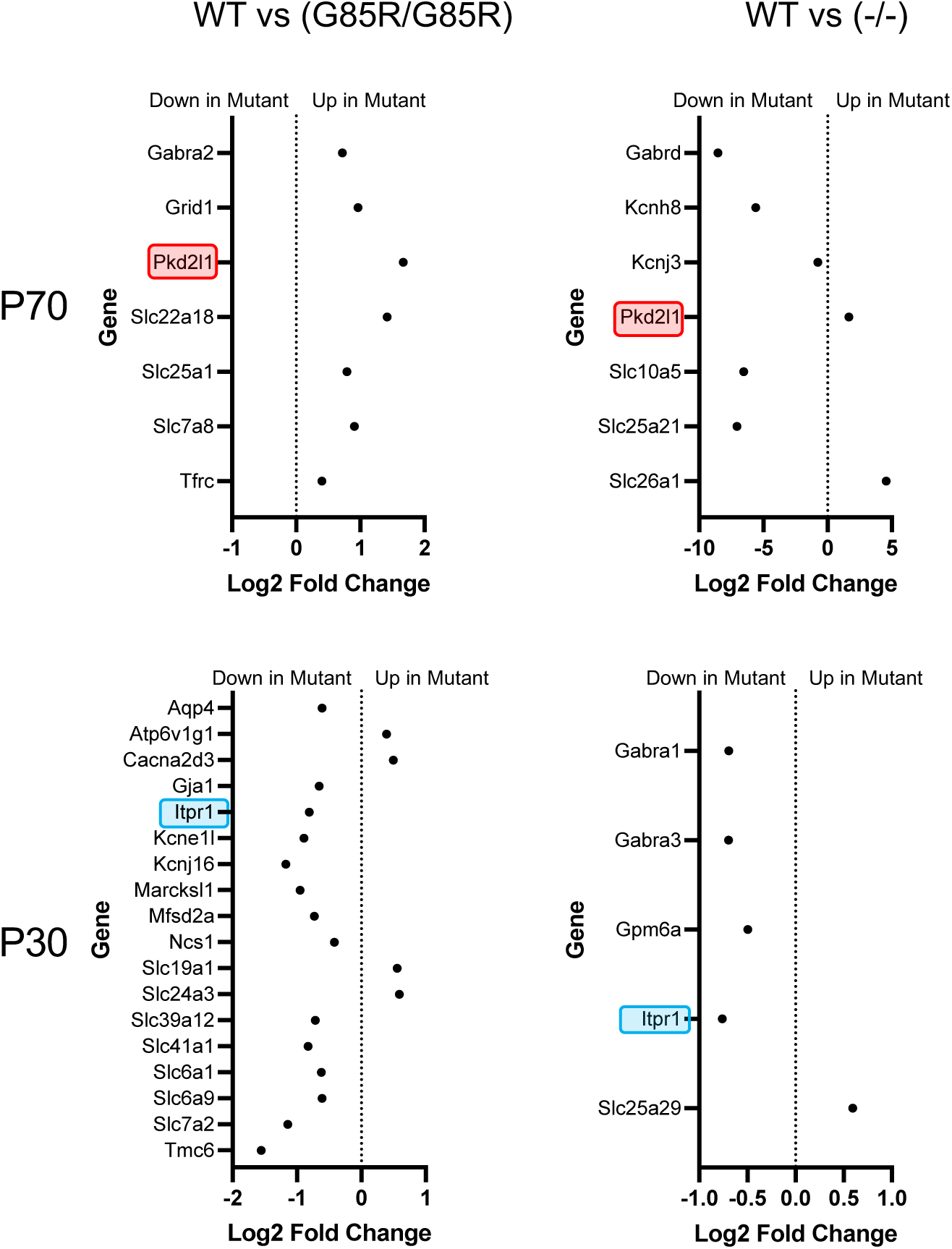
*Sod1^G85R/G85R^* and *Sod1^-/-^* motor neurons have very few common changes in ion channel or transporter gene expression at P70 or P30. We examined the expression of a set of 877 genes associated with ion channel, transmembrane transporter, and neurotransmitter Gene Ontology terms to determine which are differentially expression in the LCM motor neurons from mutant vs WT mice at P70 and P30. Of these 877 genes, all that are differentially expressed in WT vs *Sod1^G85R/G85R^*, WT vs *Sod1^-/-^* or *Sod1^G85R/G85R^* vs *Sod1^-/-^* are shown (padj < 0.05). Blue and red boxes highlight the only genes commonly changing in expression in the mutants.

**Figure 12-1.**
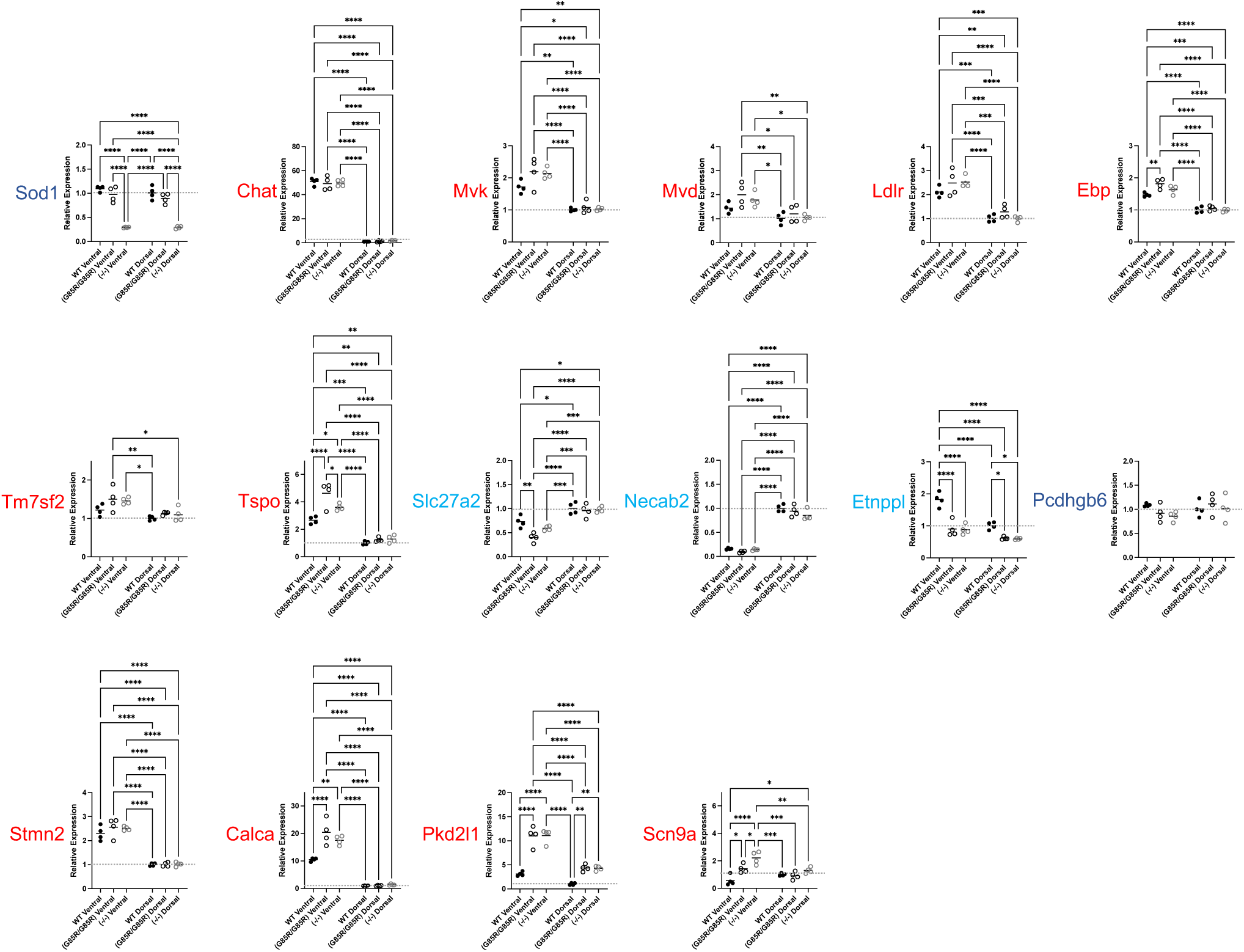
Ventral and dorsal spinal cord tissues exhibit differential expression of cholesterol pathway and other genes, some of which are significantly altered in *Sod1^G85R/G85R^* and *Sod1^-/-^* mutants. Data shown is the same data as in Figure 12 but with all the statistically significant differences indicated by brackets (Mean ± SEM; ANOVA *p < 0.05, **p < 0.005, ***p < 0.0005, ****p<0.0001).

**Table 1-1.**
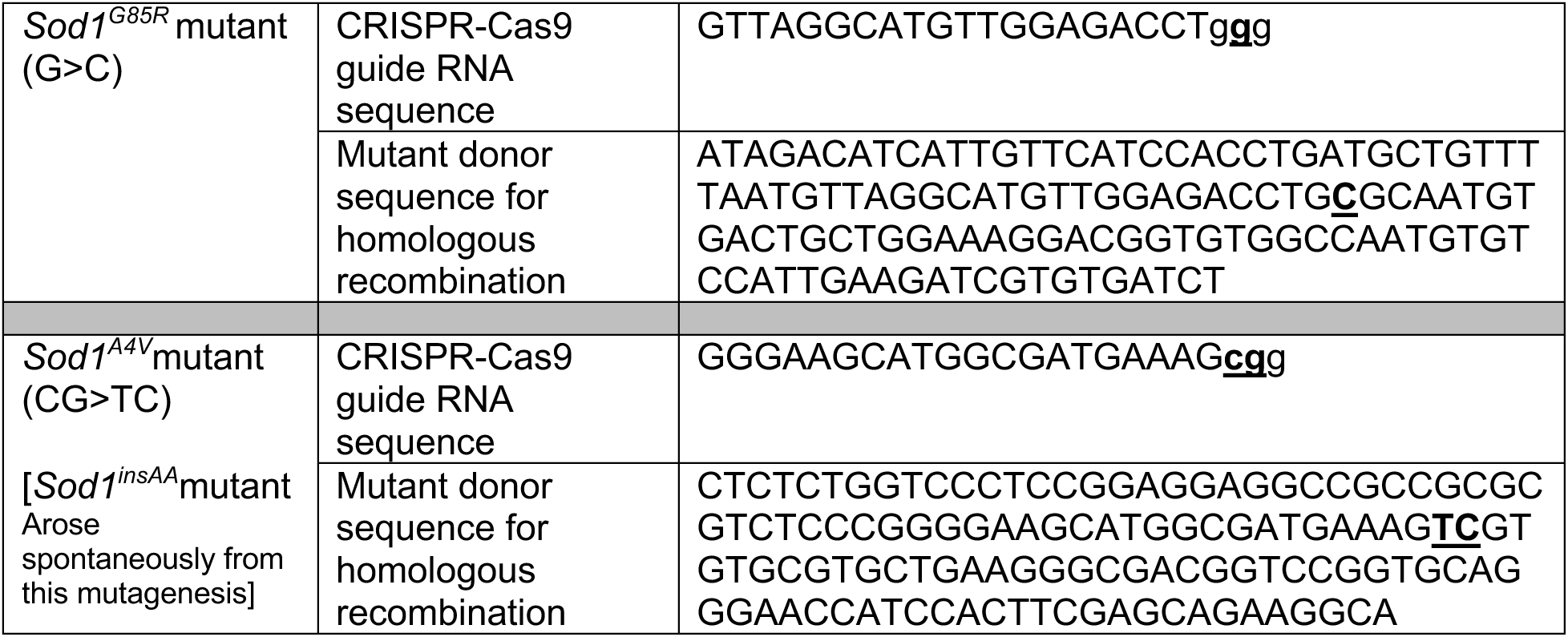
Sequences Used to Create *Sod1* Mutant Mice. Altered bases in the donor sequences to create desired mutations are shown in bold, underlined. Corresponding wild type bases in guide RNA are also in bold, underlined. The PAM sequences in the guide RNAs are in lower case.

**Table 1-2.**
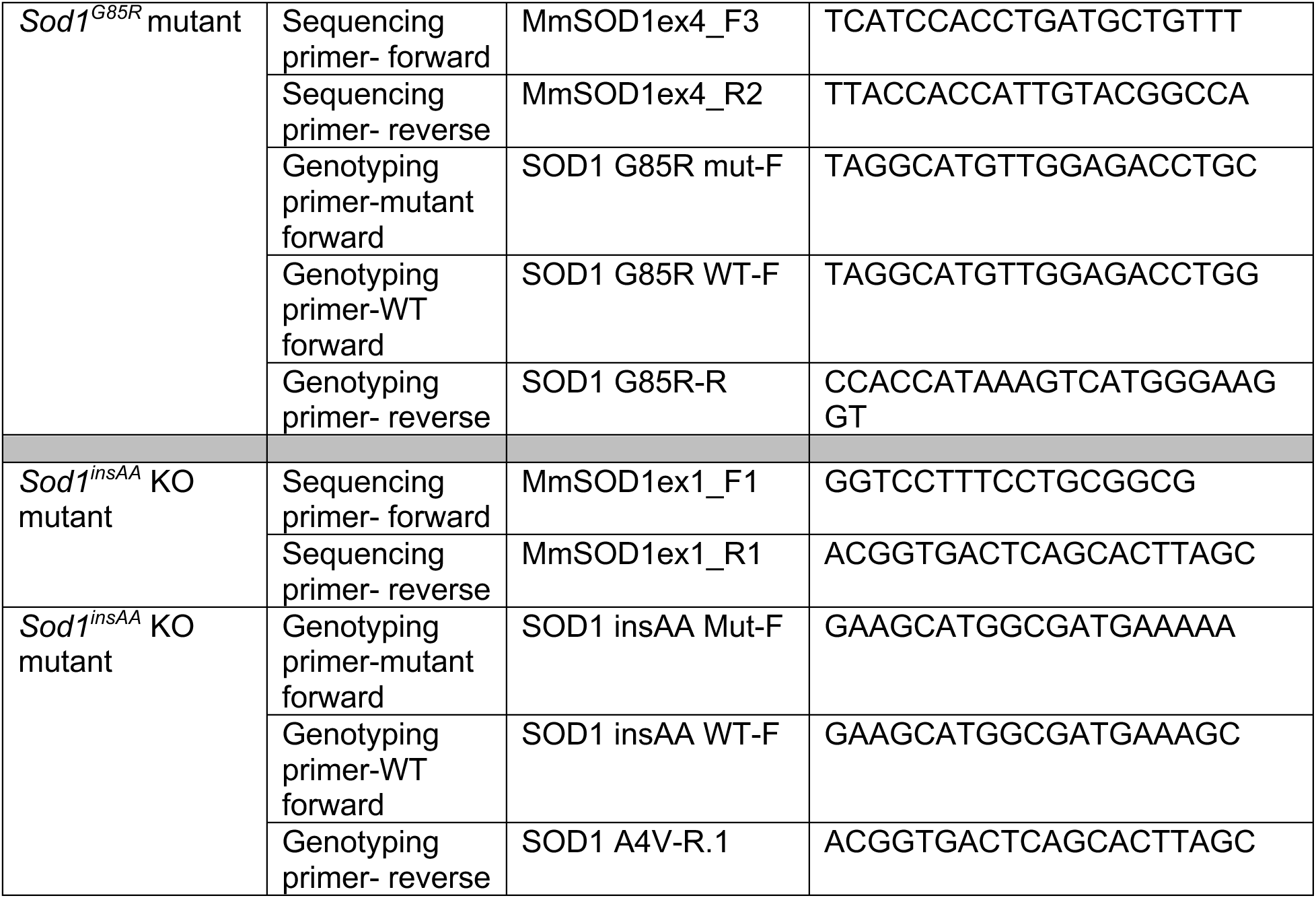
Primers Used to Sequence and Genotype *Sod1* Mutant Mice.

**Table 4-1.** Electrophysiology Analyses of Motor Neurons Innervating TA and Soleus Muscles. Averages +/- standard errors are shown for (N) number of motor neurons. The electrophysiological properties of MNs in the ventral horn of the spinal cord of *Sod1^G85R/G85R^* and WT littermates that project to either TA or soleus muscles are shown for mice at 1 and 3 months of age. Only muscle projecting neurons were analyzed (contained Evans blue dye, see Methods). Multiple electrophysiological parameters are shown including capacitance as a measure of cell size (pF), resting membrane potential (RMP, mV), input resistance (Mn), rheobase current required to trigger an action potential (pA), firing rate in response to current injection (Hz), action potential peak (mV), action potential afterhyperpolarization (AHP) peak (mV) and half width (ms). *Sod1^G85R/G85R^* and WT neurons were not distinguishable functionally across all parameters measured using multifactor ANOVA analyses. There was a significant difference in the average threshold of current injection needed to trigger an action potential (rheobase) based on the type of neuron. The rheobase which was significantly higher in neurons projecting to the TA compared to those projecting to the soleus (p < 0.001).

| TA Muscle | Capacitance (pF) | Input Resis. (MΩ) | RMP (mV) | Rheobase (pA) | Firing rate (Hz) | Threshold (mV) | AP peak (mV) | AHP peak (mV) | AHP half-width (ms) |
| --- | --- | --- | --- | --- | --- | --- | --- | --- | --- |
| Sod1<sup>+/+</sup> 1 mo. | 168.70 ± 6.2 (10) | 109.50 ± 9.6 (10) | -71.19 ± 0.9 (10) | 558.50 ± 21.6 (10) | 199.70 ± 27.0 (10) | -50.69 ± 1.9 (10) | 79.11 ± 3.2 (10) | -8.45 ± 0.3 (10) | 0.75 ± 0.0 (10) |
| Sod1<sup>G85R/G85R</sup> 1 mo. | 160.98 ± 10.4 (9) | 105.56 ± 11.6 (10) | -69.65 ± 1.0 (10) | 580.00 ± 39.8 (10) | 201.9 ± 17.5 (10) | -52.37 ± 1.5 (9) | 80.36 ± 1.8 (9) | -7.82 ± 0.3 (10) | 0.79 ± 0.0 (10) |
| Sod1<sup>+/+</sup> 3 mo. | 198.69 ± 7.1 (15) | 115.87 ± 7.7 (15) | -71.08 ± 1.1 (15) | 607.33 ± 16.4 (15) | 198.33 ± 17.1 (15) | -50.34 ± 1.2 (15) | 79.59 ± 3.2 (15) | -7.85 ± 0.3 (15) | 0.79 ± 0.0 (15) |
| Sod1<sup>G85R/G85R</sup> 3 mo. | 154.22 ± 10.6 (9) | 107.64 ± 11.2 (9) | -70.77 ± 1.1 (9) | 621.11 ± 38.0 (9) | 194.00 ± 16.1 (9) | -53.54 ± 1.3 (8) | 81.62 ± 2.3 (9) | -8.66 ± 0.3 (8) | 0.75 ± 0.0 (9) |
| Soleus Muscle | Capacitance (pF) | Input Resis. (MΩ) | RMP (mV) | Rheobase (pA) | Firing rate (Hz) | Threshold (mV) | AP peak (mV) | AHP peak (mV) | AHP half-width (ms) |
| Sod1<sup>+/+</sup> 1 mo. | 151.59 ± 7.5 (9) | 120.92 ± 9.8 (10) | -71.70 ± 0.8 (10) | 240.00 ± 35.3 (10) | 187.67 ± 14.7 (9) | -51.77 ± 1.9 (10) | 78.98 ± 2.1 (10) | -8.19 ± 0.4 (10) | 0.74 ± 0.0 (10) |
| Sod1<sup>G85R/G85R</sup> 1 mo. | 160.79 ± 10.0 (9) | 122.12 ± 8.1 (9) | -72.18 ± 1.2 (9) | 226.67 ± 22.2 (9) | 199.33 ± 9.5 (9) | -52.12 ± 1.5 (9) | 78.13 ± 2.6 (9) | -8.19 ± 0.6 (9) | 0.78 ± 0.0 (9) |
| Sod1<sup>+/+</sup> 3 mo. | 175.99 ± 10.8 (15) | 117.13 ± 5.2 (14) | -71.70 ± 0.8 (15) | 210.71 ± 12.5 (14) | 194.73 ± 20.6 (15) | -50.15 ± 1.9 (15) | 78.78 ± 2.3 (15) | -8.35 ± 0.3 (15) | 0.76 ± 0.0 (15) |
| Sod1<sup>G85R/G85R</sup> 3 mo. | 178.55 ± 20.3 (10) | 114.76 ± 13.0 (10) | -71.45 ± 0.9 (10) | 209.00 ± 20.4 (9) | 194.64 ± 18.0 (10) | -51.82 ± 1.0 (10) | 77.31 ± 2.5 (10) | -7.93 ± 0.3 (10) | 0.74 ± 0.0 (10) |

**Table 7-1.**
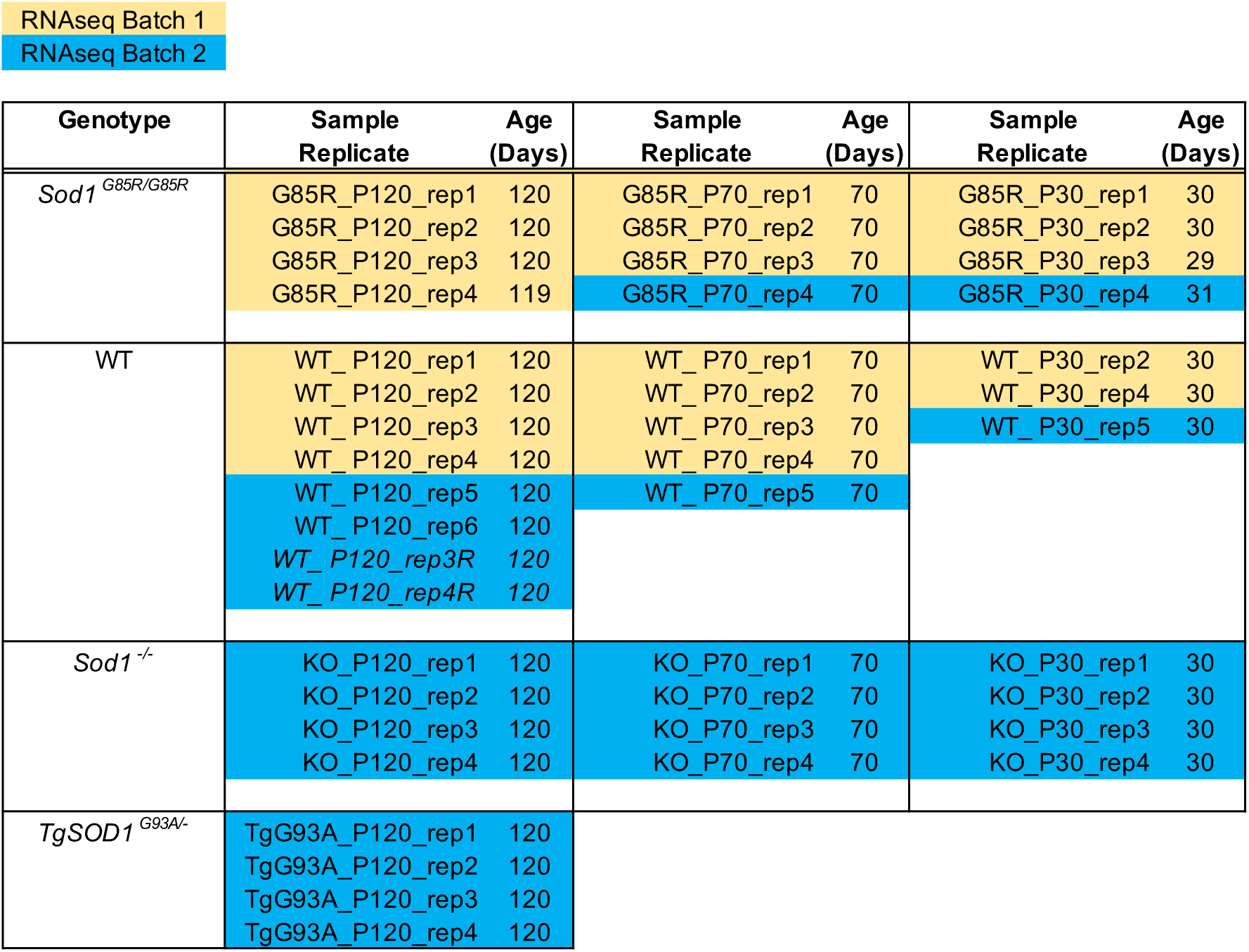
LCM Motor Neuron Samples Used in Transcriptome Analyses. WT_ P120_rep3R RNA sample is the exact same sample of RNA as Batch 1 WT_ P120_rep3 reprocessed in RNAseq Batch 2. WT_ P120_rep4R RNA sample is the exact same sample of RNA as Batch 1 WT_ P120_rep4 reprocessed in RNAseq Batch 2 WT_ P120_rep3R and WT_ P120_rep4R sample data was used in P120 Combat batch effect correction but not in differential gene expression analyses. Separate differential expression anaysis after Combat batch correction performed for quality control purposes found no differences in gene expression between WT_ P120_rep3 + WT_ P120_rep4 and WT_ P120_rep3R + WT_ P120_rep4R. For WT_P30 analyses, rep1 and rep3 samples were omitted due to unsuitable sample quality.

**Table 12-1.**
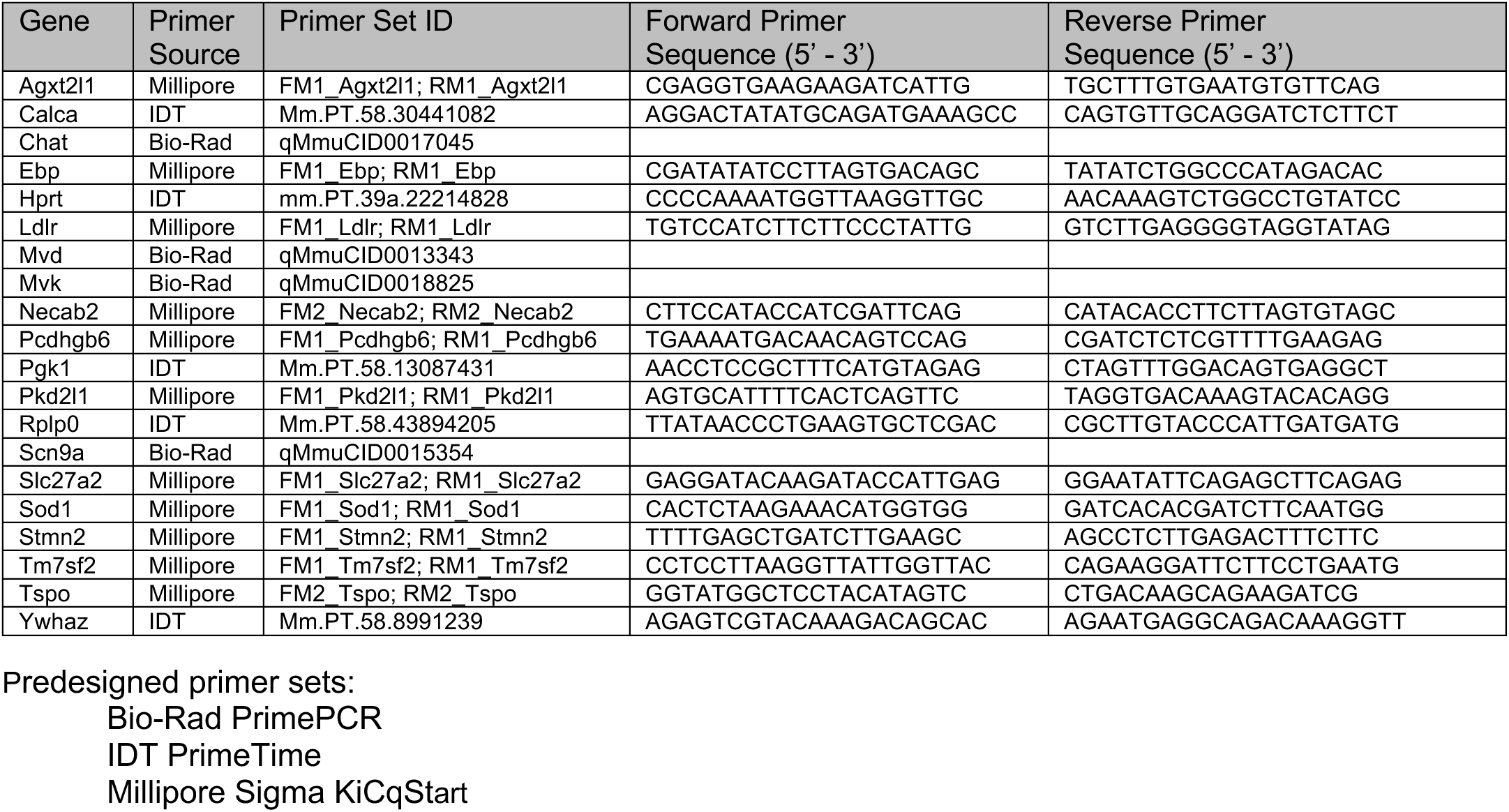
Primers Used for Quantitative RT-PCR.

